# Decoding the gene regulatory landscape through multimodal learning of protein-DNA interactions

**DOI:** 10.1101/2025.08.17.670761

**Authors:** Jimin Tan, Xi Fu, Xinyu Ling, Shentong Mo, Jiangshan Bai, Raúl Rabadán, David Fenyö, Jef D. Boeke, Aristotelis Tsirigos, Bo Xia

**Affiliations:** Gene Regulation Observatory, Broad Institute of MIT and Harvard, Cambridge, MA, USA; Institute for Systems Genetics, NYU Grossman School of Medicine, New York, NY, USA; Division of Precision Medicine, NYU Grossman School of Medicine, New York, NY, USA; Program of Mathematical Genomics, Department of System Biology, Columbia University, New York, NY, USA; Department of Biomedical Informatics, Columbia University, New York, NY, USA; Department of Pathology and Krantz Family Center for Cancer Research, Massachusetts General Hospital and Harvard Medical School, Boston, MA, USA; School of Computer Science, Carnegie Mellon University, Pittsburgh, PA, USA; Department of Biochemistry and Molecular Pharmacology, NYU Grossman School of Medicine, New York, NY, USA; Department of Biomedical Engineering, NYU Tandon School of Engineering, Brooklyn, NY, USA; Department of Pathology, NYU Grossman School of Medicine, New York, NY, USA; Society of Fellows, Harvard University, Cambridge, MA, USA

## Abstract

The identity of a cell is governed by regulatory proteins binding to the genome to control gene expression. Mapping these genome-wide binding events across thousands of proteins and cell types is essential for understanding development and disease at scale, yet has remained a major experimental and computational barrier. Here we present Chromnitron, a multimodal foundation model that learns the rules of protein-DNA binding from protein sequence, DNA sequence, and context-specific chromatin states. Unlike prior single-task and multi-task learning approaches, Chromnitron implements a multimodal learning framework that accurately predicts the binding landscape for proteins and cell types not seen during training. Using Chromnitron, we discovered and experimentally validated new protein regulators of T cell exhaustion. Chromnitron also uncovered previously uncharacterized dynamic shifts in the binding landscape of regulatory proteins during neurogenesis. This marks a critical step toward a predictive model of interpretable gene regulatory programs across cell types, enabling rapid discovery of regulatory circuits and identification of new therapeutic targets.

## INTRODUCTION

A central challenge in biology is to understand the regulatory logic that gives rise to diverse cell types and states. While the DNA sequence itself provides the basal genetic blueprint shared across all cells, this logic is executed by thousands of chromatin-associated proteins (CAPs) that interact with the genomic DNA to control cell type-specific gene expression in multicellular organisms^1–5^. These proteins – including transcription factors (TFs) that bind directly to DNA and indirectly recruited co-factors and chromatin regulators – engage thousands to millions of *cis*-regulatory elements (CREs) to orchestrate cell-type-specific gene expression programs (Fig. 1A)^6–21^. Elucidating the genome-wide landscape of protein-DNA interactions is therefore crucial for understanding the rules of gene regulation in disease and cell fate transitions^22–24^.

**Fig. 1:**
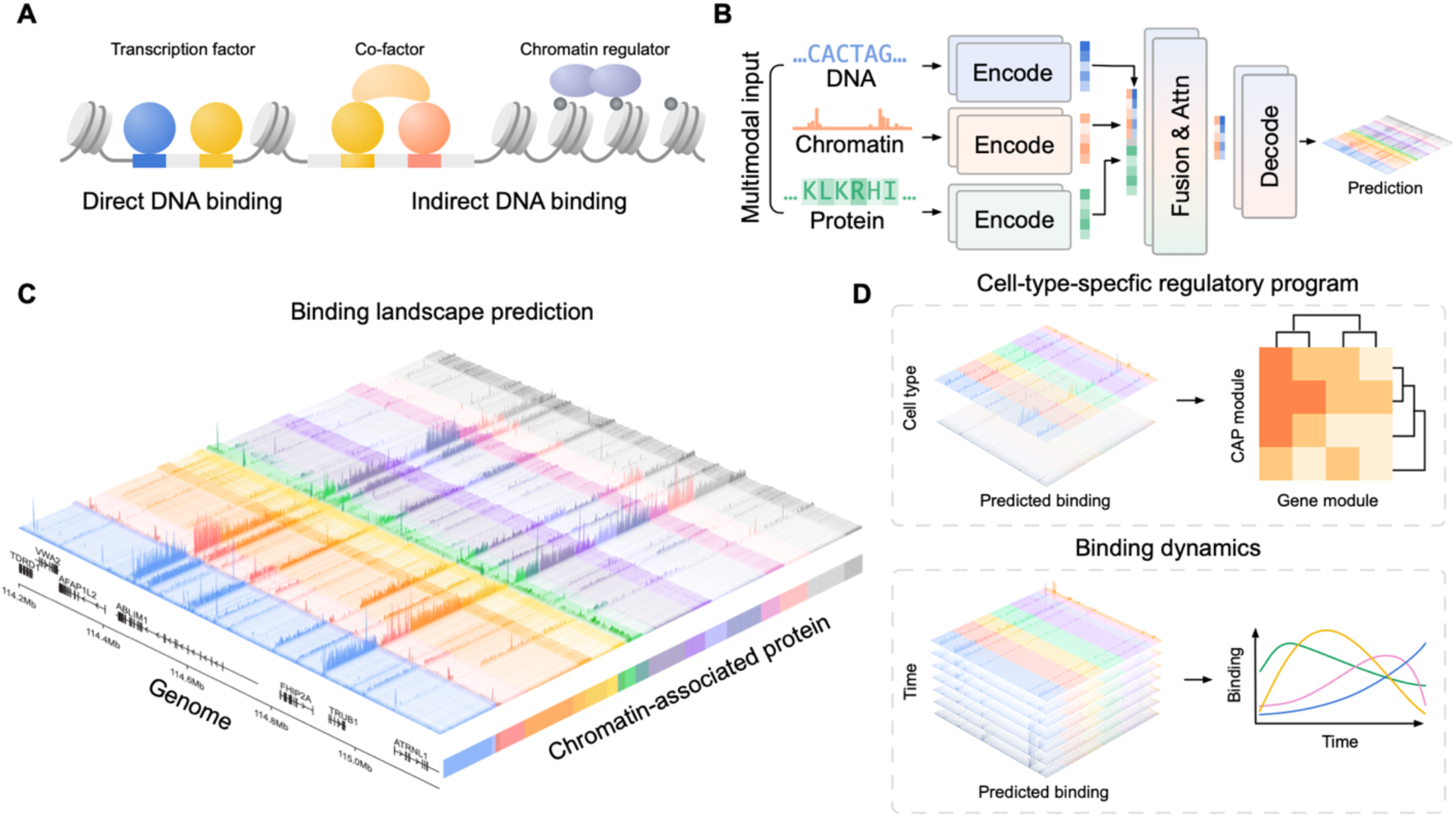
A multimodal framework for decoding the protein-DNA binding landscape. **A.** Chromatin-associated proteins (CAPs) can bind DNA directly (transcription factors) and indirectly (co-factors and chromatin regulators). **B.** A multimodal deep learning architecture that integrates DNA sequence, cell-type-specific chromatin accessibility, and protein sequence to predict CAP binding profiles. **C.** Chromnitron predicts genome-wide binding landscape for hundreds of proteins. CAPs are colored by clusters based on binding similarity. **D.** Chromnitron-predicted CAP binding landscapes enable downstream discovery applications, such as linking CAPs to CREs to identify causal regulatory factors and modules, and tracking dynamic CAP binding events across temporal trajectories.

Despite its importance, creating such a protein-DNA interaction landscape in a particular cell type has been prohibitive. Experimentally, methods such as ChIP-seq are resource intensive, measuring only one protein at a time and does not scale to primary samples due to cellular heterogeneity and scarcity in the tissue^25,26^. As a result, our knowledge of CAPs remains far from complete: genome-wide binding maps exists for fewer than half of the around 2,500 predicted human CAPs, and this data is mainly generated from cultured cell lines rather than primary cells^27–30^. Computational approaches, including recent machine learning tools that predict CAP binding from DNA sequence, have made progress but do not generalize across different cell types^31–34^. These limitations underscore the need for a predictive tool that can chart the CAP binding landscapes for any protein in any cell.

Here, we present Chromnitron (Chromatin omni-modal transformer), a biologically grounded multimodal foundation model that learns the principles of protein-DNA interaction by integrating three core modalities: DNA sequence, cell-type-specific chromatin accessibility, and protein amino acid sequences (Fig. 1B). Chromnitron was pretrained on an atlas of over 1,100 curated ChIP-seq CAP binding maps, covering 767 distinct proteins, and then fine-tuned for each CAP to capture protein-specific nuances. This cross-protein pretraining on multimodal data is key for Chromnitron to learn generalizable principles of protein-DNA interaction. Consequently, the model achieved remarkable performance in predicting CAP binding landscapes for proteins and cell types not seen during training (Fig. 1C). Chromnitron establishes a generalizable framework for decoding the global gene regulatory landscape across cell types. We demonstrate that the accurate and cell-type-specific predictions can be used to reconstruct causal regulatory programs linking CAPs to CREs, discover key regulatory factors during immune cell fate transition, and track dynamic CAP binding events along developmental trajectories (Fig. 1D).

## RESULTS

### Chromnitron learns multimodal principles of protein-DNA interactions

The specificity of CAP binding across the genome arises from protein functional domains, genomic sequence, and local chromatin context. To learn these binding principles in a single model, we designed a multimodal architecture that incorporates all three modalities as inputs (Fig. 1B and Supplementary Fig. 1). In this architecture, DNA sequence and chromatin accessibility signals were encoded by two independent convolutional modules, while CAP residues were encoded with a pretrained protein language model (ESM)^35^. The resulting embeddings are subsequently merged and processed by attention layers to allow implicit learning of both intra- and cross-modality dependencies. The final decoder module processed the chromatin embedding with a transposed convolutional module to produce CAP binding at single nucleotide resolution (Fig. 1B and Supplementary Fig. 1).

Training a high-performance foundation model requires large-scale and high-quality data^35–38^. To reduce technical variation across proteins and data sources, we first curated an atlas of 1,105 ChIP-seq datasets covering 767 unique CAPs across five different cell types (Supplementary Fig. 2, Methods). To improve consistency in data quality across CAPs and batches, we developed a fully automated and scalable preprocessing infrastructure, Chrom2Vec, which prepares raw sequencing data for training with custom denoising and signal-scale standardization across CAPs and cell types (Supplementary Fig. 2)^36,39,40^.

Next, we trained Chromnitron in two phases: cross-CAP pretraining and per-CAP fine-tuning. Our training procedure differs from traditional paradigms such as single-task learning where separate models were trained for each CAP, and multi-task learning where all CAPs were predicted simultaneously from different output heads^32,33,41,42^. The multimodal learning implemented by Chromnitron at the cross-CAP pretraining step allows the model to learn shared CAP-binding principles across proteins, establishing a single foundation model^41^. Because multimodal pretraining is computationally intensive due to its combinatorial nature, we developed an algorithm, OuterMix, that reduces the exponential sampling complexity down to linear, yielding roughly a five-fold increase in training efficiency (Supplementary Fig. 3).

Accurate and cell-type-specific prediction is essential for downstream biological discoveries. To assess whether multimodal architecture enables generalizable prediction in unseen cell-type contexts, we first trained a prototype using data from only two cell types and evaluated its *de novo* prediction performance in three leave-out cell types on leave-out chromosome 10. This prototype reliably outperformed leading unimodal frameworks, such as Enformer, Borzoi, and AlphaGenome, when predicting cell-type-specific CAP binding signals (Supplementary Fig. 4)^32,33,42^.

These results highlight the value of modeling CAP binding with multimodal learning incorporating cell-type-specific chromatin features. We therefore scaled training to all 1,105 quality-controlled ChIP-seq datasets across five cell types. During cross-CAP pretraining, we observed distinct learning dynamics across CAP groups, potentially reflecting their differential binding mechanisms (Supplementary Fig. 5). After pretraining, we performed low-rank adaptation (LoRA) fine-tuning to produce individual sub-models for each CAP, which further increased overall performance and training efficiency (Supplementary Fig. 6)^43^.

### Chromnitron accurately predicts cell-type-specific CAP binding

We systematically evaluated Chromnitron on multiple prediction tasks using chromosome 10 as a leave-out test chromosome. We found that Chromnitron accurately predicts CAP-binding profiles on par with experimental replicates (Supplementary Fig. 7). The predictions recapitulate experimental measurements across large genomic regions with varying CAP binding intensities and chromatin states (Fig. 2A, Supplementary Fig. 8-10). To assess cell-type-specific prediction performance, we benchmarked against other models on regions differentially active between liver carcinoma line HepG2 and leukemia line K562. In this context, the fully trained Chromnitron markedly outperformed both sequence-based multi-task models and other single-task models that use cell-type-specific features (Fig. 2B, Supplementary Fig. 11–13). This performance gap is particularly notable given that our evaluation was performed on a leave-out chromosome that was part of the training sets of the other models, suggesting a larger true performance improvement.

**Fig. 2:**
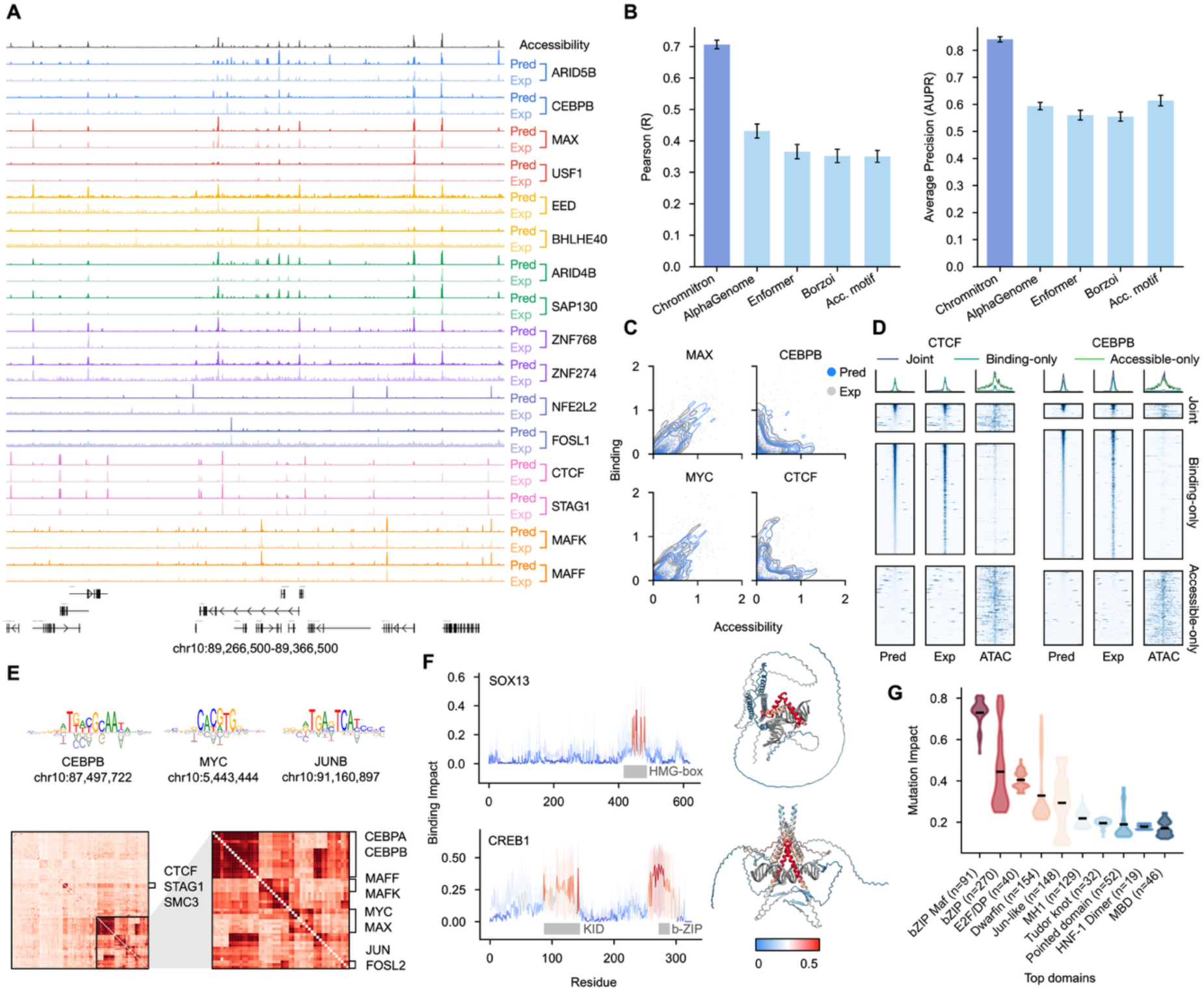
Chromnitron performance and interpretability. **A.** Paired Chromnitron prediction (Pred) and experimental (Exp) ChIP-seq tracks in leave-out chromosome. **B.** Benchmarking of prediction performance in cell-type-specific CAP-binding regions, measured with Pearson’s correlation (left) and average precision (AUPR, right). Acc. Motif, accessibility-weighted motif score. Bar plots in this figure show mean ± s.e. **C.** Scatter plots illustrating the relationship between CAP binding and chromatin accessibility for representative CAPs, and comparing experimental ChIP-seq signal (Exp, grey) with Chromnitron predictions (Pred, blue). **D.** Heatmaps comparing predicted and experimental CAP-binding signals across three classes of genomic regions: accessible with CAP-binding (Joint), inaccessible with binding (Binding-only), and accessible without binding (Accessible-only). Each row in the heatmaps represents a 2 kb genomic window centered on a peak region in leave-out chromosome. Corresponding chromatin accessibility (ATAC-seq) signals are shown in parallel. **E.** DNA sequence impact motifs learned by Chromnitron, identified through *in silico* single-nucleotide mutagenesis. Top: representative impact motifs. Bottom: heatmap and hierarchical clustering of cross-correlations among impact motifs across all pairs of CAPs. **F.** Identification of critical CAP amino acid residues via *in silico* protein residue perturbation. Left: per-residue impact scores along the protein sequences of SOX13 and CREB1. Right: corresponding impact scores mapped onto AlphaFold-predicted protein-DNA complex structures, using the same color map of binding impact scores. **G.** Distribution of per-residue impact scores across the top ten annotated protein domains, ranked by average impact score. Points in the violin plots denote individual CAPs, with the mean shown for each violin.

Notably, Chromnitron learns the non-linear, CAP-specific relationship between binding and chromatin accessibility, which is essential for generalizing across cellular contexts. Comparing experimental CAP binding signals (e.g. ChIP-seq) to chromatin accessibility, we found that many CAPs showed weak correspondence between these signals. While CAP binding is often enriched at highly accessible chromatin regions, some CAPs, such as pioneer factors, show strong binding in regions with low accessibility (Fig. 2A and 2C, Supplementary Fig. 14). Existing tools that infer CAP binding by combining motifs with accessibility peak signals are dependent on peak strength and therefore prone to false negative predictions in low-accessibility regions^44,45^. In contrast, Chromnitron accurately predicts binding signal in both high- and low-accessibility regions, owing to its ability to leverage a richer and non-linear representation of local chromatin contexts (Fig. 2D). These results highlight Chromnitron’s utility for characterizing key regulators that preferentially bind at lowly accessible regions, such as pioneer factors that drive cell fate transitions^46,47^.

### *In silico* perturbations reveal principles of protein-DNA interaction

To further interrogate the CAP binding principles learned by Chromnitron, we performed systematic *in silico* perturbation across all three input modalities. We first applied single-nucleotide mutagenesis along genomic loci and quantified the effect of each mutation on predicted CAP binding (Supplementary Fig. 15). These per-base impact scores recapitulate canonical TF consensus motifs derived from high-throughput experimental measurements^48,49^, which suggests that Chromnitron learns sequence features that determine CAP binding (Fig. 2E). Extending such *in silico* DNA mutagenesis across all CAPs revealed motif modules corresponding to distinct CAP families, along with unique ones that are previously uncharacterized (Fig. 2E and Supplementary Fig. 15). Notably, we identified coherent impact motifs for proteins lacking recognizable DNA binding domains, such as cohesin subunits, whose impact motifs resemble that of their known partner CTCF (Supplementary Fig. 15). Using the same sequence perturbation method, Chromnitron can predict cell-type-specific CAP binding changes resulting from non-coding variants in both gain-of-binding and loss-of-binding mutation events (Supplementary Fig. 16). Perturbing chromatin accessibility revealed a non-linear relationship between CAP binding and accessibility levels (Supplementary Fig. 17), with binding intensity of different CAPs saturating at varied levels. These results echoed our observation from the experimental data that many CAPs bind efficiently to inaccessible or lowly accessible chromatin regions, and support that Chromnitron learned accessibility-related features and incorporated them for prediction (Supplementary Fig. 17, Fig. 2C).

Lastly, to examine important features encoded in the CAP amino acid sequences, we applied saturated *in silico* mutagenesis to CAP amino acid sequences (Supplementary Fig. 18). This *in silico* experiment revealed that high-impact residues, which correspond to high prediction changes, align precisely with known DNA binding domains, as illustrated by SOX13 and CREB1 (Fig. 2F). Generalizing mutagenesis across CAPs revealed strong enrichments in evolutionarily conserved DNA-binding domains such as bZIP and E2F, and depletion in domains that do not contribute to DNA sequence specificity (Fig. 2G, Supplementary Fig. 18). These results suggest that Chromnitron learns protein domain features relevant for their binding specificity directly from CAP amino acid sequence. Consistent with this analysis, we found Chromnitron enables *de novo* prediction of CAP binding profiles for previously unseen CAPs. When trained on a subset of the CAPs and evaluated performance on the leave-out ones, the model can perform *de novo* predictions on unseen CAPs and outperforms the best baseline inference using motif score multiplied by accessibility (Supplementary Fig. 19). Together, these multimodal perturbation analyses show that Chromnitron captures core principles of CAP binding: (i) DNA sequence-encoded motif features and co-binding syntax; (ii) CAP-specific and nonlinear dependencies on chromatin accessibility; and (iii) protein-structural determinants of DNA recognition. These learned rules enable the model to effectively predict functional genomic data at scale and to generalize across unseen genomic loci, cell types, and CAPs.

### Chromnitron identifies ZNF865 as a *TOX* regulator during T cell exhaustion

The cell-type-specificity of Chromnitron allows the model to predict CAP binding in unseen cellular contexts. To demonstrate its utility for dissecting disease relevant gene regulation in human, we applied Chromnitron to study T cell exhaustion, an immune suppression mechanism common in cancer. T cell exhaustion is a progressive cell state transition driven by chronic antigen stimulation, leading to a dysfunctional state characterized by enhanced expression of inhibitory cell surface receptors, reduced cytokine production, and activation of *TOX*^50–57^. Single-cell ATAC-seq (scATAC-seq) of tumor-infiltrating T cells has captured the chromatin state transition from naïve to exhausted T cell but lacks mechanistic insight into the underlying regulation^58^.

To pinpoint candidate locus-specific regulators of *TOX* expression, we applied Chromnitron to track predicted CAP binding dynamics at the *TOX* locus across the T cell exhaustion trajectory derived from scATAC-seq data (Fig. 3A, Supplementary Fig. 20). Surprisingly, ZNF865, an uncharacterized C2H2 zinc-finger protein, emerged as a unique regulator at a *TOX* promoter-proximal regulatory element with a steady increase in binding along the exhaustion pseudo-time (Fig. 3B). Notably, ZNF865 show preferential binding at low-accessibility regions, including this *TOX* promoter-proximal element, whose chromatin is markedly less accessible than the promoter itself (Supplementary Fig. 20). Further *in silico* mutagenesis revealed that ZNF865 recognizes a previously uncharacterized CA-repeat sequence pattern, which explains in part its unique binding to the *TOX* promoter-proximal element (Supplementary Fig. 20).

**Fig. 3:**
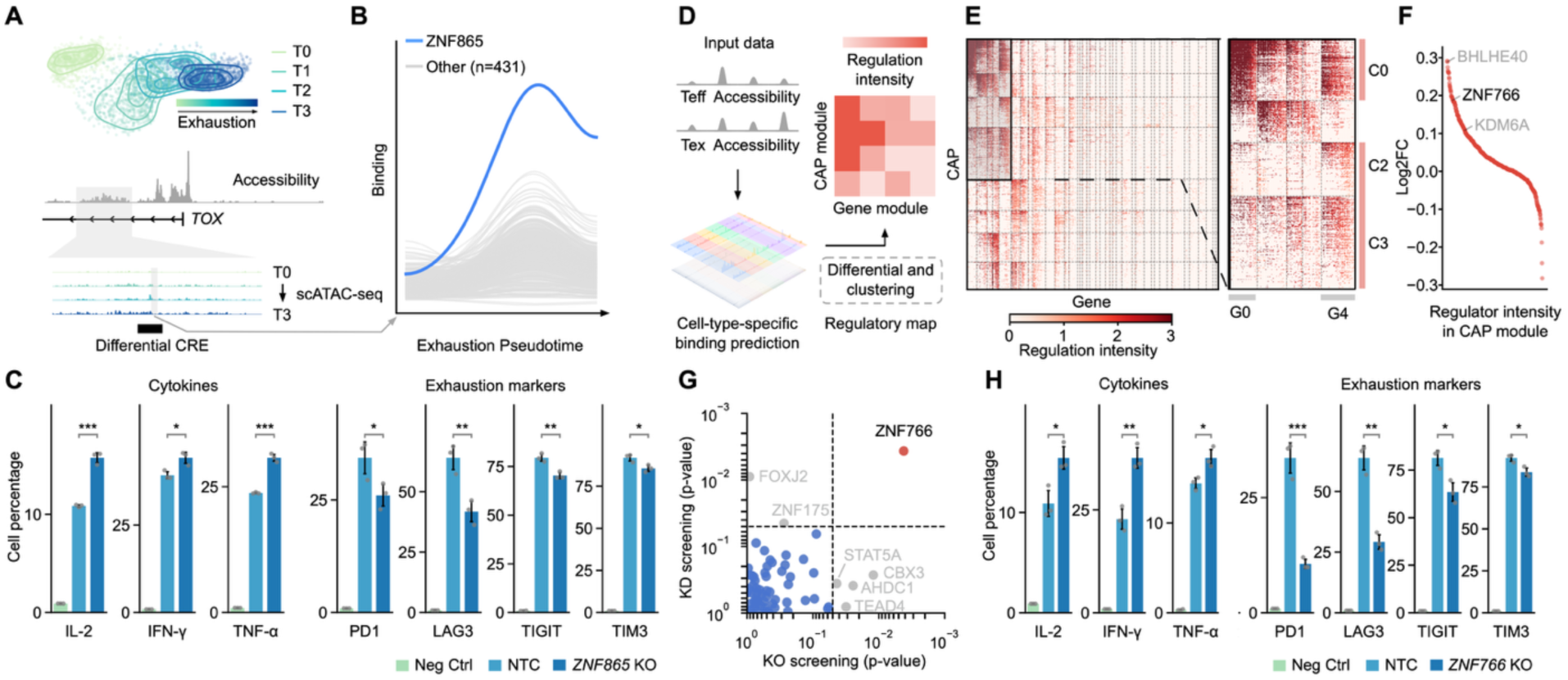
Chromnitron discovers novel regulators of T cell exhaustion. **A.** Dimensionality reduction (UMAP) of a T cell fate trajectory from naïve (T0) to exhausted (T3) states (top), with corresponding chromatin accessibility profiles highlighting a promoter-proximal regulatory element at the *TOX* locus (bottom). **B.** Chromnitron-predicted CAP-binding dynamics at the *TOX* promoter-proximal CRE across exhaustion pseudo-time, highlighting a unique and progressive increase in binding of an uncharacterized protein, ZNF865 (blue). **C.** Flow cytometry quantification of cytokines (IL-2, IFN-γ, and TNF-α) and T cell exhaustion markers (PD1, LAG3, TIGIT, and TIM3) across negative control (Neg Ctrl), non-targeting control (NTC), and *ZNF865* knockout (*ZNF865* KO) conditions. **D.** Schematic of the Chromnitron-based workflow for genome-wide characterization of co-regulation modules linking CAPs to genes by comparing effector and exhausted T cell conditions. **E.** Differential CAP-binding matrix between effector (Teff) and exhausted (Tex) T cells, clustered by both co-regulating CAPs (rows) and co-regulated gene groups (columns). A zoom-in plot highlights a subset of CAP modules (C0, C2, C3) and gene groups (G0, G4) enriching reported T cell exhaustion marker genes. **F.** Ranking of CAPs by differential binding intensity (log2 fold change between Tex over Teff), highlighting both known (grey) and unknown (blank) top candidate regulators. **G.** Scatter plot of targeting-gene sgRNA enrichment p-values from CRISPR knockout (x-axis) and knockdown (y-axis) screens, highlighting *ZNF766* as the top hit in both assays. **H.** Flow cytometry quantification of cytokines (IL-2, IFN-γ, and TNF-α) and T cell exhaustion markers (PD1, LAG3, TIGIT, and TIM3) across negative control (Neg Ctrl), non-targeting control (NTC), and *ZNF766* knockout (*ZNF766* KO) conditions. Bar plots in (**C**) and (**H**) show mean ± s.d. for *n* = 3 independent replicates. Statistical comparisons were performed using two-sided unpaired Student’s *t*-tests with significance denoted as *, P < 0.05; **, P < 0.01; ***, P < 0.001.

The timing and specificity of ZNF865 binding suggested that it may act upstream of *TOX* during T cell exhaustion. To test this, we employed an established *in vitro* T cell exhaustion model using persistent anti-CD3 antibody or anti-CD3/CD28 cocktail stimulation together with CRISPR-mediated knockout of *ZNF865*^59–61^ (Supplementary Fig. 21). Consistent with the hypothesis, knockout of *ZNF865* reduced *TOX* expression levels and consequently led to reduction in surface exhaustion markers and elevated effector cytokine production, suggesting partial restoration of the effector function (Fig. 3C, Supplementary Fig. 22).

### A genome-wide screen with Chromnitron uncovers ZNF766 as a regulator of T cell exhaustion

Having shown that Chromnitron can resolve a CAP-mediated regulatory circuit at one locus, we next asked whether it could dissect the genome-wide regulatory program driving T cell exhaustion. We predicted CAP binding across promoters whose accessibility changes between effector and exhausted T cells. Comparing the two states generated a differential gene-by-CAP matrix in which each entry represents differential binding of a CAP at a gene promoter (Fig. 3D). Clustering this matrix revealed distinct modules of co-regulated genes and CAP groups predicted to act on them (Fig. 3E). Gene modules enriched for known exhaustion marker genes aligned with CAP modules showing the strongest differential binding (C0, C2, C3; Fig. 3E, Supplementary Fig. 23). These CAP modules contained both established regulators of T cell exhaustion and several uncharacterized ones, including the KRAB zinc finger protein ZNF766 (Fig. 3F). This analysis suggested that these CAP groups may cooperate as upstream regulators of the exhaustion program.

To assess the functions of these candidate CAPs from the top regulator groups, we conducted a focused CRISPR screen targeting 65 regulators genes nominated by Chromnitron using the *in vitro* T cell exhaustion model. We designed two separate sgRNA libraries, optimized for lentivirus-based gene knockout and transcriptional knockdown, respectively. Following library transduction, human primary CD8⁺ T cells were subjected to persistent anti-CD3 treatment, with T cell proliferation serving as the phenotypic readout. Across both knockout and knockdown screens, ZNF766 consistently showed the strongest functional effect, reducing T cell persistence under chronic stimulation (Fig. 3G). Independent validation via targeted gene knockout of ZNF766 confirmed the screening results, with ZNF766-deficient T cells express significantly lower surface exhaustion markers, along with increases in effector cytokine production (Fig. 3H, Supplementary Fig. 23). RNA-seq following *ZNF766* knockout revealed broad transcriptional remodeling under chronic stimulation, consistent with the functional rescue (Supplementary Fig. 24). Together, these findings demonstrate that Chromnitron can identify genome-wide regulatory modules and uncover previously unrecognized CAPs involved in T cell exhaustion.

### Chromnitron maps the dynamic regulatory landscape during neurogenesis

To demonstrate its power in developmental contexts, we applied Chromnitron to predict the genome-wide dynamics of CAP-mediated regulation during neurogenesis using single-cell chromatin accessibility dataset (Fig. 4A–B). This yields a three-dimensional CAP-binding tensor spanning genomic position, CAP identity, and developmental time, providing a scalable framework to characterize how CAP binding and CRE activities are globally orchestrated (Fig. 4B).

**Fig. 4:**
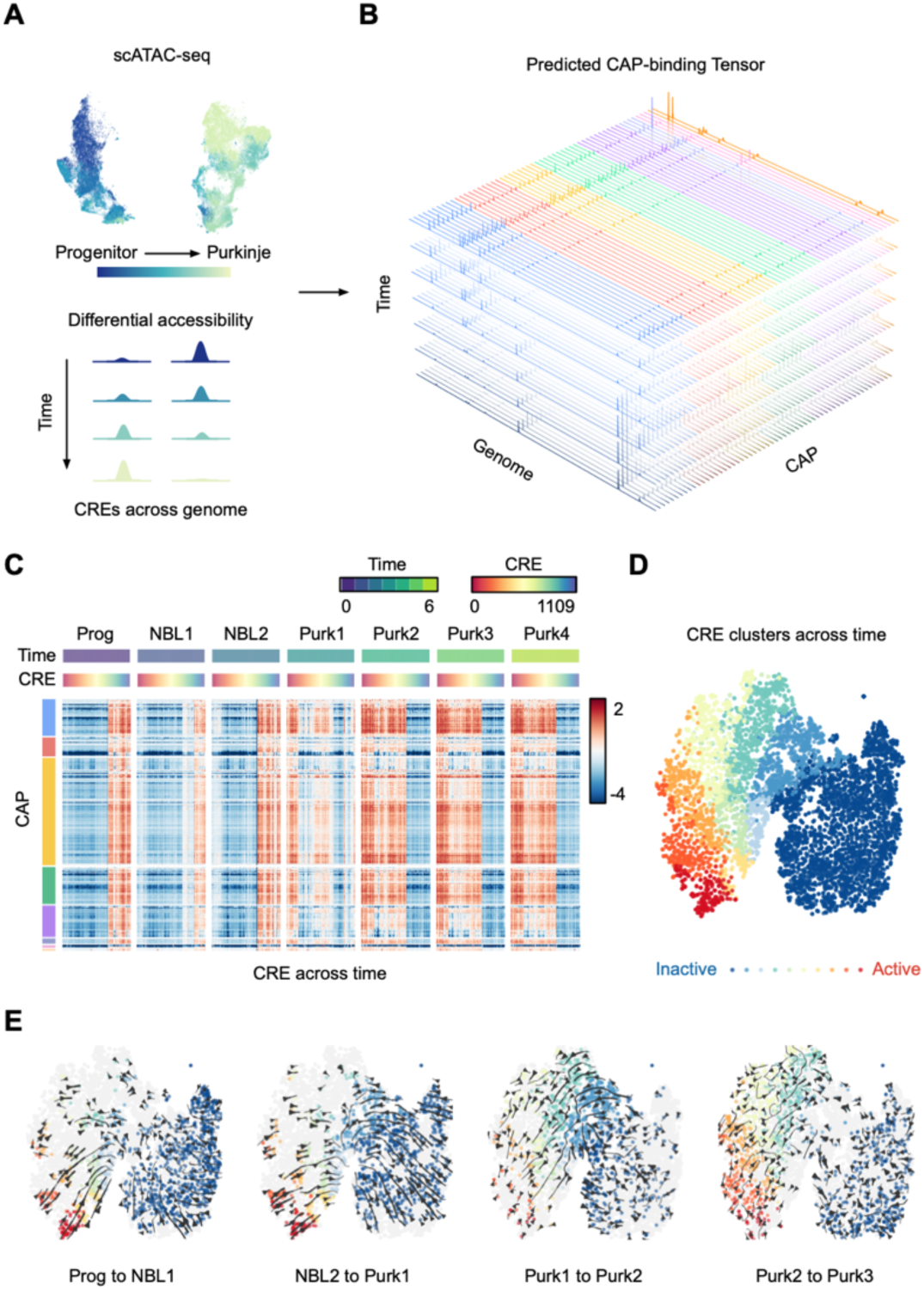
Chromnitron maps the dynamic regulatory landscape of neurogenesis. **A.** Single-cell ATAC-seq revealed differential CRE activities during Purkinje neuron development. **B.** Chromnitron predicts a three-dimensional tensor of CAP-binding landscapes spanning the genomic position, CAP identity, and developmental time. **C.** Heatmap of predicted CAP-binding signals at key CREs, with CAPs organized by binding similarity (rows) and CREs grouped by developmental stage (columns). **D.** Dimension reduction (t-SNE) projection of CREs across pseudo-time based on predicted CAP-binding profiles, revealing a continuum of regulatory element identities ranging from inactive (blue) to active (red). **E**. Vector field representation of the directional CRE state transitions between consecutive developmental stages. The vector fields were projected onto the t-SNE map in (**D**).

We applied this analytical framework to a trajectory describing Purkinje neuron maturation during first-trimester neurodevelopment^62^. When projected as CAP-by-CRE matrices grouped by developmental stage, the data revealed a clear transition in CRE activity, with a sharp shift occurring between neuroblast and early Purkinje states (Fig. 4C). To dissect stage-specific transitions in CRE activities along the developmental time trajectory, we performed dimensionality reduction on CAP features and clustered CREs. This mapping revealed a spectrum of CRE identities ranging from inactive to active states, revealing a genome-wide redistribution of regulatory element identity during maturation (Fig. 4D, Supplementary Fig. 25). Tracking individual CREs across consecutive stages generated a vector field that visualizes their “migration” through regulatory space, showing that stem-cell-specific and Purkinje-specific elements shift dynamically during early stages before stabilizing later in development (Fig. 4E–H).

Interrogating the binding tensor along its remaining axes provided additional insights into regulatory dynamics. Using developmental time as the organizing axis, we identified cohorts of early-acting and late-acting CAPs along the Purkinje cell maturation trajectory, many following monotonic gain- or loss-of-binding trends (Supplementary Fig. 26). Using the CREs as the feature dimension, we found distinct CAP binding signatures across developmental time, with cohesin-related proteins, chromatin remodelers, and repressors exhibiting more stable binding patterns across time than transcriptional activators (Supplementary Fig. 26). With its unprecedented scale of functional genomic prediction, Chromnitron enables systematic and multidimensional analyses required to elucidate CAP-mediated regulatory dynamics underlying cell-fate transitions during development.

## DISCUSSION

Understanding how CAPs configure cell-type-specific gene regulatory programs remains a major challenge in human biology. By integrating DNA sequence, chromatin state, and protein features, Chromnitron learns the multimodal determinants of protein-DNA interactions and enables highly accurate prediction of CAP-binding landscapes in unseen cell types. The model’s high predictive performance enables large-scale *in silico* experimentation, substantially narrowing the candidate search space while generating interpretable predictions that can directly guide laboratory studies. Leveraging high-throughput *de novo* prediction, we discovered previously uncharacterized regulators of T cell exhaustion and mapped global dynamics of CAP binding and CRE activities during neurodevelopment. Chromnitron demonstrates how foundation models grounded in biological principles can accelerate hypothesis generation, enable targeted experimental validation, and reveal new biological insights.

Beyond its practical applications, Chromnitron also uncovers both known CAP-chromatin binding rules and non-canonical interactions at low-accessibility or inaccessible regions with previously unrecognized sequence motifs. While sequence-based models excel at pinpointing nucleotide-level determinants of regulatory activity, Chromnitron addresses a complementary challenge: learning the fundamental rules of CAP binding through a multimodal architecture, enabling generalizable and high-performance prediction across unseen cell types and chromatin states. Chromnitron’s modular design allows the integration of pretrained DNA language models as alternative or additional encoders to further increase training efficiency and performance^32,33,35,37,42,63,64^.

In addition, this multimodal framework provides a natural foundation for incorporating CAP binding data generated from perturbation experiments. Whereas conventional single- or multi-task architectures typically require specialized model modifications to accommodate perturbed CAP profiles, Chromnitron can natively integrate protein-residue perturbation datasets and learn relationships between amino acid sequence variation and DNA binding at scale. As more diverse and perturbed CAPs are incorporated in future iterations, Chromnitron’s ability to perform *de novo* predictions for previously unseen factors can further improve.

Despite its high prediction accuracy, Chromnitron’s performance is dependent on training data quality. The current training datasets are generated primarily from conventional ChIP-seq experiments, which are subject to variable antibody quality, lack of dosage measurement, and batch effects. Although substantial computational efforts were made to denoise and standardize the data, not all processed ChIP-seq datasets reach high quality. These challenges underscore the need for generating standardized, high-quality CAP-binding datasets. Importantly, the pretrained foundation model can be readily updated by fine-tuning with improved CAP-binding datasets. Generating standardized CAP-binding datasets covering the remaining uncharacterized human CAPs will further expand the scope and power of Chromnitron’s prediction catalogue.

Predicting how a cell will behave under a given perturbation remains one of the most important and challenging problems in regulatory genomics. Like many other biological processes, cellular responses are intrinsically multifaceted, emerging from molecular interactions that span across multiple hierarchies and typically result in regulatory changes in gene expression. CAPs sit at the core of these regulatory mechanisms, directly controlling the induced changes in gene expression and cell state transitions that underlie development and disease. Chromnitron represents a substantive step toward a predictive, interpretable, and mechanistic framework for understanding gene regulation, highlighting the potential of multimodal foundation models to transform functional genomics. By accurately modeling the CAP binding landscape, Chromnitron provides a powerful tool for dissecting the mechanisms that drive cellular identity changes, building mechanistic virtual cell models, and enabling rational cell-fate engineering.

## Materials and Methods

### Model architecture

We designed Chromnitron (Chromatin Omni-modal Transformer) to integrate three information streams that jointly determine CAP-chromatin binding: DNA sequence, local chromatin accessibility, and the amino-acid sequence of the assayed CAP. Chromnitron comprises three successive components: modality-specific encoders, a multimodal self-attention module, and a decoder (Supplementary Fig. 1).

The model has three modality-specific encoders. The encoders for DNA sequence and chromatin accessibility consist of similar convolutional blocks (ConvBlocks). Each block begins with a strided Conv1D layer that covers large local windows while halving the sequence length. The convolution is followed by group normalization (groups = 1) and a Gaussian Error Linear Unit (GELU) activation function^65^. Feature maps then traverse 3 residual ConvBlocks (kernel = 9; stride = 1) that maintain length while expanding the hidden dimensionality to 384. Outputs from the DNA and accessibility encoders were projected to the same embedding of 512 × 384. The two embeddings were fused on the feature dimension to form a joint embedding of size 512 × 768. The joint embedding was scaled down to a 512 × 384 chromatin embedding with a header for subsequent fusion with the protein embedding.

For the protein encoder, we leveraged a pretrained protein language model ESM2 as a protein feature extractor^35^. As running ESM2 inference during training can add to the computational overhead, we extracted and cached the protein features before the training step to increase model training efficiency. A non-redundant list of human transcription factors was assembled by merging all proteins for which we had ChIP-seq data and the available proteins in UniProt database. To encode the embeddings, we used the publicly available ESM2 transformer with 36 layers and 3 billion parameters (esm2_t36_3B_UR50D, hidden size = 2560). For each protein, we requested the hidden representations of the final, 36th transformer block. Since ESM2 accepts up to 1,024 tokens, we split any long proteins into non-overlapping 1,024-residue segments, each embedded independently. The resulting matrices were concatenated along the sequence dimension to reconstitute the full-length protein representation. The number of residues was capped at 4,096 to accommodate most of the full-length CAP sequences while keeping a low compute cost. The per-residue embeddings were converted to arrays and stored in a compressed npz file. During training these cached embeddings are directly loaded, transformed by a header, and fused with genomic features in the multimodal attention module.

We implemented the attention mechanism in transformer to fuse contextual information both within and across modalities^66^. Chromatin feature embeddings and protein embeddings were concatenated along the length dimension, separated by a single zero vector that marks the modality boundary. Positional information was added via fixed sinusoidal encodings. The resulting fused feature embedding has a length of 769 (512 from chromatin embedding, 256 from protein embedding, and a separator with a length of 1). The fused embedding was processed by 16 layers of pre-layer-normalized transformer. Each transformer layer has 8 attention heads, 384 hidden state dimension, 2,048 feed-forward layer dimension, and a dropout rate of 0.1. Self-attention allows every genomic position to attend not only to distant nucleotides but also cross-modal to any residue of the corresponding CAP.

After transformer module, we only kept features corresponding to genomic positions and forwarded them to the decoder. The decoder has four ConvTranspose blocks successively up-sample the latent representation to the original 8,192-bp resolution while preserving contextual information through residual connections. A final convolution header produces a one-dimensional per-base binding profile for each target CAP.

Pretrained Chromnitron model has 72 million parameters. Each LoRA fine-tuned sub-model weighs 9 million parameters. The fully fine-tuned Chromnitron model with pretrained and LoRA weights has over 6.9 billion parameters in total.

### Genomic data collection

To obtain training data with more standardized quality, we primarily focused on consortium-generated datasets for consistency. We developed a systematic approach to collect and curate high-quality CAPs ChIP-seq datasets from the ENCODE database^6^. Our data collection strategy focused on obtaining non-revoked and non-archived datasets with sufficient sequencing depth to ensure reliable modeling. We required a minimum of 40 million reads to achieve adequate genome coverage for CAPs ChIP-seq and ATAC-seq, while immunoprecipitation control experiments required 100,000 reads. Chromatin accessibility information is essential for understanding the regulatory context of transcription factor binding. To ensure experimental coherence and enable integrative analysis, we retained only those CAPs ChIP-seq datasets that had good-quality ATAC-seq data in the corresponding cell type, resulting in 1,105 qualified ChIP-seq from five different cell types, covering 767 unique CAPs.

### Genomic data preprocessing with Chrom2Vec

To prepare precise and highly consistent training data, we designed Chrom2Vec, a modular bioinformatic platform that streamlines training data processing for Chromnitron. The Chrom2Vec platform starts from raw FASTQ files and delivers training-ready data tensors. Chrom2Vec is based on Apache Airflow and orchestrates next-generation-sequencing processing for both ChIP-seq and ATAC-seq. It implements configurable pipelines that handle distributed workers, tracking, and reproducible containers, performs robust assay-specific normalization, and signal compression to optimize data storage. Pipelines are described by lightweight Directed Acyclic Graphs (DAGs) that call a library of preprocessing operators, which are Airflow wrappers around containerized shell or Python scripts to ensure maximum reproducibility. While tasks are being scheduled and executed by Airflow, the metadata are streamed to Neo4j for full data-lineage tracking. To reduce storage and I/O overheads without sacrificing data quality on key genomic regions, we implemented an adaptive dynamic-bin compression algorithm. We observed as high as 50% reduction in storage for some of the high-quality tracks. The final compressed training dataset is around 952 GB in size. This architecture was optimized for machine-learning workflow as it guarantees reproducibility, supports scaling, and writes outputs in chunked Zarr format which is directly ingestible by the model and supports fast random-access during training.

To enable quantitative comparisons across cell types and experimental conditions, we performed peak-based quantile normalization on bulk ATAC-seq, ChIP-seq and pseudo-bulk single cell ATAC-seq data. We followed a two-step approach that first removed noise and then rescaled data. The noise level can be represented by signal variations in the background genomic regions. To define these regions, we used cPeaks from Meng et al., a union of accessible peaks across human cell types^67^. The background region is defined as its complementary regions on the genome, excluding ENCODE blacklist^68^.

For ChIP-seq, we performed a two-stage correction that explicitly models both biochemical background and sequencing bias. First, chromatin signal was extracted from immunoprecipitated (IP) and matched input libraries. Using the pre-defined background regions, we calculated the mean IP and input signal in the background bins. The input was rescaled to reach similar background signal intensity as IP. The rescaled input was then subtracted from the IP track, effectively removing experimental bias across the genome. To harmonize dynamic range across ChIP-seq experiments, the corrected IP signal was divided by the average of the 99 to 99.9 percentile values with 1 percent step sizes, computed over the pre-defined active peak regions from cPeaks. This normalization puts the strongest peaks near a value of one while retaining linearity at regions with lower signal intensities. The result is a bias-corrected, unit-less collection of ChIP-seq datasets that are comparable across CAPs and experiments, dramatically facilitating training stability. After scaling the training datasets, we further measured a background-noise level for each dataset by calculating standard deviation in the background chromatin regions defined above. The training data quality metric was calculated by (1 – noise) for downstream model performance evaluation.

For ATAC-seq, where a control library was not available, we removed noise from background directly followed by percentile scaling. The algorithm first estimates systematic noise by averaging the coverage signal over the same background regions. This background mean was subtracted from every genomic position, eliminating basal accessibility level. We collected mean cPeak signals, and computed the scaling constant based on top 99 to 99.9 percentiles similar to ChIP-seq. Dividing the background-corrected track by this constant compresses the dynamic range so that the signals of typical accessible loci fall between 0 and 1 (though exceptionally strong regulatory regions can exceed 1), producing normalized ATAC-seq datasets that are consistent and comparable across batches and cell types.

For single cell ATAC-seq data, we started with fragment files and processed them through a pipeline with shared preprocessing steps as Chrom2Vec and generates quantitative chromatin accessibility profiles. Fragment coordinates were standardized to the hg38 reference genome. Cells with 1,000-50,000 fragments per cell were retained to exclude low-quality cells with insufficient chromatin accessibility signal and potential doublets with artificially inflated fragment counts. Individual cells were subsequently aggregated into pseudo-bulk profiles based on cell cluster annotations (i.e. cell types or states) in original publication, ensuring each pseudo-bulk has over 10 million unique sequencing reads (or 20 million Tn5 transposase insertions). These accessibility reads from all cells within each cluster were summed to generate high-resolution chromatin accessibility tracks with improved signal-to-noise ratios compared to individual cells. Each pseudo-bulk data represents a unique cell type or state, and was normalized by the same pipeline described in the data normalization section.

### Model pretraining and fine-tuning

Training data were assembled from the three input feature modalities and the target CAP binding profiles. Each training sample unit was set to an 8,192-bp genomic region. For the two chromatin input features, we extracted a five-channel (A, C, G, T, N) one-hot representation of the hg38 DNA sequence and a single-channel chromatin-accessibility track (ATAC-seq Tn5-transposase insertion frequency signal). To stabilize training, the ATAC-seq signal was transformed with a monotonic log1p function. CAP binding profiles were retrieved from the normalized ChIP-seq coverage for each CAP in every cell type. In a separate preprocessing step, we embedded the primary amino-acid sequence of every CAP with the 36-layer ESM2 protein language model (esm2_t36_3B_UR50D). This embedding was cached and loaded on to the protein header during training.

Random samples from different modalities are recombined to form the final training batch. Batched tensors were shaped as (batch, length, channels) for DNA and ATAC, and (batch, residues, channels) for ESM2 embeddings. The model takes in these inputs and predicts the target CAP binding profile of shape (batch, length). Validation loss was calculated as the mean-squared error (MSE) between predicted and observed log1p-signal. We used Adam^69^ as the optimizer with a learning rate of 0.00002 and weight decay of 0. The scheduler used ReduceLROnPlateau strategy with a patience of 5 epochs. After validation loss stabilizes at 390 epochs, we fine-tuned the model for another 10 epochs for each CAP with low-rank adaptation (LoRA), updating the CAP-specific sub-model parameters^70^. LoRA layers were directly applied to feedforward layers in the attention. For convolution and convolution transpose layers, we custom-designed the LoRA layers, with details described in the following sections.

The model was pretrained on a node with 4 NVIDIA A100 80GB for 300 Hours with FP32 full precision. The pretraining stage took around 1,200 GPU hours and has a total compute of 8.7e19 FLOPs (floating-point operations). The fine-tuned model was trained in parallel on a cluster with 32 NVIDIA A100 GPUs using TF32 precision, which has much higher throughput. Each fine-tuning was set to run 8 hours for 10 epochs. The fine-tuning step was performed on all 767 CAPS, leading to a total compute of 3.4e21 FLOPs, and took around 6,000 GPU hours using the lower precision TF32 tensor format. The total amount of compute for training and fine-tuning is around 3.5e21 FLOPs.

Compared with previous methods that trained separate models for individual CAPs, Chromnitron training employed a cross-CAP pretraining with per-CAP fine-tuning strategy which dramatically increased both training efficiency and model performance. To measure the training efficiency difference between individualized training approach versus LoRA finetuning, we used CTCF as an example to compare the two approaches. We found that training the model from scratch took more than 30 epochs to reach good performance, whereas fine-tuning from a pretrained model only took 1 epoch to reach comparable performance. As the pretraining compute is only a small fraction of the fine-tuning compute (< 3% by estimation), assuming all CAPs requires a similar level of compute training from scratch, we estimated that the pretraining with fine-tuning strategy is at least one order of magnitude more efficient compared with individually training models for each CAP from scratch. The stored LoRA weight matrix is also 8 times smaller than full weight matrix. Meanwhile, pretraining substantially increases overall model performance through its generalized learning of CAP-binding principles across different CAPs.

### Efficient multimodal sampling with OuterMix

To improve model training efficiency at the cross-CAP pretraining phase, we developed OuterMix, an algorithm to generate Cartesian product of encoded features from randomly sampled data points on each dimension as the training units. OuterMix maximizes throughput by re-using modality-specific embeddings generated by each encoding step, thus dramatically reduces training complexities, and increases data loading and encoding efficiency exponentially.

First, a naïve sampling approach for multimodal training through randomly selecting a fixed number of examples for each batch would not be practical due to exponentially increased complexities. Each sample consists of random examples from different modalities. Given a fixed U number of samples from each modality with V modalities in total, the compute complexity would be O(*U^V^*). Formally, if we are sampling *N* ⋅ *M* ⋅ *K* samples in a batch representing the total amount of genomic loci, chromatin feature, and CAPs during pretraining, the three encoders need to encode all the samples. Using *l*, *t*, *c* to represent locus, cell type and CAP, respectively, the total random sample space for a batch *B_rand_* can be defined as:

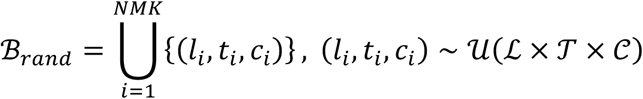

where each encoder needs to run across all samples.

OuterMix algorithm was designed to reuse embeddings generated by each encoder and reduce an exponential complexity O(*U^V^*) down to a linear O(UV) complexity, increasing data loading and encoding efficiency exponentially, especially with large encoders and a high number of modalities. With OuterMix, we started by randomly selecting fixed number of samples from each dimension and produce a Cartesian product of loci (*l_i_*), cell types (*t_j_*) and proteins (*c_k_*):

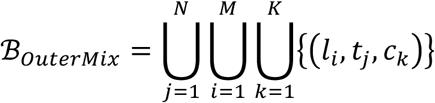

Here, each encoder only runs through samples from their predefined subset (Supplementary Fig. 3). In our implementation, the genome was first divided into non-overlapping 100kb chunks. In each iteration, the data loader selects one genomic chunk per cell type, followed by sampling 8,192 bp windows and applying lightweight augmentations. For each window, we randomly selected up to 4 CAPs with their corresponding DNA sequence, ChIP-seq, and ESM2 data. Modality-specific encoders then generate feature-specific embeddings, which are tiled and concatenated to form the Cartesian product samples ready to be fed into the Transformer. This design allows the same sequence-ATAC tensors to be paired with multiple proteins, increasing Transformer trunk throughput while avoiding additional disk I/O and encoder overheads.

### LoRA implementation during fine-tuning

Since Low-Rank Adaptation (LoRA) was originally developed for fine-tuning natural language applications such as Large Language Models (LLMs), its implementation for one-dimensional convolution and convolution-transpose layers used in Chromnitron is not trivial^70^. Here we implemented LoRA for both Conv1D and ConvTranspose1D. On a high level, LoRA for convolution layers was implemented by freezing the original kernel and learning a low-rank update that can be added onto the kernel at evaluation time. The 1-dimensional convolutional layer weights are 3D tensors representing input dimension, kernel size, and output dimension. In this implementation, we reshaped input and kernel size into one dimension and used it to construct two low-rank matrices.

For Conv1D, the convolutional kernel weights *W* ∈ ℝ*^C^*^in×^*^C^*^k×^*^C^*^out^ are kept frozen during fine-tuning, with *C*_in_, *C*_k_, and *C*_out_representing input size, number of kernels, and output size, respectively. The weight is separated into two low-rank matrices A and B with rank r, defined as:

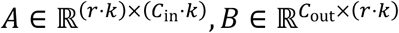

and is updated through Δ*W* = *B A*. In the forward pass, the weight is added to the frozen weight to apply fine-tuning changes:

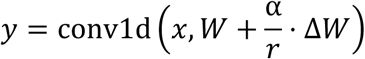

where *y* is the convolution activation and α defines the strength for LoRA weights in which the default is set to 1.

For ConvTranspose1D, the same decomposition is applied with a custom conv_transpose1d forward pass, applying the Δ*W*, similar to conv1d:

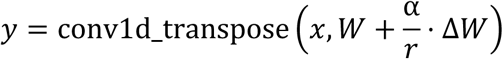

We tested a range of ranks from 4 to 16 and did not observe significant performance difference. To reduce weight size and fine-tuning efficiency, we used r = 4 for the final large-scale fine-tuning. The custom LoRA implementation is available on https://github.com/tanjimin/LoRA.

### Predicting CAP-binding profiles

During prediction, the LoRA checkpoint corresponding to each CAP was loaded together with the backbone weights. Since Chromnitron prediction operates on individual 8,192 bp windows, to predict CAP-binding signals across the entire chromosome, we used a sliding window approach with 5,120 bp step size to create overlap between adjacent predictions. Chromnitron’s predictions are robust to window shifts, but the predicted signal at the window edge can have lower quality due to the absence of flanking chromatin context. To mitigate this effect, we retained only the central 6,144 bp predicted profiles out of the 8,192 bp as valid predictions. With a 5,120 bp step size and a 6,144 effective window size, adjacent predictions will have a 1,024 bp effective overlap. To further ensure a high-quality transition between the overlapped regions between adjacent predictions, we implemented a weighted average in the overlapping regions to give more weights to bases that are closer to the center of their prediction windows. The weighted signal *S* is an average of left signal *S_l_* and right signal *S_r_*, weighted by their relative base location *i* in the region:

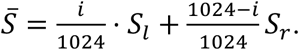

### Prediction performance evaluation

All prediction performance evaluations were conducted on the leave-out chromosome 10. To compare with experimentally measured ChIP-seq profiles, we first predicted CAP-binding profiles across all five cell types on chromosome 10. For each CAP, we extracted profiles from both prediction and ChIP-seq profiles on the chromosome 10p arm between 6 Mb and 30 Mb. We evaluated model performance by calculating a set of metrics (detailed in the following section) between prediction and experimental ChIP-seq profiles in the leave-out chromosome across all cell types. For cell-type-specific prediction performance evaluation, we first calculated the experimental data difference and the predicted CAP-binding difference between HepG2 and K562 cells, the two cell types with the highest number of overlapping CAP profiles. We then applied the same set of evaluation metrics to compare the difference signal between the two groups.

For a comprehensive comparison, we computed in total eight statistics, including Pearson’s correlation (*r*), Spearman’s correlation (*ρ*), the coefficient of determination (*R* squared), F1 score, area under the receiver-operating characteristic curve (AUROC), and area under the precision-recall curve (AUPR) with both target-centric and prediction-centric labelling. We repeated the statistics over a grid of different bin sizes and top-percentile subset of the bins. Predicted or ChIP-seq signals were first converted to numerical vectors, with missing values set to zero. Subsequently, the vectors were mean-reduced into non-overlapping bins of 50, 250, 500 or 1000 bp. This multi-scale representation allows assessment of both fine-grained and coarse regulatory patterns. The top-percentile bins measure how well the model can differentiate between peaks and their relative enrichment level. For this, we set an exponential increment of percentile using a list of 100% (1), 50% (1/2), 25% (1/4), 12.5% (1/8), …, 0.05% (1/2048). For AUROC and AUPR calculation, we used the median value of the selected bins as threshold to separate positive and negative labels. We also tested an alternative strategy to select most characteristic bins by selecting both top and bottom percentile bins (Supplementary Fig. 6). The performance metrics across methods and all bin sizes were in included in Supplementary File 2.

To categorize CAP-binding prediction performance, we focused on the Pearson’s correlation (*r*) as a simple metric to help interpret the prediction results. We categorized CAP-binding prediction performance with both validation performance and cell-type-specific performance on the leave-out chromosome (chr10) for all CAPs in cell types where training data are available. The validation performance was defined as the highest performance across all cell types, while the cell-type-specific performance was calculated as described above. To establish a relationship between validation performance and cell-type-specific performance, we correlated the validation performance against cell-type-specific performance and found a general positive correlation. The cell-type-specific performance is a relatively more challenging metric due to additional experimental noises across cell types. We selected 187 CAPs that are shared between HepG2 and K562 and fitted a line between validation performance and cell-type-specific performance. After custom evaluation of prediction performance, we assigned CAPs to different performance groups based on both validation and cell-type-specific prediction performance metrics. The specific performance cutoff was chosen based on the fitted curve between the two metrics (Supplementary Fig. 4f, and summarized Supplementary File 1). We classified a CAP into high-performance group if its prediction performance has either a validation performance > 0.6 or the corresponding cell-type-specific performance. CAPs with either a validation performance > 0.4 or the corresponding cell-type-specific performance were assigned in medium-performance group. The rest of CAPs were categorized into low-performing group. Notably, although some CAPs were classified into low-performance group by Pearson’s correlation, their predicted binding profiles recapitulate ChIP-seq peaks (Supplementary Fig. 5–6). Thus, the proposed CAP-performance categorization should be regarded as a practical metric for interpreting prediction results, rather than a strict measure of biological validity.

### Global comparison of prediction to experimental data at CAP-binding regions

To evaluate prediction performance across different chromatin backgrounds, we compared CAP-binding signals by aligning Chromnitron predictions with matched ChIP-seq and ATAC-seq profiles centered in peak regions. For each CAP of a particular cell type, three one-dimensional signal tracks were collected: (i) Chromnitron-predicted CAP-binding signal, (ii) experimental ChIP-seq data (reference), and (iii) ATAC-seq data to represent chromatin accessibility. Chromnitron-predicted CAP-binding data on chromosome 10 were generated and documented, as described in the previous performance evaluation section. Peaks were independently identified in prediction, ChIP-seq and ATAC-seq by thresholding the signal at 0.5 (half of very high peaks), merging contiguous bins above this level, and discarding events narrower than 100 bp. The mean signal within each retained interval was recorded to generate a peak set annotated by genomic coordinates and intensity.

To visualize and directly compare prediction to the reference CAP-binding data, all peaks detected in Chromnitron prediction were stratified into two groups according to chromatin accessibility signal. Each predicted peak was labelled according to its overlap with ATAC-seq peaks, generating an accessible set and an inaccessible set. For every peak in each group, we extracted a 2kb genomic window centered at the highest score in each peak. Finally, we visualized the predicted CAP-binding signal, ChIP-seq signal, and ATAC-seq signals as heatmaps to evaluate global prediction performance across chromatin background.

### Ablation study

We performed ablation studies to measure how the model performs under different constraints. This analysis includes performance evaluation across models trained without fine-tuning, and models trained with missing cell types or CAPs to test their generalizability across modalities in unseen contexts. We performed ablation studies on a random subset of chromosome 10 to reduce computation and storage consumption. Our preliminary calculation found that for one set of experiments, inferring 767 CAPs on chromosome 10 for five cell types could take two weeks and 6.5 TB of storage. Instead, we used Pearson’s correlation coefficient calculated during the validation step embedded in the evaluation process to save computation and storage. This evaluation was performed on 500 random loci on chromosome 10 across all cell types and works as a proxy of the genome-wide prediction performance statistics. The performance statistics used for both the full Chromnitron model and ablated models were all calculated from validation step using the same metrics.

To evaluate the performance improvements of fine-tuning, we extracted performance metrics from the pretrained model at its last epoch (390) and metrics from fine-tuned models at their last fine-tuning epoch (≤ 10). The last fine-tuned epoch can change due to our early stopping strategy. Then, we compared the performance individually across cell types and CAPs.

For cell-type-specific prediction evaluation through ablation study, we kept data from HepG2 and K562 cells as the training set, and left out data from IMR-90, GM12878 and HCT116 cells as testing. Chromosome 10 region was again used as leave-out chromosome. This model was then pretrained (i.e. without fine-tuning) and compared with the full Chromnitron pretraining model in leave-out cell types. For CAP-specific evaluation, we randomly left out half of the CAPs (n=382) and kept the other half (n=385) in the training set. Then we extracted the performance metrics from the ablated model and compared with the full Chromnitron pretraining model on leave-out CAPs (again in chromosome 10 regions).

### Benchmarking prediction performance

Although there was no directly comparable model that used the same multimodality framework, we benchmarked performance of Chromnitron against previous machine learning approaches that aimed for similar goals, including Enformer, Borzoi, AlphaGenome, maxATAC, and accessibility-weighted motif score as a baseline approximation^41,42^. We compared inference of CAP binding tracks from different models against the experimental data reference (Chrom2Vec processed ENCODE ChIP-seq profiles). We describe first the overall setup for different benchmarks and then the procedure for obtaining CAP-binding predictions from each model.

The *de novo* cell type performance benchmark was performed using a prototype model separately trained on HepG2 and K562 and evaluated on leave-out HCT116, IMR-90 and GM12878 cell types and at leave-out chromosome 10. The prototype model is a pretrained model and not fine-tuned for each CAP. The ATAC-seq corresponding to each cell type is used as input for prototype Chromnitron. Since sequence-based models do not accept cell-type-specific information as input and do not generalize to new cell types, the most robust estimation is computing average prediction profiles across cell types. For each CAP, we averaged prediction profiles across all cell lines except for the cell type that is under evaluation. Since most of the region on the genome have rather conserved binding across cell types, we selected cell-type-specific regions by performing pair-wise differential analysis for 3 pairs between HCT116, IMR-90, and GM12878. The genome is binned at 1 kb. The top 1% most differential bins based on signal log2 fold change was used as regions for evaluation. We note that all selected regions are on Chromnitron leave-out chromosome 10 for stringent evaluation, but many of which are in the training set of other sequence-based models due to different data splitting strategies. We compared performance of prototype Chromnitron against aggregated prediction from sequence-based models and accessibility-weighted motif baseline on these cell-type-specific regions. Since each model has different numbers of prediction target, we selected the joint CAP subset shared between Chromnitron, AlphaGenome, Borzoi and Enformer for benchmarking purpose.

The overall evaluation with the full Chromnitron model is performed with a similarly setup on HepG2 and K562. The model used in this evaluation is pretrained on all cell types and fine-tuned for each CAP as described in the model training method section. The same prediction aggregation is performed for all sequence-based models. The same region selection strategy is used to obtain differential regions between HepG2 and K562 on Chromnitron leave-out chromosome 10. The chromosome-wide evaluation has the same model prediction setup but is performed on the entire chromosome 10.

Enformer: We used GRCh38 reference genome sequence as input to Enformer. Chromosome 10 was divided into non-overlapping 100 kb windows; for each window, 196,608 bp centered on the window mid-point were retrieved (zero-padded at chromosome ends) and one-hot encoded. Each sequence was forwarded through the PyTorch Enformer implementation (EleutherAI/enformer-official-rough) which has similar performance as the TensorFlow implementation. The model returned an 896-bin output tensor per window which was cached on disk. CHIP channels relevant to HepG2 or K562 cells were identified from human ChIP-seq targets. For every requested TF channel, the central 50,176 bp (±50,176 bp) of each window were extracted to avoid model padding artefacts. The selections were written into an array representing chromosome 10 at 1 bp resolution. Overlaps occurred only at window boundaries and were overwritten by later windows. The final arrays were exported to BigWig files for visualization and comparison with other approaches.

Borzoi: We used the same reference sequence and processed chromosome 10 in non-overlapping 523,264 bp windows. For each window, a 524,288 bp sequence centered on the window midpoint was retrieved (zero-padded at chromosome ends if needed) and one-hot encoded via the Baskerville DNA utilities. The sequence was fed to a single-replicate Borzoi model (SeqNN) initialized from the published parameters/targets (params_pred.json, targets_human.txt) and restored fold0 weights (model0_best.h5). The model returned per-track predictions for the full window; raw output tensors were cached for downstream analysis. Processing advanced across chromosome 10 in 523,264 bp steps; trailing partial windows were skipped to avoid padding artefacts. Optional post-processing (not run in the main loop) converts selected tracks to dense 1 bp resolution arrays and exports BigWig files for visualization and downstream comparison.

AlphaGenome: We tiled chromosome 10 in non-overlapping 1,048,576 bp windows and skipped trailing partial windows. For each window, we called the AlphaGenome DNA API (dna_client.create) with organism HOMO_SAPIENS, OutputType.CHIP_TF only, and no ontology terms. We requested predictions via API over the exact interval without padding. The prediction server returned a CHIP-TF tensor (output.chip_tf.values) plus accompanying track metadata. We cached the raw prediction arrays and metadata CSVs. The resulting prediction arrays are merged and converted to individual bigwig files for further analysis.

MaxATAC: Raw ATAC-seq data were obtained from ENCODE as BAM files (HepG2: ENCFF939JOT, K562: ENCFF804MXV). These files were down-sampled to 40M reads and deduplicated to generate the BAM files required for maxATAC pipeline. MaxATAC pre-processing (“maxatac prepare”) was performed with “-c chr10” to build the ATAC coverage tracks necessary its prediction. For every CAP that is included in both Chromnitron and maxATAC model, we used the corresponding sub-model to predict the CAP binding profile as a BigWig file using “maxatac predict”.

Accessibility-weighted Motif (Acc. Motif) score: Non-redundant motif from Jeff Vierstra (https://resources.altius.org/~jvierstra/projects/motif-clustering/releases/v1.0/) were filtered to only retain human CAPs. Motif scanning results on chr10 were converted into a dense 1-bp signal for each motif with the corresponding matching scores. A 20-bp moving average was applied to the dense track. The final results were stored as BigWig files. To weight motif by chromatin accessibility, we loaded ATAC-seq for all five cell types (HepG2, K562, GM12878, IMR-90, HCT116). For each cell type, the smoothed motif track was multiplied elementwise by the corresponding ATAC-seq signal and stored as BigWig files.

### *In silico* DNA sequence mutagenesis

To identify the impact of DNA sequence on the binding of a CAP, we focused on regions where Chromnitron predicted intense CAP-binding signals. For each CAP-binding peak, we extended the locus by ±3 kb and records its midpoint base location. We defined a 300-bp “mutation window” (±150 bp) flanking the midpoint and systematically substituted every base within the window. Subsequently, we performed *in silico* saturated DNA base mutagenesis of the 300 bp window, one base at a time. For every position, we generated four sequences, each carrying a single A, C, G or T substitution, and predicted the corresponding CAP-binding signal for each sequence. We recorded the original sequence and its predicted CAP-binding signal as wild-type (WT) reference. We then calculated the effect of every mutation as the difference between mutant and WT predictions. The resulting inferred CAP-binding data at this window was stored as a 300 × 4 × 8,192 tensor, with the three dimensions representing the number of mutation positions, number of substitution nucleotides, and the window size of the CAP-binding prediction, respectively. The signals represent the predicted CAP-binding intensities. To represent only the CAP-binding region, the tensor was cropped from 8,192 bp to the central 1 kb and averaged, generating a 300 × 4 matrix whose values represent CAP-binding intensities across the 300 bp mutated positions and 4 nucleotides. This matrix served as the nucleotide-resolution perturbation signals for downstream motif analyses and interpretation. For visualization purposes, the impact motif was normalized by subtracting the median CAP-binding intensities across the 4 nucleotides (i.e. the mean binding intensities of the middle two nucleotides) at each position. In essence, the impact motif is computed from a single locus, differing from the conventional motif calling which computes consensus sequence from a large number of loci.

### Cross-CAP motif discovery and clustering

To analyze motifs across all CAPs, we generated weight matrices (represented by median-subtracted CAP-binding signals as described above) for all CAPs at their top binding loci. These weight matrices were used as input for TF-MoDISco to generate motifs for each individual CAP^71^. For every CAP, the tool scans up to 20,000 high-scoring 200-bp windows (“seqlets”, or high-importance windows of the sequences, as referred in TF-MoDISco), clustering them by k-mer similarity and reports positive and negative patterns (i.e. seqlets). After identifying relevant seqlets for a CAP, we mapped their corresponding seqlets back to the genomic coordinates and applies a strict quality control procedure: a pattern must contain at least one seqlet whose high (90th-percentile) absolute weight difference is nontrivial (>0.1) after median subtraction. This procedure excluded noisy results from the analyses. We then stored the remaining patterns for every CAP. To pick the most representative seqlets for each CAP, we ranked them by (1) the number of retained seqlets, (2) the overall motif intensity represented by the median difference value and (3) the highest motif intensity represented by median 90th percentile difference. Patterns whose seqlets overlapped more than ten distinct loci and repeated in different CAPs were discarded to avoid trivial repeats.

Then, we built normalized weight matrices for cross-CAP comparison from the top ranked seqlets. For each seqlet, we subtracted its per-position median and scaling by its mean absolute difference. Each motif was flattened into a single vector (length by 4). Pairwise distances were computed as the maximal cross-correlation between motif A and motif B or B’s reverse complement. We performed hierarchical clustering with Ward linkage based on the distance metrics, generating the clustering heatmap for different motifs.

### *In silico* chromatin accessibility perturbations

We measured the changes of predicted CAP binding upon *in silico* scaling of chromatin accessibility signal to examine Chromnitron’s sensitivity to local chromatin states. We employed a scaling approach through multiplying the ATAC-seq profiles by a defined factor (multiplier) to alter accessibility intensities while preserving peak shape. Meanwhile, the size of the perturbation window (i.e. number of bases) also matters in this *in silico* perturbation. Thus, we created an *in silico* perturbation grid by combining multiplicative factor between 0 to 2, and with a perturbation radius that ranged between 50 and 1,000 bp. For a given radius, the slice of the ATAC vector centered on the locus was multiplied by the selected factor; the remainder of the signal was kept the same. For each combination of multiplier and perturbation radius size values, we predicted the corresponding CAP-binding signal for perturbation and calculated the binding differential by subtracting the unperturbed condition, generating a data tensor of 40 × 20 × 8,192 (multipliers × window-sizes × predicted profile). Similar to *in silico* DNA sequence mutagenesis, we averaged the center 400 bp region of difference between perturbed prediction and wild-type prediction to generate a 40 × 20 matrix that measures the accessibility perturbation effects.

### *In silico* protein residue mutagenesis

To investigate how the model leverages protein embedding features in prediction, we performed systematic mutagenesis directly on the input CAP residues without referring to particular protein domain or functional data. Each perturbation operates on a 30-amino acid sliding window with single-residue steps, replacing the window with random amino acids to generate the corresponding mutant protein sequence. Mutated CAP sequence was feed into the same protein encoder (ESM2) to generate the corresponding mutant embeddings, which were used in downstream prediction of mutant binding profile and comparison with wild-type protein.

To ensure more robust comparison of protein mutation effects between the mutant and wild-type CAP, we selected the top 200 Chromnitron-predicted binding peaks for each CAP in HepG2 cells, filtered for ATAC-seq overlap, and equally extended to the flanking sequences up to 8,192 bp to facilitate prediction. Chromnitron predictions of the corresponding CAP were generated for wild-type and mutant embeddings across all selected loci. We then extracted maximum scores from the center 400 bp region of each window (recorded as “wildtype_max” and “mutant_max”, respectively). CAP mutation impact scores were calculated as 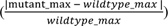. For protein domain interpretation, we retrieved InterPro domain annotations and computed precision-recall curves using domain boundaries as labels and mutation scores as predictions^72^. Domain enrichment was quantified using AUPR compared to random baselines.

### Inferring CAP interactions through colocalization analysis

To investigate whether model prediction serves as a potential source to study protein-protein interaction between CAPs, we followed our previous approach and applied linear non-Gaussian acyclic model (LiNGAM) causal-discovery algorithm to a selection of high-quality CAPs prediction across HepG2 ATAC-seq peaks^38,73^. We found that, as expected, the strongest colocalizations were often between transcription factors that belong to the same family as they usually share similar conserved DNA-binding domains. To systematically evaluate whether colocalization calculated from experimental ChIP-seq or predicted CAP-binding data reconciles with known physical protein-protein interactions, we benchmarked our causal discovery-based interaction graph with Pearson’s correlation-based interaction graph. This analysis was performed on both experimental and predicted ChIP-seq data using confident interaction annotation from STRING database (using scores over 700 as positive, and 400 to 700 as negative)^74^. The causal discovery on predicted data yields the highest AUPR (0.426), followed by causal discovery on observed data (0.372), while the correlation-based approaches achieved AUPR scores of 0.351 and 0.321 for predicted and experimental ChIP-seq data, respectively (Supplementary Fig. 11). The lower performance when using ChIP-seq data could be a result from heterogeneous noise distribution in the experimental data compared to the predicted results, leading to prediction of only strongest interactions.

### Dosage correction of predicted CAP binding

Here we describe the post-inference dosage-correction module to adjust predicted CAP binding by the inferred availability of the corresponding CAP, using either CAP promoter accessibility or CAP RNA abundance as a proxy for dosage.

For each CAP and cell type we obtain (i) promoter/TSS chromatin accessibility (log-transformed read depth or signal at the TF’s annotated TSS window) or (ii) bulk or pseudo-bulk RNA-seq expression (log-transformed TPM/CPM or normalized counts). These scalars were matched to the CAP predictions for the same cell type. These scalars can be used to either scale the prediction or gate it. When used as a dosage multiplier, we multiply the log-transformed scalar to the corresponding predicted CAP binding profile. When used as gating, we define a threshold representing zero dosage of the CAP. Thresholds can be chosen empirically from the distribution of accessibility or expression across TFs to balance false suppression of true binders against removal of non-expressed CAPs. Predicted binding profile of CAPs that fall below the threshold are set to zero.

Dosage correction is applied post inference and does not affect prediction. It can be toggled between the scaling and gating modes depending on whether a graded or binary adjustment to TF availability is desired. This module was not used in the cell-type-specific analyses to avoid potential bias and turned off by default.

### Single-cell differential ATAC-seq peak calling

To identify differentially accessible chromatin regions between pairs of cell types from single-cell ATAC-seq data, we employed a computational workflow utilizing SCANPY^75^. For each pairwise comparison, we first extracted the cells from the two specified populations the from complete peak-by-cell matrix. Peaks with low signal, defined as those with fewer than 500 total fragment counts across the selected cells, were removed from consideration. To account for technical variations in sequencing depth, we normalized the counts for each cell to a total of 100,000, followed by a log1p transformation to stabilize the variance. Subsequently, we identified highly variable peaks between the two cell populations by using a dispersion-based method implemented in SCANPY (scanpy.pp.highly_variable_genes), retaining peaks that exhibited a normalized dispersion of at least 0.1. The genomic coordinates of these candidate regions were used as candidate differentially accessible regions and exported for downstream analysis.

### Predicting CAP-binding at single locus across T cell states

We used a public single-cell ATAC-seq data of T cell exhaustion collected from human patient PBMC samples^58^. Four representative T cell stages (from effector to exhausted states) were prepared to relate CAP binding changes to T cell states. Pseudo-bulk ATAC-seq data for each stage was generated by sampling 10M reads from cells in the vicinity of selected center. The pseudo-bulk ATAC-seq was then processed and normalized as described in the single cell data processing method section. Differential accessible peaks of the corresponding cell populations were called as described in the above section. These peak regions served as putative *cis-*regulatory elements that might play important functions in T cell states. Next, for a given regulatory element of interest, we predicted CAP-binding profiles at an element-centered 8,192 bp window across the cell states. As single locus is highly susceptible to noise, we selected a set of inference CAPs using a more stringent cutoff of 0.5 (Pearson’s *r*) validation performance for this analysis. Predictions for every T cell state were cropped to a 300 bp window around the CRE center. The binding signal was normalized by median on the genome dimension to remove global scaling differences. Trajectories were plotted as ridge plots that overlay the complete track profile per CAP. The mean CAP-binding signals across trajectory were smoothed with Gaussian kernel and visualized with line charts tracing binding dynamics across cell state transition trajectory.

### Genome-wide identification of regulatory CAPs in T cell exhaustion

The same T cell exhaustion scATAC-seq and annotation in single-locus-analysis were used for this analysis. We first isolated reference pseudo-bulks for both effector T cells and exhausted T cells according to the original cell type annotations^58^. To obtain a cleaner causal relationship between CAPs and target genes, we focused on promoter regions as the putative regulatory elements. We used GENCODE v45 transcript annotation (GFF3) and the hg38.chrom.sizes file hosted by IGV to define promoter coordinates and chromosome boundaries^76^. Promoter regions (± 500 bp around the transcription-start site, TSS) were extracted from the GENCODE annotation. Genes on sex chromosomes or mitochondrial DNA were discarded. We also removed transcripts not matching a whitelist of supported biotypes (protein-coding, lincRNA, miRNA, snRNA, snoRNA, rRNA). The maximal and mean ATAC-seq accessibility signal were computed in both T cell states for each promoter. Promoters with a cross-condition maximum accessibility ≤ 1 were removed to exclude inactive loci. We then calculated log2 fold change (log2FC) of accessibility for all the loci, and partitioned them into three categories: (1) up-regulated promoters, defined as top 2,000 positive log2FC values, (2) down-regulated promoters defined as top 2,000 negative log2FC values, and (3) stable promoters, defined as 100 promoters with the smallest absolute log2FC. We also considered alternative TSS as separate promoters and implemented additional constraints to merge overlapping TSS loci.

Due to the high compute requirement for this task, Chromnitron prediction was sharded by cell type and all CAPs, and executed in parallel on a GPU cluster. For this task, we used the full set of CAPs since genome-wide analysis can better tolerate noisy predictions. After all shards completed, we merged the predictions by iterating over chromosomes, stacking the individual CAP arrays to generate genome_length × CAPs matrices. This matrix was rechunked to optimize downstream biological analyses for characterizing putative regulator CAPs.

Promoter-level average CAP-binding signals were extracted by recording the maximum predicted signal in the 1 kb window centered at TSS, generating a three-dimensional tensor with dimension (cell types × promoters × CAPs). Downstream analyses used the log-transformed values. A promoter-specific differential CAP-binding score was calculated by subtracting the predicted CAP-binding signals of the exhaustion group by that of the effector T cell group. To account for basal CAP-specific variability, the difference matrix was z-standardized by subtracting the mean and dividing by the standard deviation of the difference values measured at stable promoters. To increase signal-to-noise ratio, mid-range values between the 10th and 90th percentiles were set to zeros separately across loci and CAPs, thereby highlighting the most differential – potentially functional – signals for informing downstream analysis. Next, we performed k-means clustering on both the loci and CAP dimensions. Silhouette analysis over k-means clustering using k ranging from 3 to 40 was used to derive optimal number of clusters. To identify important CAPs and loci within each cluster, we ranked loci and CAPs by revaluating the average binding against the opposing cluster assignments iteratively on both dimension until convergence.

### Global CAP and CRE dynamics characterization during Purkinje cell differentiation

Single-cell ATAC-seq datasets for Purkinje cell development were obtained from Mannens et al^62^. We adopted the original assignment of Purkinje cell differentiation trajectory encompassing seven stages: progenitor cells (Prog), two neuroblast stages (NBL1, NBL2), and four Purkinje cell maturation stages (Purk1 to Purk4). All datasets were processed using the scATAC-seq preprocessing steps described earlier. Only autosomes were included for this analysis.

To identity genomic regions of interest, we used multiple filtering approaches based on chromatin accessibility dynamics across developmental stages. Initial candidate regions were generated from cPeaks representing the repertoire of accessible regions across human cell types^67^. We extended each peak center by 500 bp flanking region on each side to create 1 kb inference windows to include the complete peak region. Then, we filtered these loci by classifying them into two categories based on temporal accessibility patterns: differential and stable regions. Differential loci were defined as regions exhibiting significant accessibility changes across developmental stages (log2 fold-change larger than 0.1 compared with average accessibility), representing dynamic regulatory elements. Stable loci have relatively minimal accessibility change, defined as having absolute log2FC smaller than 0.1. Similar to the genome-wide regulator screening in T cell exhaustion, these stable loci served as normalization controls for downstream analysis. We selected the top 1,000 differential loci and top 100 most stable loci. To ensure non-redundant genomic coverage, overlapping regions were merged. Regions with insufficient chromatin accessibility across all conditions were excluded to prevent analysis of inactive or noisy genomic elements. The final dataset comprised 1,073 high-confidence genomic loci distributed across autosomes.

This task was computationally intensive and was thus parallelized on a GPU cluster. To reduce compute, we only keep a set of high-performance CAPs similar to the single locus T cell exhaustion inference experiment. Individual CAP predictions were concatenated to construct a 3D data tensor of developmental time × genomic loci × CAPs. For each genomic locus, we calculated the maximum peak heights within 1 kb windows centered at peak summits. The resulting 3D tensor was filtered for missing values, infinite numbers, and outliers. Data normalization was performed using stable loci as internal controls to account for technical variation and global accessibility changes across developmental stages. We performed z-score normalization using the mean and standard deviation of differential patterns calculated from stable loci. Outlier detection and removal were implemented using percentile-based clipping (1st and 99th percentiles) to prevent extreme values from dominating downstream clustering analyses.

Regulatory pattern discovery was performed using multiple complementary approaches to capture different aspects of the temporal regulatory landscape. The 3D tensor was reshaped into matrices emphasizing different analytical perspectives. There are three ways to transform the 3D tensor to 2D matrix for specific analyses, each enabling discovery of distinct regulatory relationships. First, we can use CAP as features, reshaping the tensor to a matrix with dimension (time × loci) by CAPs to identify loci whose activities change across time. Second, we can use loci as features to generate the (time × CAPs) by loci matrix. This approach can represent how CAP binding profiles change across time. Finally, using time as the feature, the reshaped matrix can tell how a binding event specific to a CAP and a locus varies across time.

We focused on the first analysis using CAP as features. To observe CRE activity shifts across time, we performed hierarchical clustering of CREs on the first pseudo-time point. This approach grouped CREs with similar regulatory dynamics. To further elucidate the relationship between CRE and time, we applied k-means clustering using a k between 5 and 20 to identify regulatory locus modules across all time points and CREs. The optimal cluster number was determined using silhouette score. Clustering was performed on unit-normalized data using Euclidean distance metrics, with multiple random initializations to ensure stable cluster assignments. For visualization and interpretation, t-distributed Stochastic Neighbor Embedding (t-SNE) was used to generate 2D embeddings of high-dimensional regulatory patterns. Each dot on the t-SNE embedding represents a CRE at a specific time point and was colored by clustering results representing its identity.

To visualize transition of CREs between adjacent pseudo-time points in Purkinje cell development, we calculated a vector field where the start and end were defined by the t-SNE location of each locus at first and second time points. This velocity field represents how CREs change their regulatory identity between consecutive developmental stages. We applied Gaussian kernel smoothing to the vector field to reduce noise while preserving temporal trends.

### Cell culture and primary cell isolation

HEK293T cells were obtained from the American Type Culture Collection (ATCC). HEK293T cells were maintained in Dulbecco’s Modified Eagle Medium (DMEM; Gibco, 11965092) supplemented with 10% fetal bovine serum (FBS; Gibco, A5670701) and 1% penicillin–streptomycin (Pen/Strep; Gibco, 15140122). Human Peripheral blood mononuclear cells (PBMCs) were isolated from healthy donor leukopaks (Charles River Laboratories Cell Solutions, 103051-034) using Ficoll-Paque PLUS density gradient centrifugation (Cytiva, 17144002) according to the manufacturer’s protocol. Isolated PBMCs were cryopreserved in 90% human AB serum (MP Biomedicals, 092938249) plus 10% dimethyl sulfoxide (Sigma-Aldrich, D4540) at a density of 30M cells per ml. For in vitro T cell assays, thawed PBMCs or purified CD8⁺ T cells (isolated using EasySep™ Human CD8+ T Cell Isolation Kit, STEMCELL Technologies, 17953) were cultured in X-VIVO 20 medium (Lonza, BW04-448Q) supplemented with 5% heat-inactivated human AB serum (MP Biomedicals, 092938249) and 10 ng/ml recombinant human IL-2 (PeproTech, 200-02-50UG). T cell cultures were maintained at 37 °C in a humidified incubator with 5% CO₂.

### sgRNA library preparation

For each selected gene, five sgRNAs were designed to either knock out or repress gene expression using the CRISPick online tool (Broad Institute). The complete list of sgRNA sequences is provided in Supplementary File 3. A total of 370 DNA oligonucleotides corresponding to sgRNAs were synthesized as pooled libraries (500 ng per library) by Twist Bioscience. For gene knockout experiments, sgRNAs were cloned into the lentiviral vector pXPR_023 vector (Broad Institute, also available through Addgene #52961), which encodes Cas9, a puromycin resistance gene, and the sgRNA scaffold. For gene repression (CRISPRi), sgRNAs were cloned into pXPR_066 vector (obtained via Broad Institute Genetic Perturbation Platform), which expresses KRAB-dCas9 under the EFS promoter along with a puromycin resistance cassette and sgRNA scaffold. Oligonucleotides were cloned into the respective backbone vectors using Golden Gate assembly with BsmBI restriction enzyme (New England Biolabs, R0739S), following the protocol described by the Feng Zhang lab (Broad Institute). Ligation products were transformed into NEB Stable Competent *E. coli* (New England Biolabs, C3040H) and plated on LB-agar plates supplemented with carbenicillin (100 μg/ml). Colonies were scraped from the plates and pooled, and plasmid DNA was extracted using the ZymoPURE II Midiprep Kit (Zymo Research, D4201) and quantified by NanoDrop for downstream lentivirus production.

### Lentivirus production

Lentivirus was produced in HEK293T cells (ATCC; passage number <20) seeded at 4 × 10⁷ cells per 150-mm tissue culture dish. Twenty-four hours after seeding, the culture medium was replaced with fresh pre-warmed DMEM containing 10% FBS. Transfection was performed using polyethyleneimine (PEI 40K; Polysciences, 24765) as follows: 13.3 μg of the sgRNA vector, 10 μg of packaging plasmid psPAX2, and 6.7 μg of envelope plasmid pMD2.G were mixed in 400 μl of Opti-MEM (Thermo Fisher Scientific, 31985070) (Tube A). Separately, 90 μl of PEI (1 mg/ml) was diluted in 310 μl of Opti-MEM (Tube B). The two solutions were incubated separately for 5 min at room temperature, combined, and incubated for an additional 15 min before being added dropwise to the HEK293T cells. Seventy-two hours post-transfection, the culture supernatant was harvested and centrifuged at 3,000g for 15 min to remove cellular debris. Lentiviral particles were precipitated by adding polyethylene glycol 8000 (PEG8000; Sigma-Aldrich, P2139-500G) to a final concentration of 8% (w/v) and incubating the mixture overnight at 4 °C. The precipitated lentiviral particles were pelleted by centrifugation at 1,500g for 30 min and resuspended in 500 μl of X-VIVO 20 medium (Lonza, BW04-448Q). Aliquots were stored at −80 °C until use.

### T cell stimulation assays

Cryopreserved human CD8+ T cells were thawed and resuspended in T cell culture medium (X-VIVO 20 supplemented with 5% human AB serum and 10 ng ml⁻¹ recombinant human IL-2) at a density of 1 × 10⁶ cells per ml in 12-well plates. Cells were stimulated by the addition of Human CD3/CD28 T cell Activation Beads (BioLegend, 422603) at a 1:1 bead-to-cell ratio and incubated for 48 h at 37 °C with 5% CO₂. Following activation, beads were removed using a magnetic separation rack according to the manufacturer’s instructions. Cells were then washed twice with fresh T cell medium and split into two different stimulation conditions. For the acute stimulation group, cells were maintained in T cell medium alone. For the chronic stimulation group with antibody cocktail, cells were cultured in T cell medium supplemented with ImmunoCult Human CD3/CD28 T Cell Activator (Stem Cell Technologies, 10971), following the manufacturer’s protocol. For the chronic stimulation group with anti-CD3 antibody (eBioscience, 16-0037-81), the antibody was coated at 12-well plate with 5 µg/ml concentration. Cells in the chronic condition were passaged every 48 h, with medium replacement and re-activate for three consecutive stimulation rounds. At the end point, cells were harvested and divided into two groups for either surface staining of exhaustion markers or intracellular cytokine analysis by flow cytometry.

### Lentiviral transduction and titration in CD8⁺ T cells

To determine optimal lentiviral transduction conditions for primary human CD8⁺ T cells, cells were activated for 48 h using CD3/CD28 beads as described above. After bead removal, cells were plated in 96-well plates at 2 × 10⁵ cells per well in 200 μl of T cell medium (X-VIVO 20 supplemented with 5% human AB serum and 10 ng/ml IL-2). Serial dilutions of lentivirus encoding puromycin resistance gene were added to the wells in the presence of 8 μg/ml polybrene (Millipore, TR-1003-G). Plates were centrifuged at 1,500g for 90 min at 37 °C, the medium was changed to T cell medium after additional 4 h incubation, the cell was continued culture at 37 °C, 5% CO2. At 48 h post-transduction, cells were diluted to 1 × 10⁵ cells per ml and cultured in the presence of 1 μg/ml puromycin for 48 h cell viability was assessed by counting viable cells using trypan blue. Viral titer and multiplicity of infection (MOI) were calculated based on the fraction of puromycin-resistant cells relative to total input.

### CRISPR screening in primary CD8⁺ T cells

Human CD8⁺ T cells were isolated from PBMCs by negative selection and activated with CD3/CD28 beads for 24 h After bead removal, cells were transduced with pooled lentiviral sgRNA libraries at an MOI of 0.3 in the presence of 8 μg/ml polybrene. Cells were spinoculated at 1000g and 32 °C for 90 min. 6 h after spin infection, the cell were resuspended to T cell medium for overnight culture. 24 hours after spin infection, the cell were treated with puromycin (1.5 μg/ml) to select the transduced cells. Following selection, cells were washed and resuspended into two treatment conditions corresponding to acute and chronic stimulation, as described previously. Cells were maintained under these conditions for 10 days, with medium refreshed and IL-2 re-supplemented every 2 days. On the final day of the screen, genomic DNA was harvested from each condition using the QIAamp DNA Blood Maxi Kit (QIAGEN, 51192) following the manufacturer’s instructions. Genomic DNA was quantified using Qubit dsDNA BR Assay Kit (Thermo Fisher Scientific, Q32850), and sgRNA sequences were PCR-amplified from extracted DNA using a one-step amplification protocol compatible with next-generation sequencing, as previously described by Broad Institute Genetic Perturbation Platform (“sgRNA/shRNA/ORF PCR for Illumina Sequencing” linked through https://portals.broadinstitute.org/gpp/public/resources/protocols), PCR primers were provided in Supplementary File 4. PCR products were purified using AMPure XP beads (Beckman Coulter, A63880). Libraries were quantified, normalized, and sequenced on an Illumina NovaSeq X platform using a 25B flow cell, generating paired-end 150 bp reads. Raw sequencing data were demultiplexed using bcl2fastq (Illumina) and further processed as described below.

### CRISPR screening analysis

Sequencing reads from CRISPR screens were first demultiplexed and trimmed to extract sgRNA sequences. Reads were aligned to the sgRNA reference library using the MAGeCK count function in the MAGeCK software package with default parameters^77^. Only sgRNAs with perfect matches to the reference library were retained for downstream analysis. Read count matrices were generated for all samples and normalized using median normalization. Differential sgRNA abundance between experimental conditions (Chronic vs. Acute stimulation) was analyzed using the MAGeCK test module, which applies a negative binomial model to assess gene-level enrichment or depletion. For each gene, MAGeCK aggregates sgRNA-level statistics to compute a robust gene-level score and associated p value using a robust ranking aggregation (RRA) algorithm. Genes were ranked by their adjusted p values (Benjamini–Hochberg FDR correction), and significance thresholds were defined as indicated in the figure legends. All analyses were performed using MAGeCK with default settings unless otherwise noted.

### RNP electroporation in primary human CD8⁺ T cells

To perform Cas9 RNP-mediated gene knockout, two synthetic sgRNAs targeting the same gene were used per condition (sequence available in Supplementary File 4). Each sgRNA was resuspended to a final concentration of 100 μM in RNase-free water and stored at −80 °C until use. Cas9 protein (EnGen® SpyCas9 NLS, 20 μM stock, New England Biolabs, M0646T) was used for complex formation. For each reaction, 60 pmol of Cas9 protein was complexed with 180 pmol total sgRNA by mixing 3 μl of Cas9 protein, 0.9 μl of each sgRNA, and 0.4 μl of 10× NEBuffer 3.1 in a total volume of 5.2 μl. The mixture was incubated at room temperature for 15 min to allow RNP formation. Primary CD8⁺ T cells were activated for 48 h with CD3/CD28 beads as described above, then washed once with 1× PBS and resuspended in Lonza P3 electroporation buffer at a density of 1 × 10⁶ cells per 20 μl. The entire 5.2 μl RNP mixture was added to 20 μl of cell suspension, mixed gently, and transferred into a well of a 16-well Nucleocuvette™ strip (Lonza). Air bubbles were removed by tapping the strip on the benchtop to prevent disruption of the electric pulse. Electroporation was performed using the Lonza 4D-Nucleofector system with the program EH115, optimized for primary human T cells. Immediately after electroporation, 80 μl of pre-warmed T cell medium (X-VIVO 20 supplemented with 5% human AB serum and 10 ng ml⁻¹ IL-2) was added to each well. The cells were incubated in the cuvette strip at 37 °C for 10 min to enhance recovery before being gently transferred to pre-warmed 24-well plates.

### Flow cytometry staining and analysis

For intracellular cytokine staining, T cells were stimulated for 3 h at 37 °C with Cell Activation Cocktail containing Brefeldin A (BioLegend, 423303). Cells were washed with Cell Staining Buffer (BioLegend, 420201), fixed and permeabilized with Fixation/Permeabilization solution (BD Biosciences, AB_2869008) for 20 min at 4 °C, and washed twice with 1× Perm/Wash buffer. After Fc receptor blocking (BioLegend, 422302), cells were stained for 2 h at 4 °C in the dark with fluorophore-conjugated antibodies in staining buffer (BioLegend, 420201) along with NIR Live/Dead dye (1:10000 dilution, ThermoFisher, L34975). Antibodies included APC anti-human CD8 Antibody (BioLegend, 344722), Brilliant Violet 785™ anti-human IFN-γ Antibody (BioLegend, 502541), Brilliant Violet 605™ anti-human TNF-α Antibody (BioLegend, 502935), and Brilliant Violet 421™ anti-human IL-2 Antibody (BioLegend, 500327).

For surface marker staining, cells were blocked with FC receptor blocking reagent (BioLegend, 422302), then stained with antibodies and NIR Live/Dead dye (1:10000 dilution, ThermoFisher, L34975) at 4 °C for 1 h. Antibodies included CD8 Antibody (BioLegend, 344722), Brilliant Violet 785™ anti-human TIGIT (VSTM3) Antibody (BioLegend, 372735), Brilliant Violet 605™ anti-human CD366 (Tim-3) Antibody (BioLegend, 345017), Brilliant Violet 510™ anti-human CD223 (LAG-3) Antibody (BioLegend, 369317), and Brilliant Violet 421™ anti-human CD279 (PD-1) Antibody (BioLegend, 329920). Flow cytometry was performed on a CytoFLEX LE (Beckman Coulter, six-laser configuration). Gating strategy included sequential gating on FSC-A/SSC-A (lymphocytes), FSC-A/FSC-H (single cells), and exclusion of dead cells using viability dye. Fluorescence compensation was calculated using single-stained compensation beads (BioLegend). Data were initially analyzed with FlowJo v10.8, with gates being defined using isotype controls, followed by custom reproduction of the analysis for visualization purpose only.

### Surveyor assay for genome editing efficiency

To evaluate genome editing efficiency following RNP electroporation, genomic DNA was extracted from T cells 72 h post-nucleofection using the Monarch® Genomic DNA Purification Kit (New England Biolabs, T3010S), following the manufacturer’s instructions. Genomic regions flanking the Cas9 cleavage site were PCR-amplified using 2× Phanta Max Master Mix (Vazyme, P525-02) in a 50 μl reaction volume (PCR primers provided in Supplementary File 4). PCR products were verified by agarose gel electrophoresis and purified using the Zymo DNA Clean & Concentrator Kit (Zymo Research, D4014), with final elution in 30 μl of nuclease-free water. To generate heteroduplex DNA for mismatch detection, 200 ng of purified PCR product was combined with 2 μl of 10× NEBuffer 2 (NEB) and nuclease-free water to a total volume of 19 μl. The mixture was denatured and reannealed in a thermocycler using the following program: 95 °C for 5 min; ramp at −2 °C/s to 85 °C; then ramp at −0.1 °C/s to 25 °C; and held at 4 °C. After annealing, 1 μl of T7 Endonuclease I (NEB, M0302) was added to each reaction and incubated at 37 °C for 60 min. Reactions were terminated by adding 1.5 μl of 0.25 M EDTA and analyzed by electrophoresis on a 1% agarose gel to visualize cleavage products.

### mRNA knockdown analysis by quantitative PCR

To assess transcript-level gene knockdown following RNP electroporation, edited T cells were collected at the end point for RNA extraction. Total RNA was isolated using a RNeasy Mini Kit (Qiagen, 74104), and RNA concentration was quantified by nanodrop. Reverse transcription was performed using the HiScript® III 1st Strand cDNA Synthesis Kit (+gDNA wiper) (Vazyme, R312-01) according to the manufacturer’s instructions. Gene-specific primers targeting the transcript of interest were used during amplification (Supplementary File 4). Quantitative PCR was performed using PowerUp™ SYBR® Green Master Mix (Thermo Fisher Scientific, A25742) on a CFX96 Real-Time PCR Detection System (Bio-Rad). ACTB (β-actin) was used as the internal control gene for normalization. Each sample was run in triplicates, and specificity of amplification was confirmed by melt-curve analysis. Relative gene expression was quantified using the ΔΔCt method, and results were reported as fold change compared to the control condition.

### RNA-seq analyses

Total RNA was extracted on the final day from ZNF766 knockout and NTC control groups using the RNeasy Mini Kit (Qiagen, 74004). Two biological replicates were collected per condition. RNA-seq libraries were generated with the NEBNext Ultra II Directional RNA Library Prep Kit (NEB, E7765L) following the manufacturer’s poly(A) mRNA workflow, including enrichment with the NEBNext Poly(A) mRNA Magnetic Isolation Module (NEB, E7490L). Raw RNA-seq reads were mapped to the human reference genome (GRCh38/hg38) using STAR (v2.7.2a) with default settings. Strand-specific read counts were aggregated into a gene-level count matrix for subsequent analyses. Transcriptomic variation across samples was evaluated by principal component analysis (PCA) based on the 500 most variable genes. Differential expression analysis was conducted using DESeq2 (v1.40.2), and genes with an absolute log2 fold change greater than 0.5 and an adjusted P-value below 0.05 were defined as significantly differentially expressed. Functional enrichment analyses were performed using clusterProfiler (v4.10.0), together with org.Hs.eg.db (v3.18.0), enrichplot (v1.22.0), and DOSE (v3.28.2). Significant genes were converted to Entrez Gene identifiers and assessed for enrichment in Gene Ontology biological processes and KEGG pathways. Enrichment results were visualized as dot plots (GO) and bar plots (KEGG).

## Data Availability

All ATAC-seq, ChIP-seq used in this study are public data collected from the ENCODE data portal (https://www.encodeproject.org). T cell scATAC-seq data is obtained from Gene Expression Omnibus (GEO) under accession GSE129785. Purkinje neuron developmental scATAC-seq data is obtained from https://github.com/linnarsson-lab/fetal_brain_multiomics. CRISPR screen data is available on GEO under accession number GSE305611. RNA-seq data on ZNF766 KO is available on GEO under accession number GSE305610.

## Code Availability

The code for Chromnitron is available at https://github.com/tanjimin/Chromnitron.

## Acknowledgement

This work is supported by NIH grants DP5OD033430 and R03DE033371 (to B.X.), R01CA252239 and P01CA229086 (to A.T.). X.F. and R.R. are supported by R35CA253126 and P30CA013696. J.T. was supported by the Judith & Stewart Colton Center for Autoimmunity at NYU Langone and Arthritis National Research Foundation Grant Award (No. 1471427). X.L. is a Fellow of The Jane Coffin Childs Fund for Medical Research. B.X. is a Junior Fellow of the Harvard Society of Fellows. We thank the High Performance Computing Core Facility at NYU Grossman School of Medicine and the Broad Institute for the support of computation. We thank the Broad Genetic Perturbation Platform and Broad Clinical Labs for the support of genetic and genomic experiments. We thank Sonya Craig from scientific editing team at NYU Langone for editing the manuscript. We also acknowledge Milk Bar, where X.F. and J.T. enjoyed coffee and met weekly for over two years to work on Chromnitron. We thank Huiyuan Zhang, Chenzhen Zhang, Hanqing Liu, Lloyd Bod, Jason D. Buenrostro, Eric S. Lander, Anshul Kundaje, Romanos S. Pistofidis, Liz Gaskell, Wenke Liu, Jose U. Scher, and the members of the Xia lab and Tsirigos lab for their valuable suggestions and discussion. Lastly, we are grateful to the ENCODE consortium for generating, organizing, and sharing standardized genomics data that enabled this study.

## Author Contribution

B.X. and J.T. conceived the project. J.T., B.X. and A.T. designed the computational and experimental components and interpreted the results, with inputs from X.F. and X.L.. J.T., F.X. and S.M. designed, implemented, and optimized the neural network. J.T. and X.F. built the preprocessing platform. X.F. performed single cell data preprocessing and protein feature analysis. J.T. trained the model, performed inference and downstream computational analysis. X.L. performed the experimental components. J.B. performed RNA-seq analysis. R.R., D.F., and J.D.B. provided feedback and contributed to discussion. J.T. and B.X. led the writing of the manuscript with input from X.F., X.L. and A.T. All authors commented on the manuscript.

## Supplementary Figures

**Supplementary Fig. 1:**
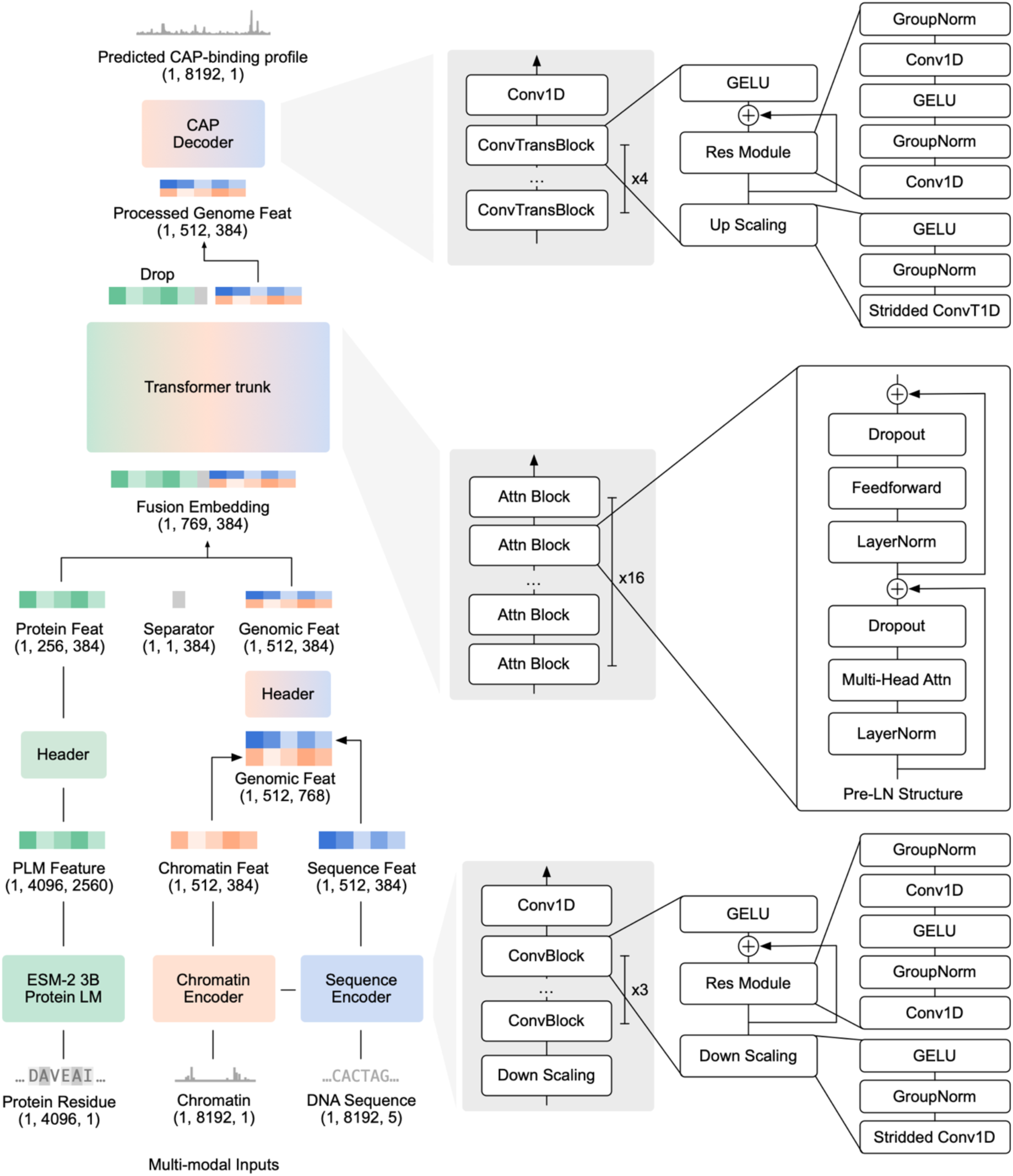
Chromnitron model architecture. Left, Overview of model structure, showing modality-specific encoders for DNA sequence, chromatin accessibility, and protein sequence, followed by multimodal fusion, transformer trunk, and a CAP-specific decoder that outputs base-resolution binding profiles. Right, Detailed layer configurations for the convolution encoders, transformer attention blocks, and decoder modules. Convolutional blocks include Group Normalization, Gaussian Error Linear Unit (GELU) activation, and residual connections with strided down-scaling or transposed up-scaling layers as indicated. Transformer layers use a pre-layer-normalized (Pre-LN) architecture with multi-head self-attention and feedforward sublayers. Dimensions of inputs, intermediate representations, and outputs are indicated in brackets as (batch size, length, channels).

**Supplementary Fig. 2:**
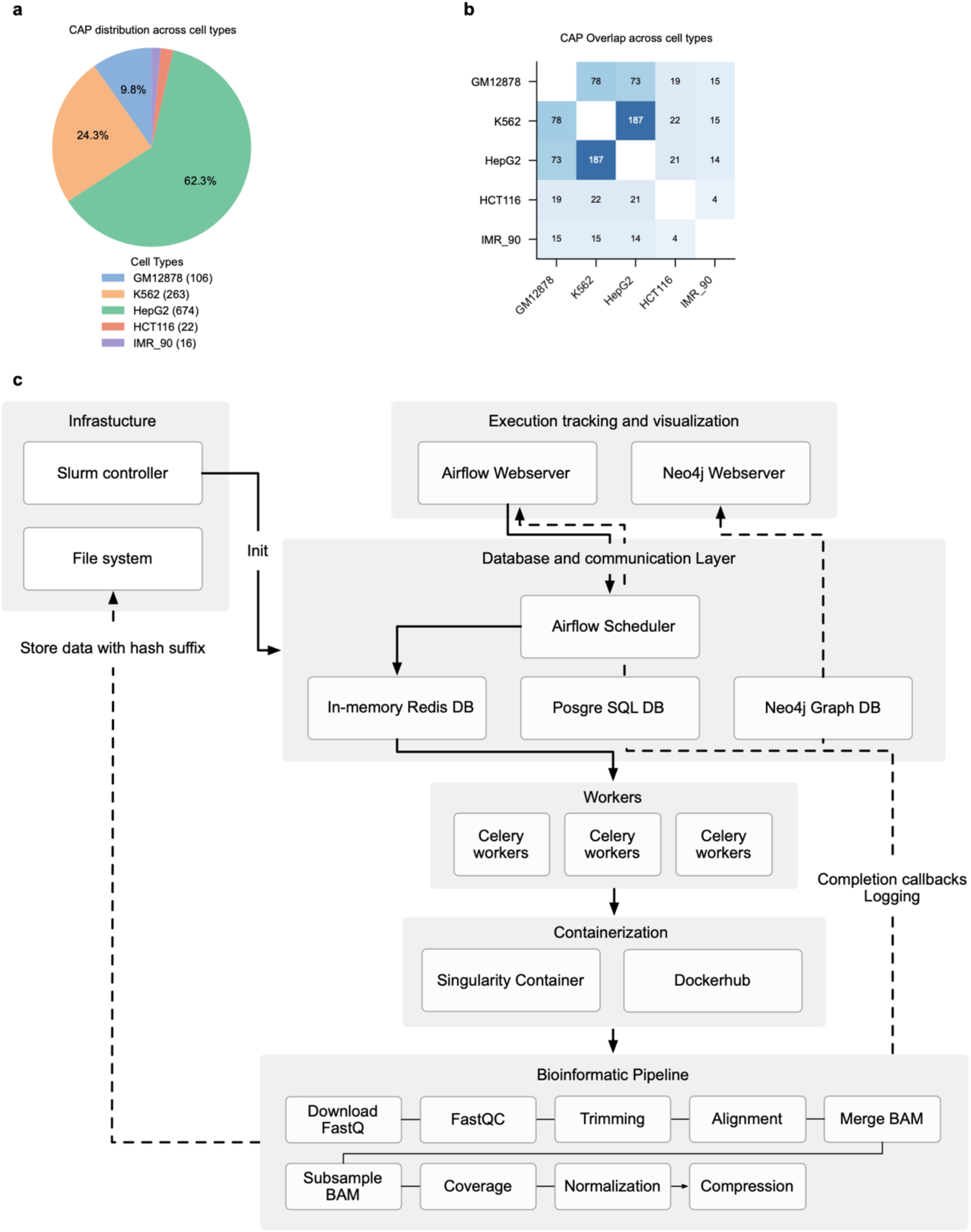
Training data composition and the Chrom2Vec preprocessing platform. **a.** Distribution of curated CAP-binding ChIP-seq datasets across five human cell lines. **b.** Pairwise overlap of CAP-binding datasets between cell lines, displayed as a heatmap indicating the number of shared CAPs. Diagonal entries are removed. **c.** Workflow of Chrom2Vec platform for processing training datasets. The workflow integrates job scheduling, data storage, execution tracking, containerized bioinformatic pipelines, and automated normalization and compression steps. Full details about the platform were described in Methods.

**Supplementary Fig. 3:**
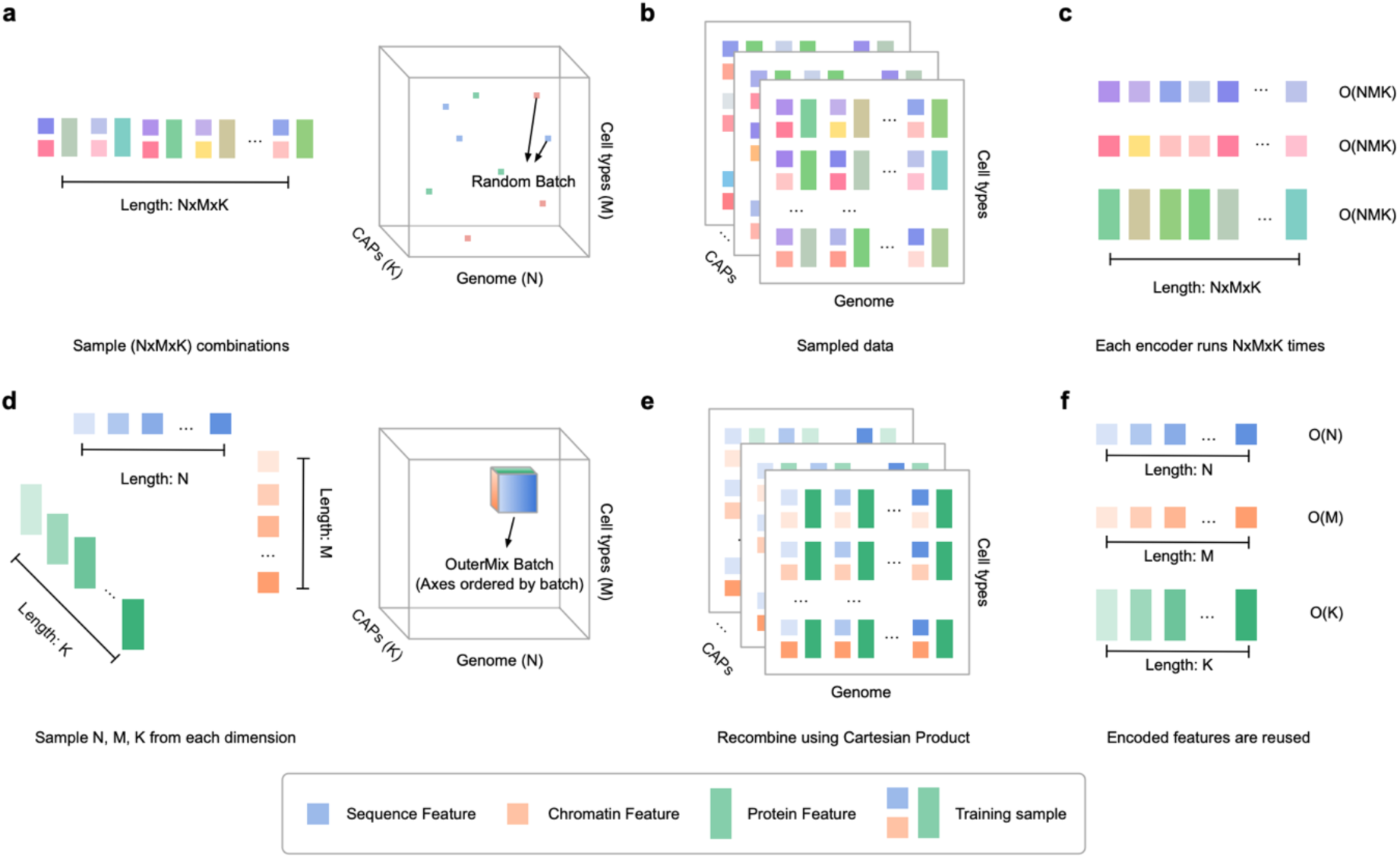
OuterMix algorithm improves multimodal training efficiency via factorized batch sampling. **a-c.** Sampling and computation with random sampling strategy. A randomly sampled batch has *N×M×K* combinations of genome sequence (N), chromatin feature (M), and CAP (K), selected from the full three-dimensional data manifold (**a**). This random processing approach generates large data matrices for encoding (**b**). Thus, each encoder for the corresponding modality would process every combination independently, yielding a computational complexity of O(*N×M×K*) per encoder (**c**). **d-f.** Sampling and computation with OuterMix strategy. OuterMix first samples *N, M, K* training samples from each dimension, effectively sampling a dense data cube from the full three-dimensional data manifold (**d**). Each sampling was encoded, with the encoded embeddings being recombined via a Cartesian product to form *N×M×K* training samples (**e**). These encoded features are reused in the final Cartesian product and avoids redundant computation, reducing encoder computation complexity down to O(*N*) + O(*M*) + O(*K*).

**Supplementary Fig. 4:**
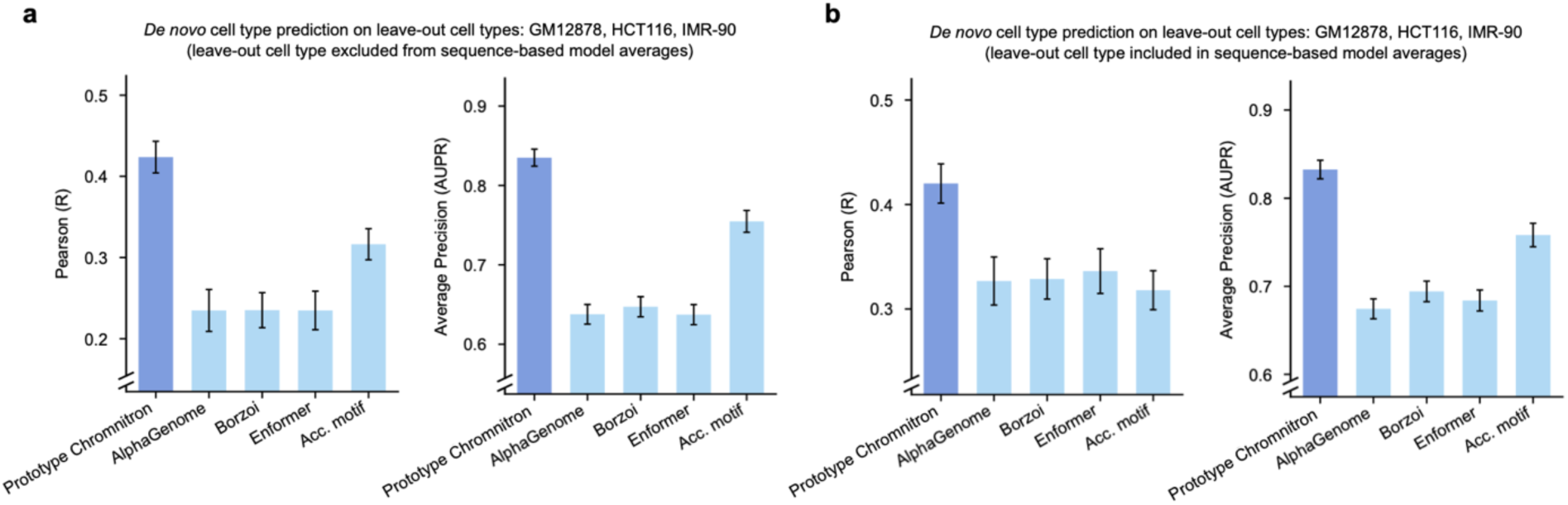
Cell-type-specific CAP-binding prediction performance of Prototype Chromnitron. **a.** Performance comparison between Prototype Chromnitron, trained on datasets from two cell types (HepG2 and K562), and other models in pair-wise cell-type-specific regions among GM12878, HCT116 and IMR-90. Cell-type-specific ATAC-seq was used as input for prediction and accessibility-weighted motif score (Acc. Motif) baseline. Average prediction across cell lines excluding the target cell type was used for sequence-based models (AlphaGenome, Borzoi, and Enformer). **b.** The same benchmark as in (**a**), except that sequence-based model predictions were averaged across all available cell types, including the validation cell line.

**Supplementary Fig. 5:**
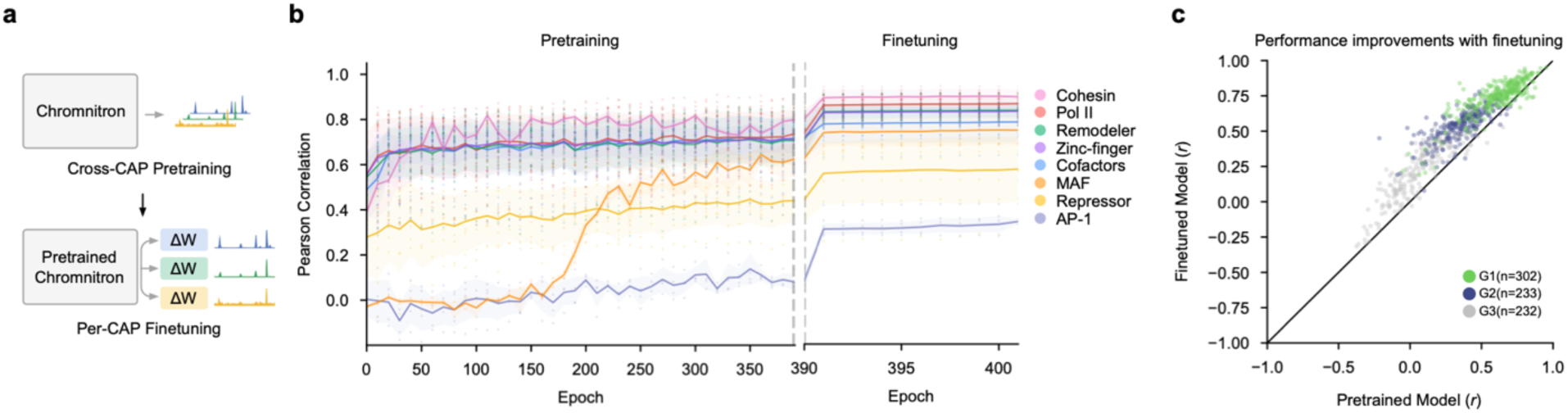
Full-scale Chromnitron pretraining and finetuning. **a.** Two-phase training strategy showing cross-CAP pre-training using shared weights (grey) followed by individual fine-tuning for each CAP with low-rank adaptation (LoRA), visualized as weight deltas (ΔW). **b.** Model performance across pretraining and fine-tuning epochs, grouped by CAP categories. Line and the shade represent scores of mean ± s.d. for each CAP category. **c.** Scatter plot of model performance (Pearson’s correlation between prediction and experimental data of individual CAPs) between fine-tuned model and pretrained model.

**Supplementary Fig. 6:**
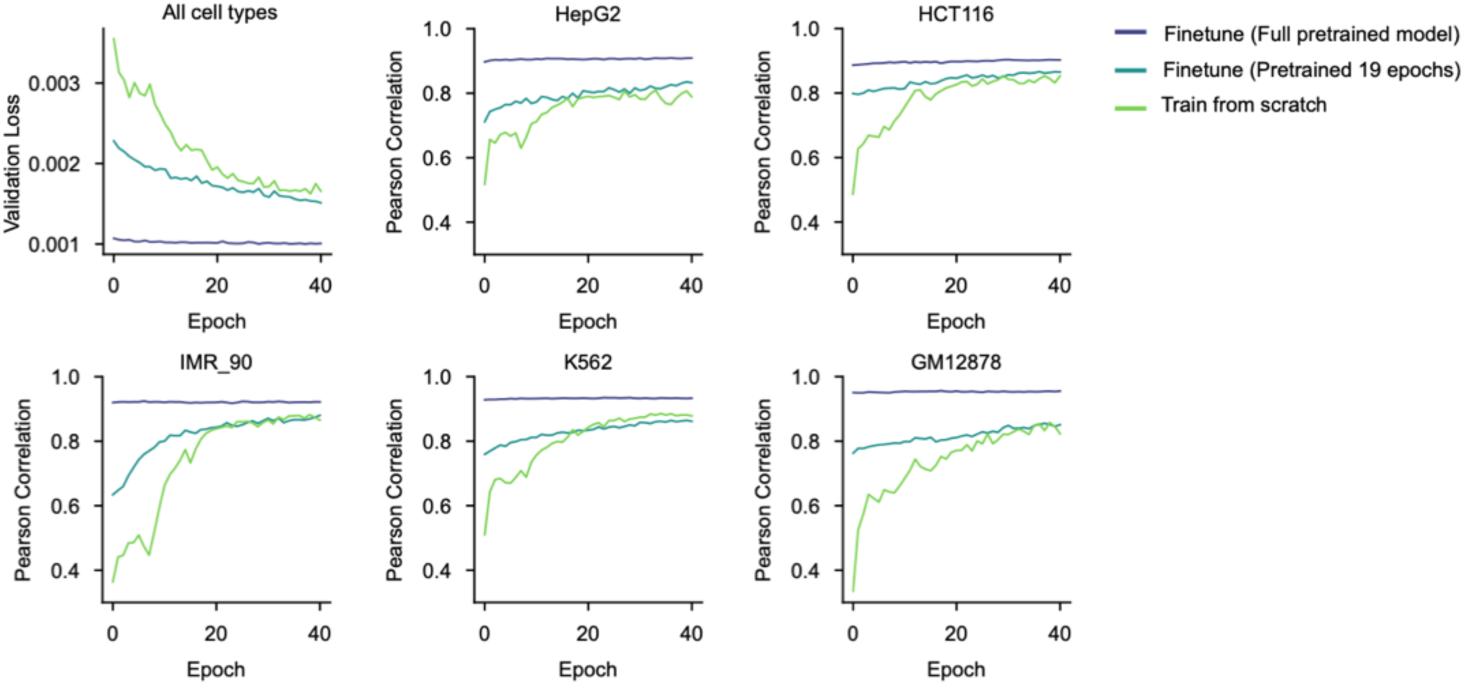
Model-training efficiency comparison between finetuning and training from scratch. Training curves comparisons across three model-training strategies: fine-tuning a fully pretrained model (dark blue), fine-tuning a checkpoint after 19 epochs of pretraining (light blue), and training from scratch (green). The comparison includes overall validation loss (across all cell types) and Pearson’s *r* relating to reference data in each cell type across training epochs.

**Supplementary Fig. 7:**
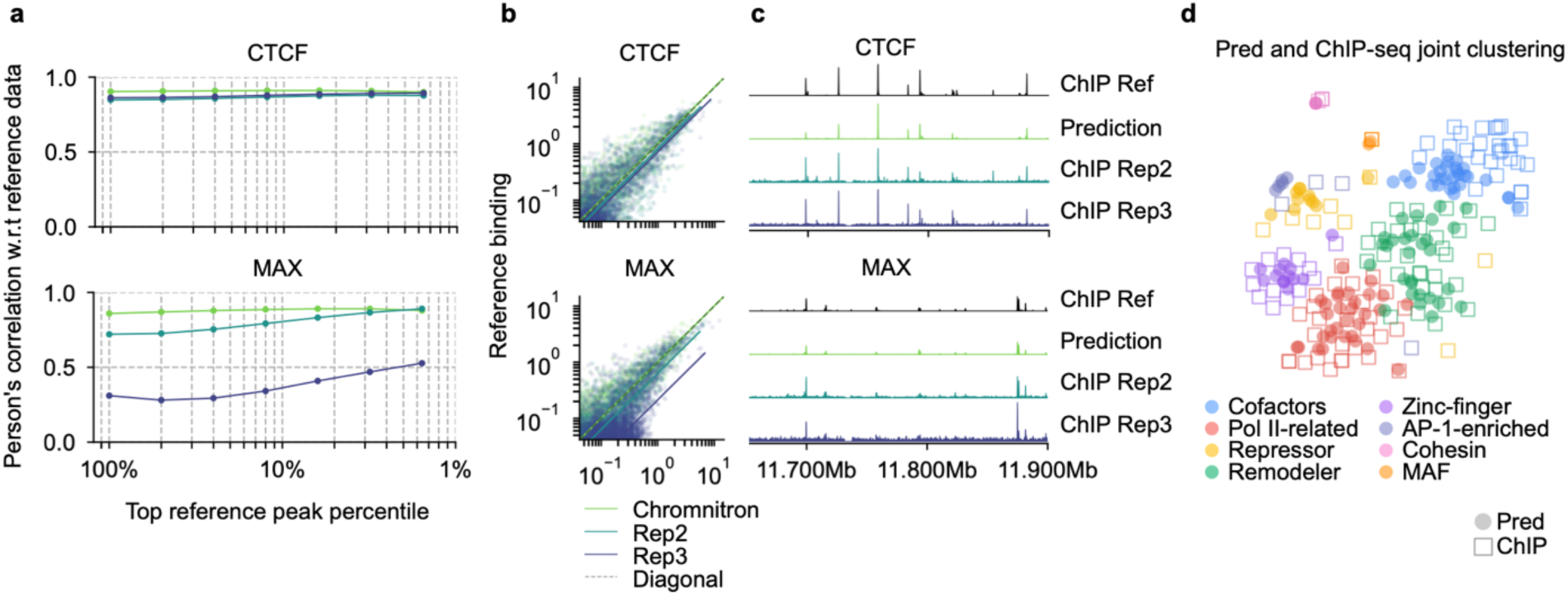
Chromnitron predictions match experimental ChIP-seq replicates. **a-c.** Evaluation of Chromnitron predictions against experimental ChIP-seq replicates for CTCF (top) and MAX (bottom). Each evaluation was performed by comparing prediction performance with that of replicating experimental measurements. These evaluations include Pearson’s correlation to the reference experimental data across different percentiles of genomic bins ranked by mean signal strength (**a**), scatter plot of predicted or experimental replicate binding signals (x-axis) versus the reference experimental data (y-axis, **b**), and direct visualization of CAP-binding tracks of prediction alongside two other experimental replicates (Rep2, Rep3), all aligning to the reference experimental data (Ref, **c**). **d.** Dimensionality reduction (t-SNE) visualization of predicted (circle) and experimental (squares) binding profiles from a subset of CAPs. Colors define CAPs clustered with similar binding profiles.

**Supplementary Fig. 8:**
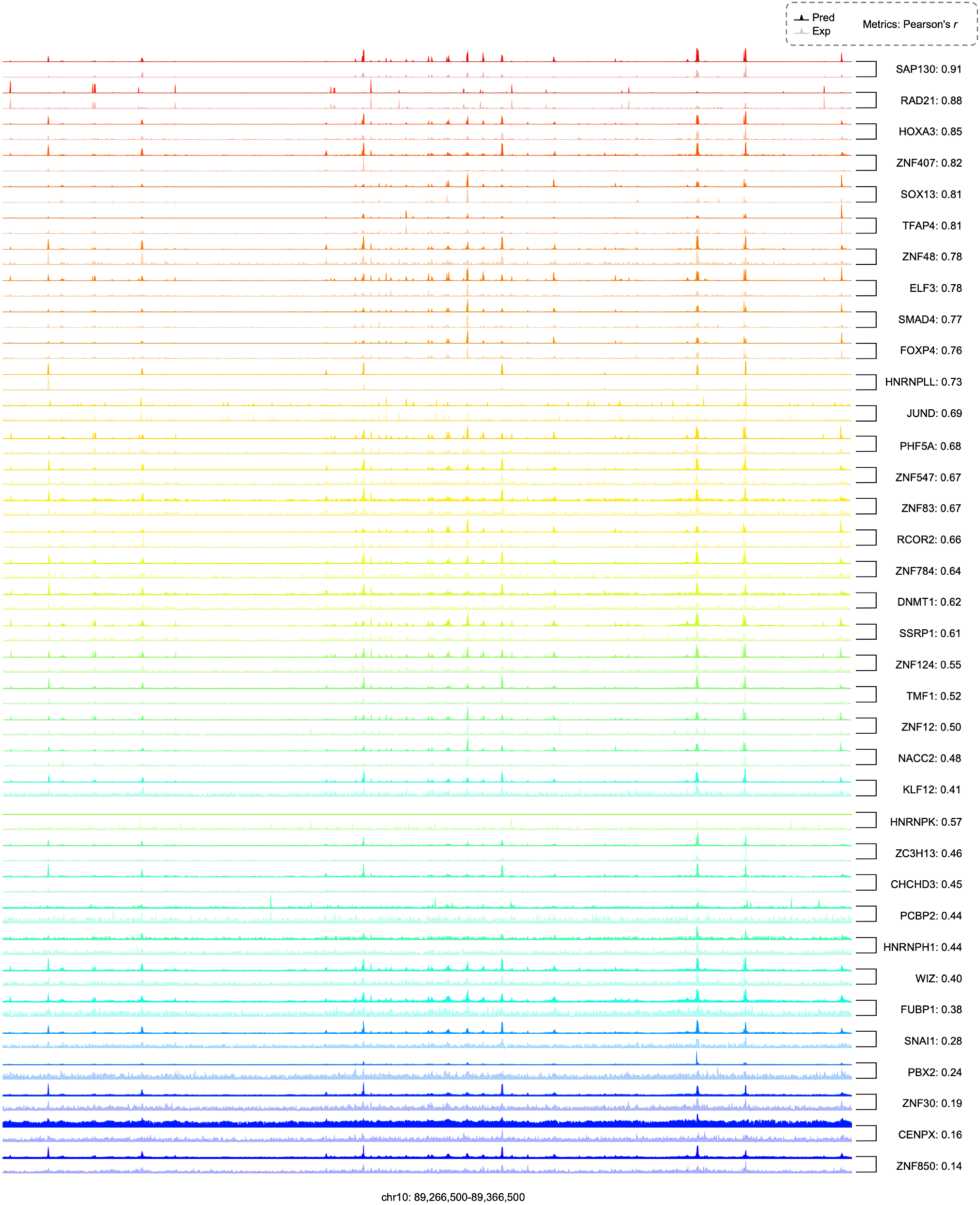
Paired visualization of Chromnitron-predicted and ChIP-seq-measured CAP binding profiles. Paired CAP-binding tracks of Chromnitron prediction (top) and experimental data (bottom) for 36 randomly selected CAPs across the leave-out genomic region (chr 10). CAP-binding tracks were colored by their performance (Pearson’s *r*).

**Supplementary Fig. 9:**
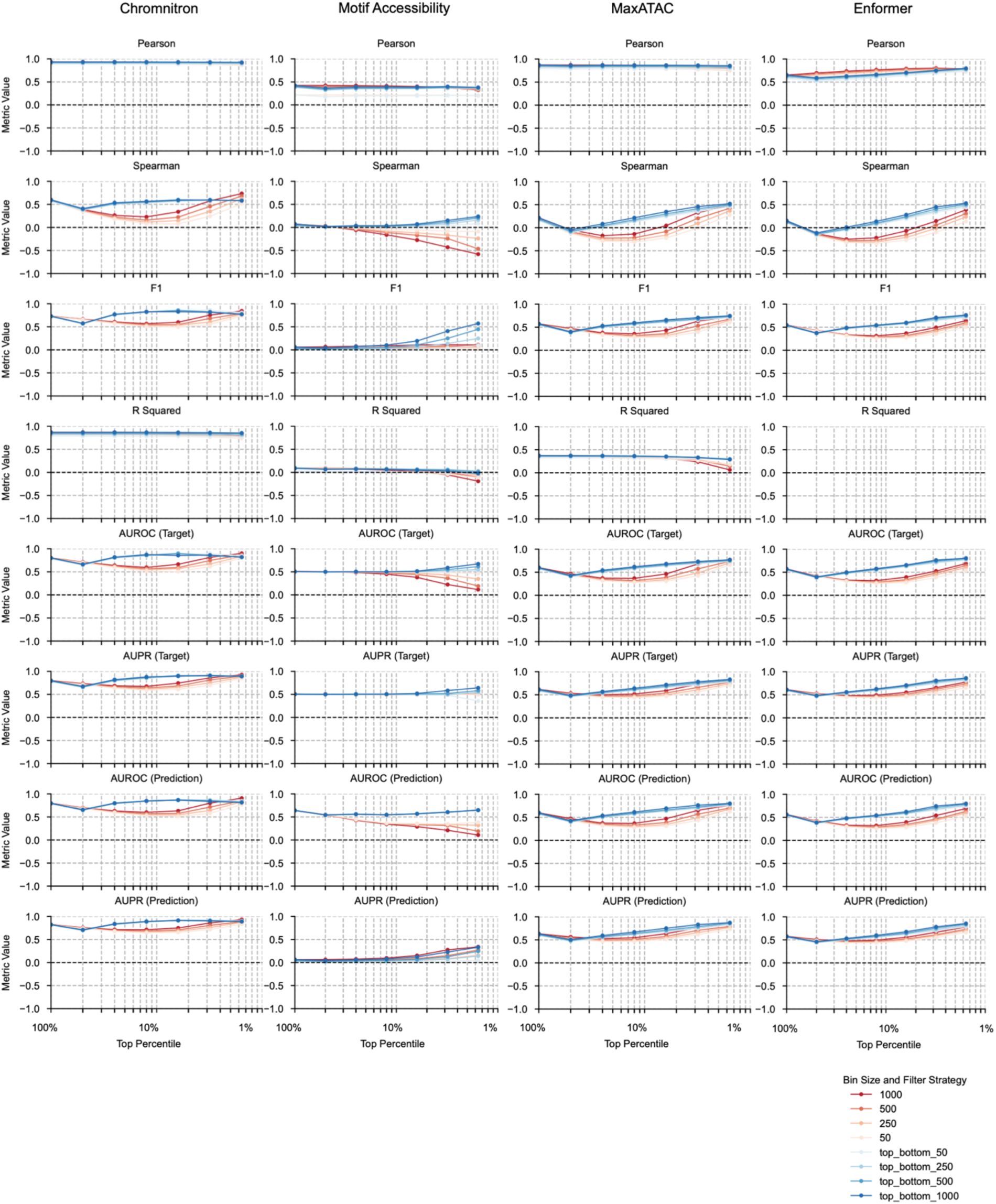
Performance metrics comparison between Chromnitron and three other models, using MAX as an example. All performance metrics were performed against reference experimental ChIP-seq data of MAX. A total of eight statistics were calculated, including Pearson’s correlation (*r*), Spearman’s correlation (*ρ*), the coefficient of determination (*R* squared), F1 score, and area under the receiver-operating characteristic (AUROC) and precision–recall (AUPR) curves, computed using both target-centric and prediction-centric labelling. Each of these methods was calculated using bins sizes of 50, 250, 500, and 1,000 bp (color intensity denotes bin size). Each plot shows a performance metric (y-axis) dynamic as the different percentages of genomic bins included in the analysis (x-axis, log-scaled). Red lines indicate selection of bins from the top percentiles only, while blue lines indicate selection of bins from both the top and bottom percentiles of bins.

**Supplementary Fig. 10:**
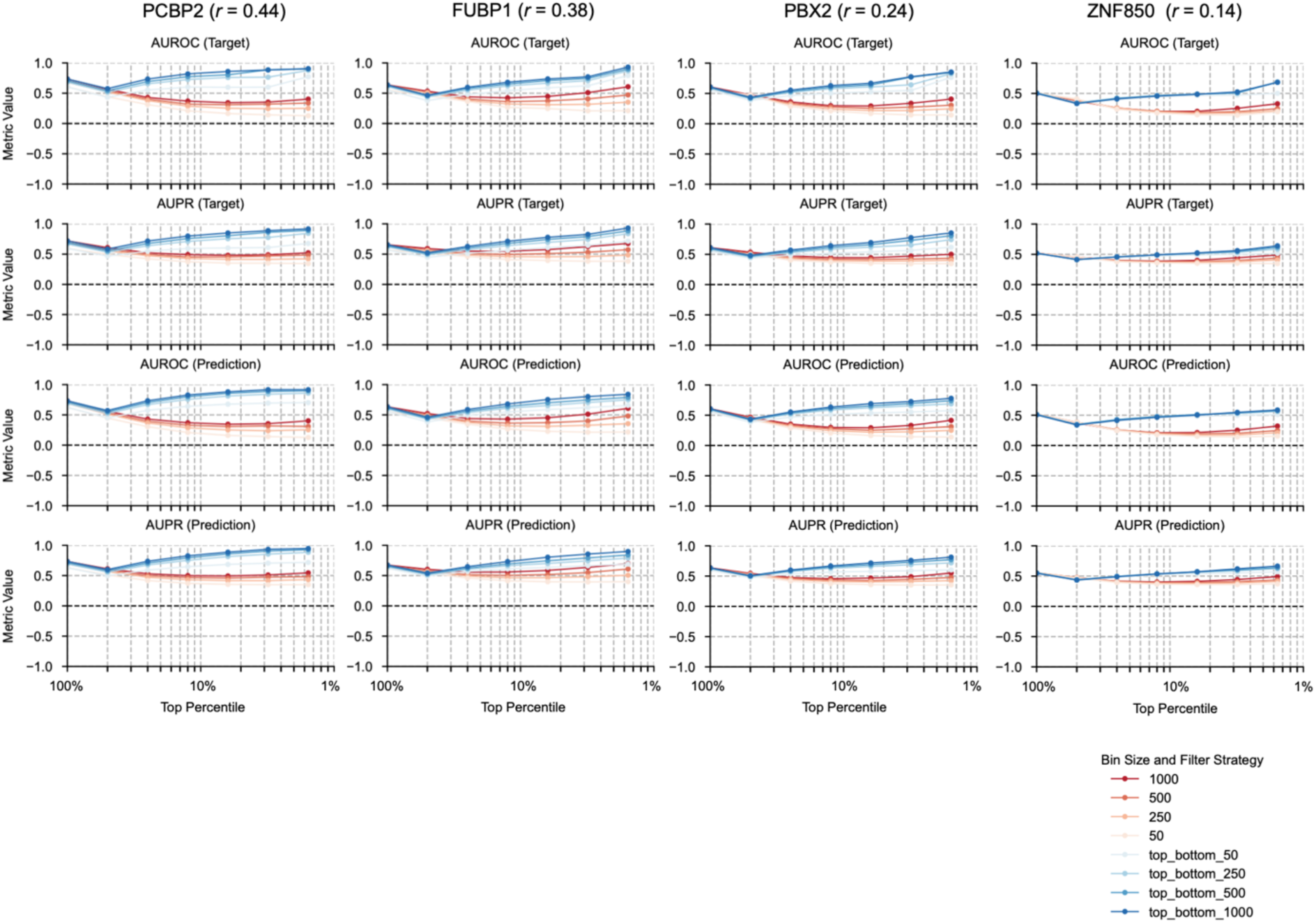
Robust peak-level performance for CAPs with low global correlation. Performance metrics for four randomly selected low-correlation (Pearson’s *r* < 0.5) CAPs, including PCBP2, FUBP1, PBX2, and ZNF850. Despite the modest correlation coefficients in these CAPs, Chromnitron-predicted CAP-binding peaks show high correspondence with reference ChIP-seq peaks, measured by top and bottom percentile AUPR and AUROC (blue line). Pearson’s correlation was calculated by comparing to reference experimental ChIP-seq data. For each CAP, four statistics were calculated, including AUROC and AUPR scores using both target-centric and prediction-centric labelling. Metrics were calculated using bin sizes across 50, 250, 500, and 1,000 bp (color intensity denotes bin size). Red curves indicate selection of bins from the top percentile only, while blue curves indicate selection of bins from both the top and bottom percentiles of bins.

**Supplementary Fig. 11:**
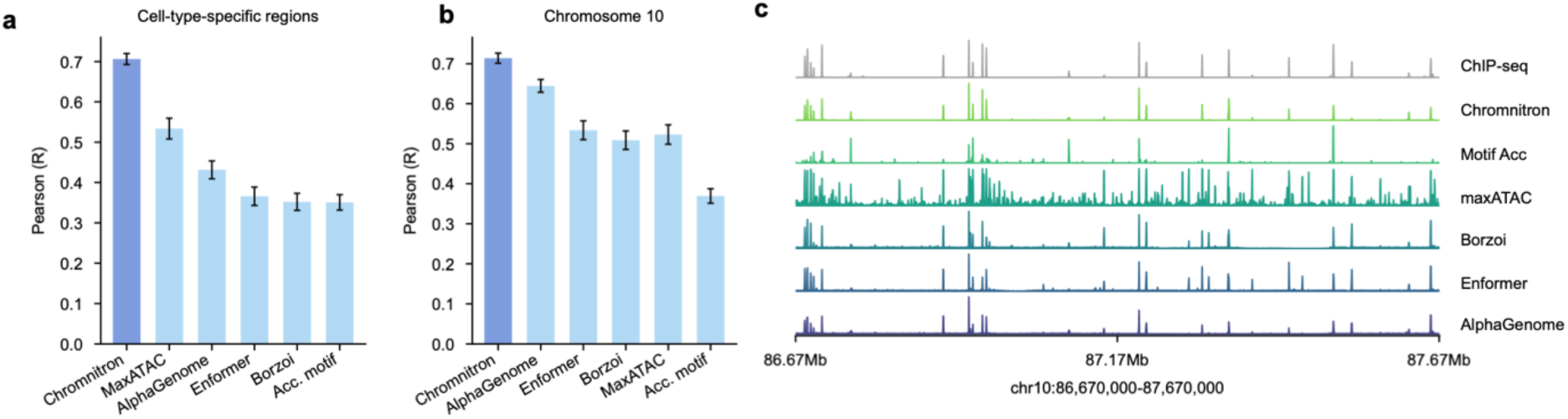
Chromnitron performance benchmark. **a.** Performance comparison between Chromnitron and other models in cell-type-specific regions between HepG2 and K562. Cell-type-specific ATAC-seq was used as input for Chromnitron, maxATAC and accessibility-weighted motif score (Acc. Motif) baseline. Average prediction across all cell lines was used for sequence-based models. **b.** Performance comparison between Chromnitron and other models across the entire Chromosome 10 in HepG2 and K562 cells, including both cell-type-specific and non-specific binding regions. **c.** Representative CAP binding tracks comparing Chromnitron, other models and experimental ChIP-seq measurement on chromosome 10.

**Supplementary Fig. 12:**
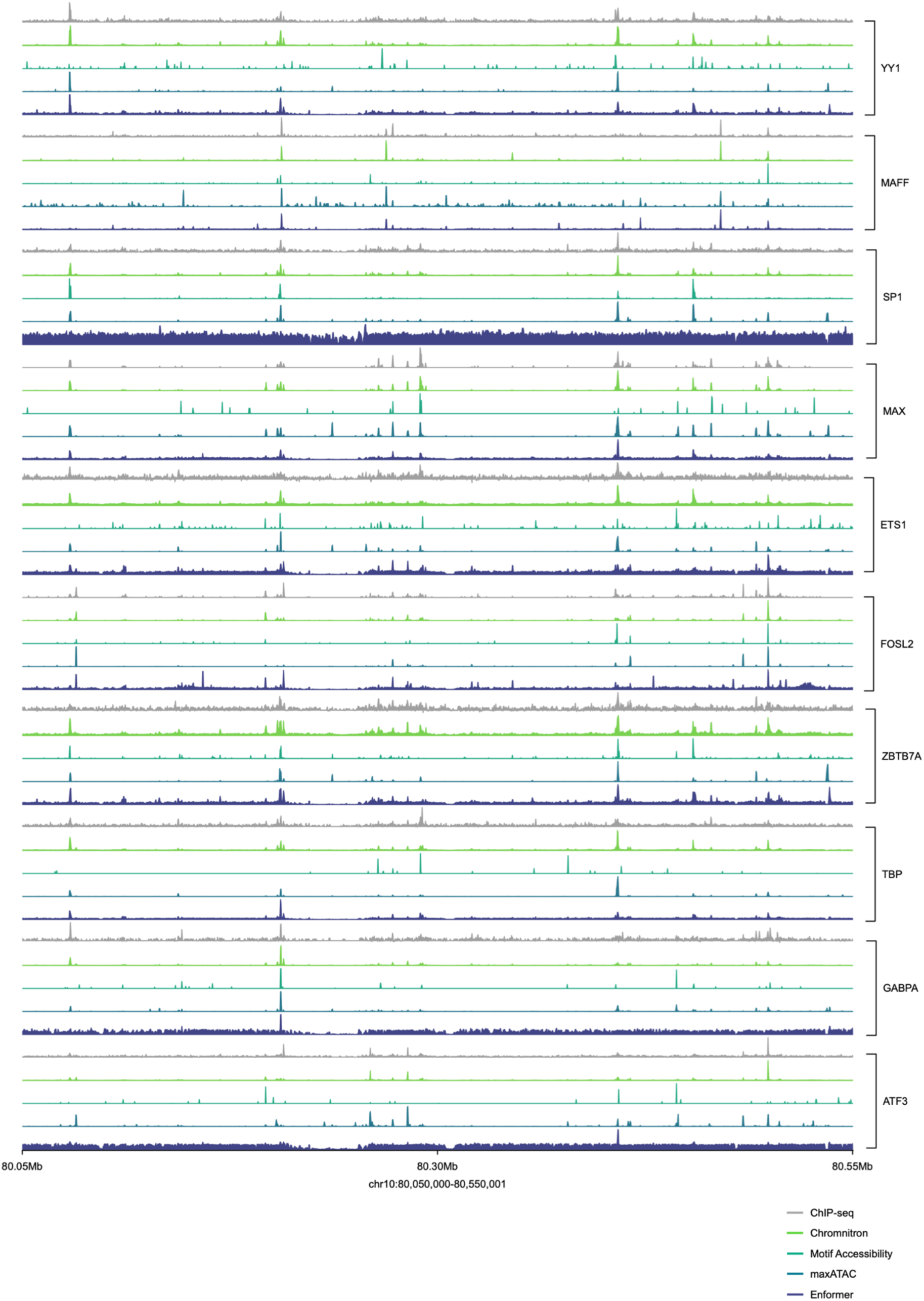
Comparison of CAP-binding predictions across models. Predicted CAP-binding profiles for 10 randomly selected CAPs were presented at the same 500kb genomic window in the leave-out chromosome 10. Each CAP includes predicted binding tracks from Chromnitron, accessibility-weighted motif score (Acc. Motif), maxATAC and Enformer models, all referring to the experimental ChIP-seq data in leave-out chromosome 10. Min-max scaling of the binding signal was applied to each track separately.

**Supplementary Fig. 13:**
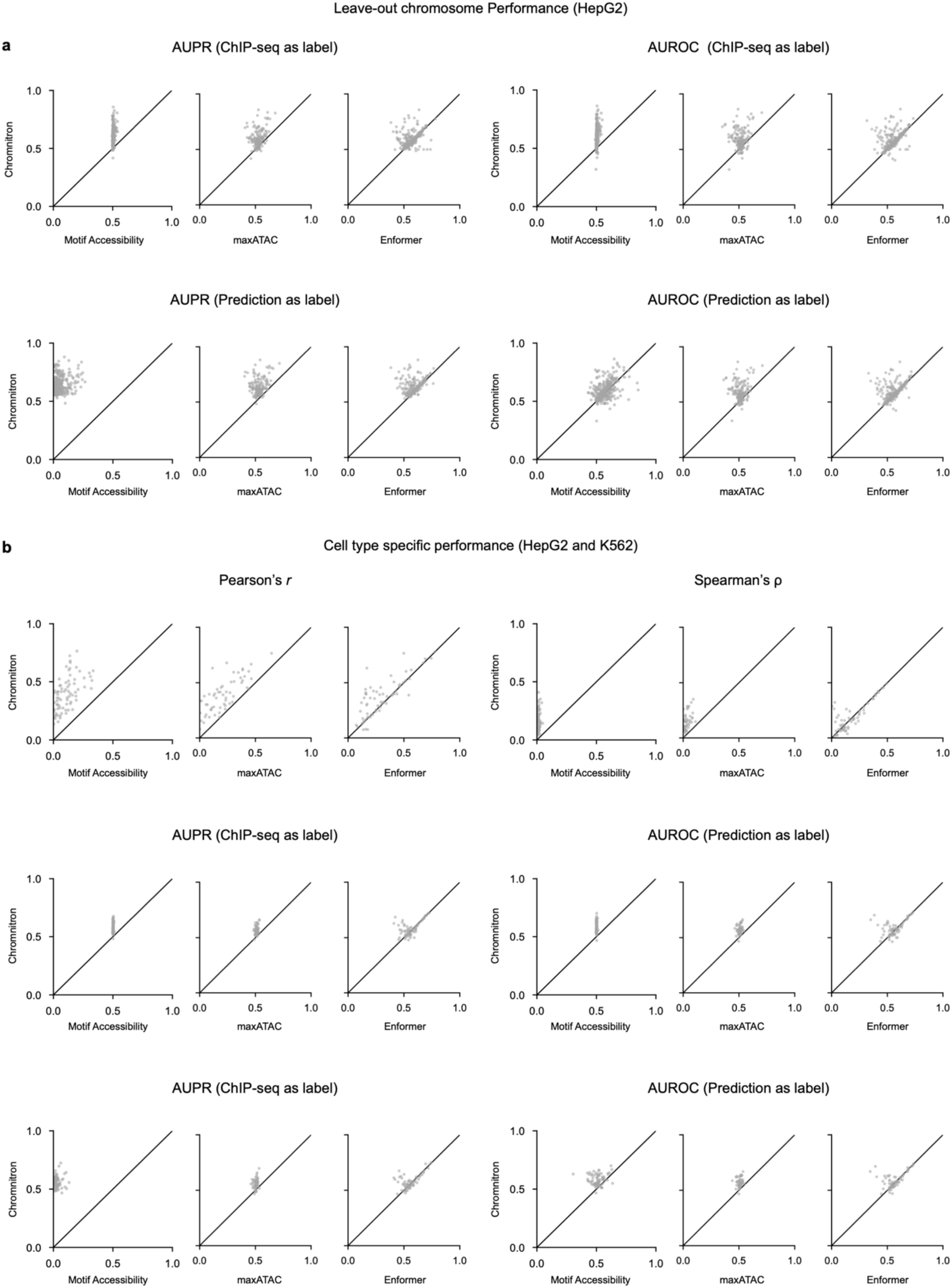
Performance statistics across cell types. **a.** Benchmarking prediction performance on Chromnitron’s leave-out genomic region (chromosome 10) in HepG2 cells. Each dot represents one CAP, plotted as Chromnitron prediction performance (y-axis) versus the comparison model (x-axis). Performance was quantified using AUPR and AUROC, calculated by using ChIP-seq (top) or prediction (bottom) as the reference label of binarized binding status. **b.** Performance benchmarking by predicting cell type-specific CAP-binding. The benchmarking was performed by comparing model-predicted binding difference between HepG2 and K562 cells (HepG2 – K562) to the experimental ChIP-seq difference (HepG2 – K562). Pearson’s *r* and Spearman’s *ρ* were included as evaluation metrics in addition to the AUPR and AUROC as in (**a**).

**Supplementary Fig. 14:**
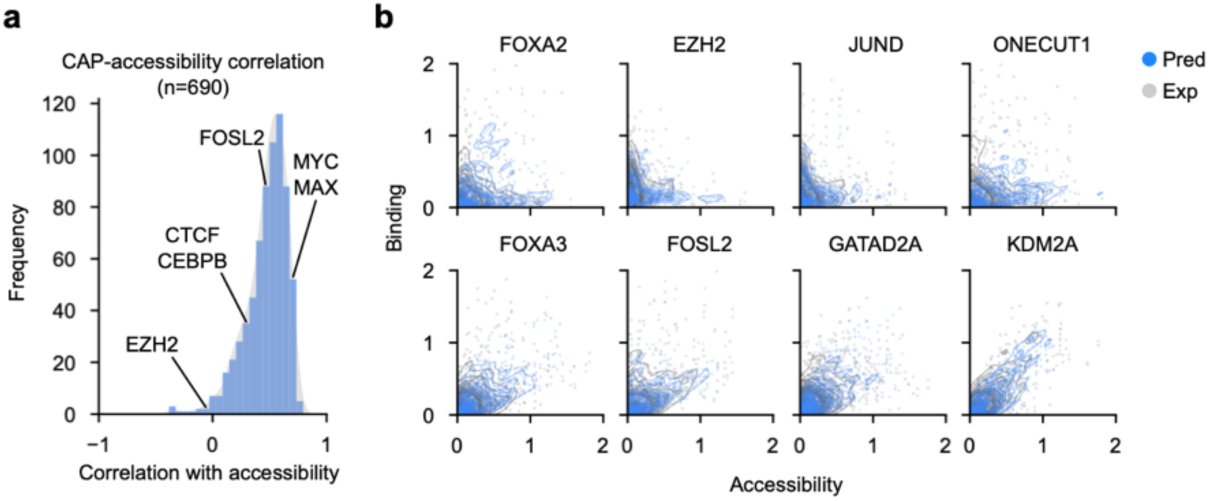
Comparing chromatin accessibility, CAP binding, and Chromnitron prediction. **a.** Distribution of correlation coefficients (Pearson’s *r*) between experimental ChIP-seq signal and chromatin accessibility in HepG2 cells. **b.** Scatter plots of CAP binding versus chromatin accessibility signals for experimental ChIP-seq signal (Exp, grey) and Chromnitron predictions (Pred, blue).

**Supplementary Fig. 15:**
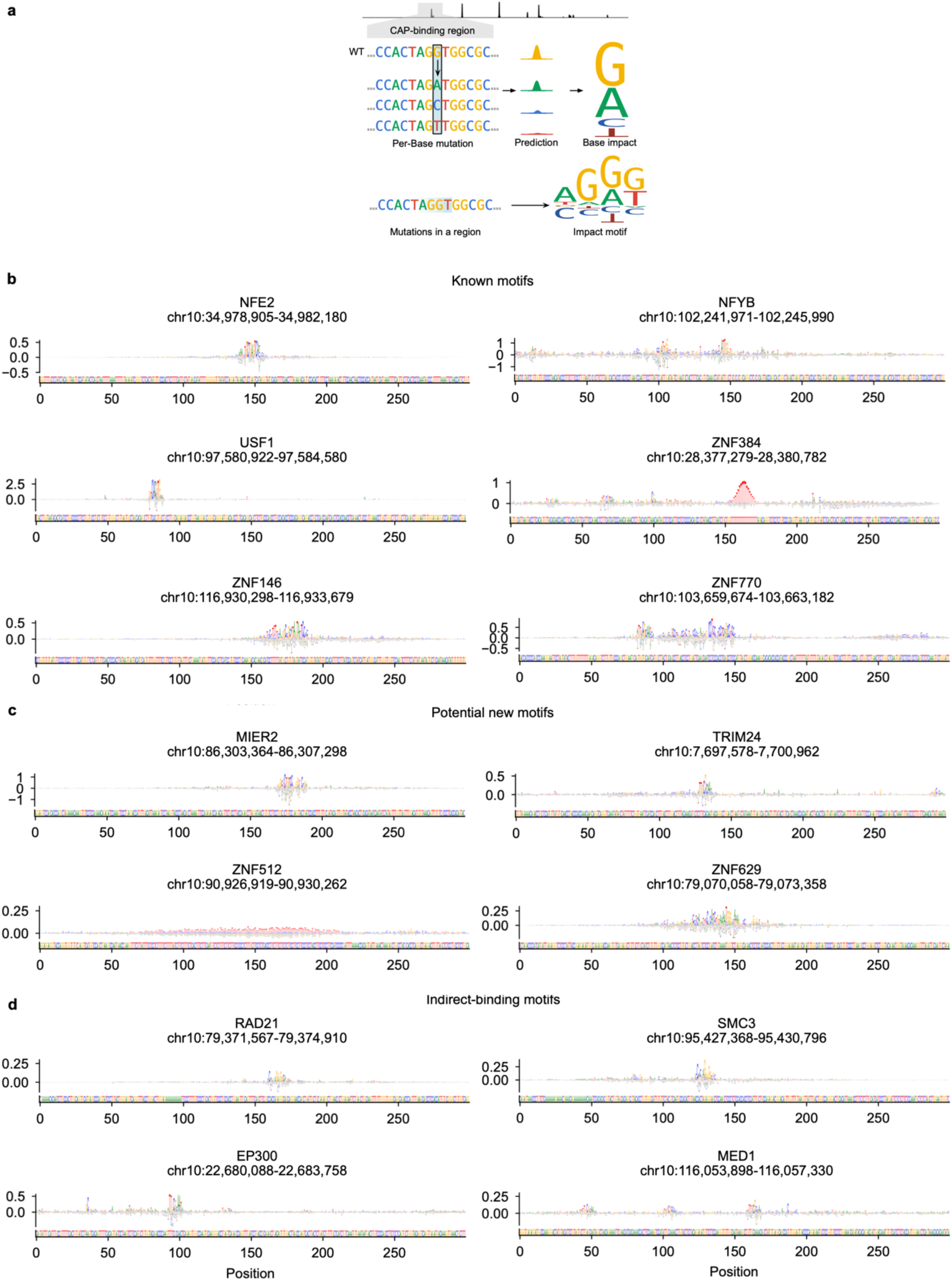
*In silico* DNA sequence mutagenesis identifies CAP impact motifs and enables variant effect prediction. **a.** Quantification of CAP-specific sequence impact scores through base-resolution *in silico* mutations and Chromnitron prediction. **b-d.** Chromnitron prediction-based *in silico* DNA sequence mutagenesis identifies CAP-specific impactful DNA bases (i.e. impact motifs) that resembles both known (**b**) and novel (**c**) motifs, as well as indirect binding motifs that resemble their interaction partner CAPs which binds DNA(**d**). Impact scores were presented in the form of sequence logos which were normalized per base to median value. Each impact motif was predicted at a specific locus as denoted in panel title.

**Supplementary Fig. 16:**
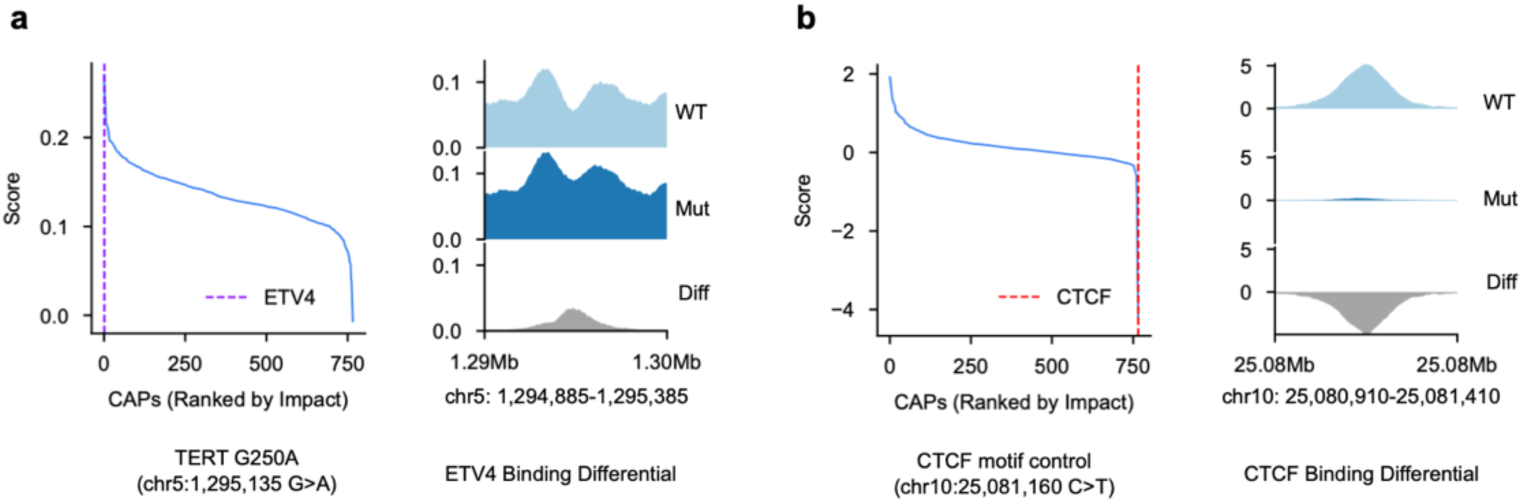
Chromnitron learns DNA sequence features at single-nucleotide resolution to enable cell-type-specific variant effect prediction. **a.** Chromnitron predicts gain-of-binding effect at *TERT* promoter G250A variant (hg38 chr5:1295135 G>A). Left: CAPs ranked by mutation impact score with a vertical dashed colored line marking ETV4, an ETS-family transcription factor. Right: binding profiles of ETV4 on wild type (WT), mutant (Mut) sequences, and their corresponding differential binding (Diff: Mut – WT). **b.** Chromnitron predicts loss-of-binding effect at a CTCF-binding site (hg38 chr10:25081160 C>T). Left: CAPs ranked by mutation impact score with red dashed line indicating CTCF. Right: CTCF binding profiles across WT and Mut sequences and their differential binding.

**Supplementary Fig. 17:**
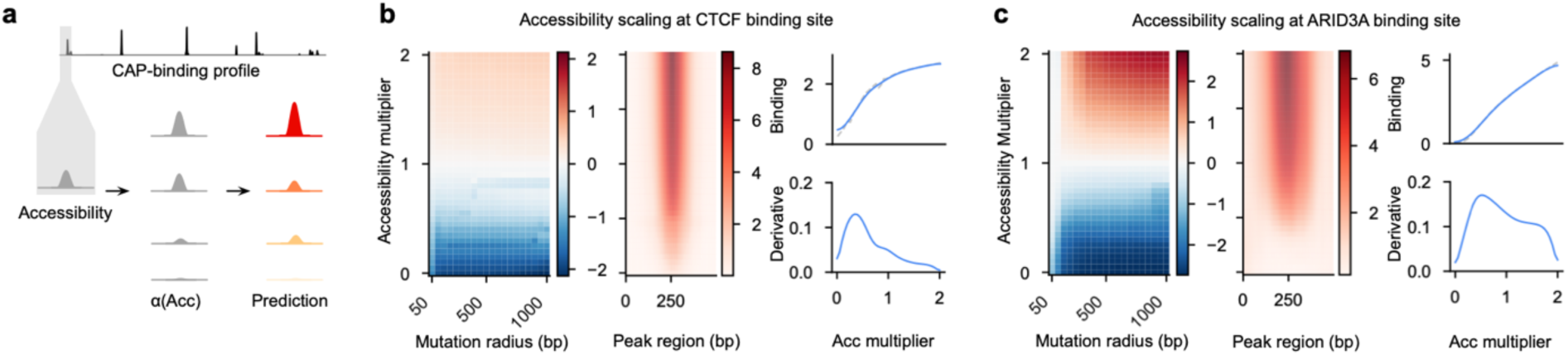
*In silico* chromatin accessibility perturbation revealed non-linear relationship between accessibility and CAP binding. **a.** Chromatin accessibility perturbation using *in silico* scaling multipliers. **b-c.** *In silico* chromatin accessibility perturbation results at CTCF (**b**) and ARID3A (**c**) binding sites. Left: predicted CAP binding changes responding to perturbation radius and accessibility multiplier. Middle: heatmap of predicted binding intensity upon perturbing accessibility at fixed 500 bp window, centered at binding peaks. Right: CAP binding changes (top) and the rate of change (bottom) as a function of accessibility multiplier.

**Supplementary Fig. 18:**
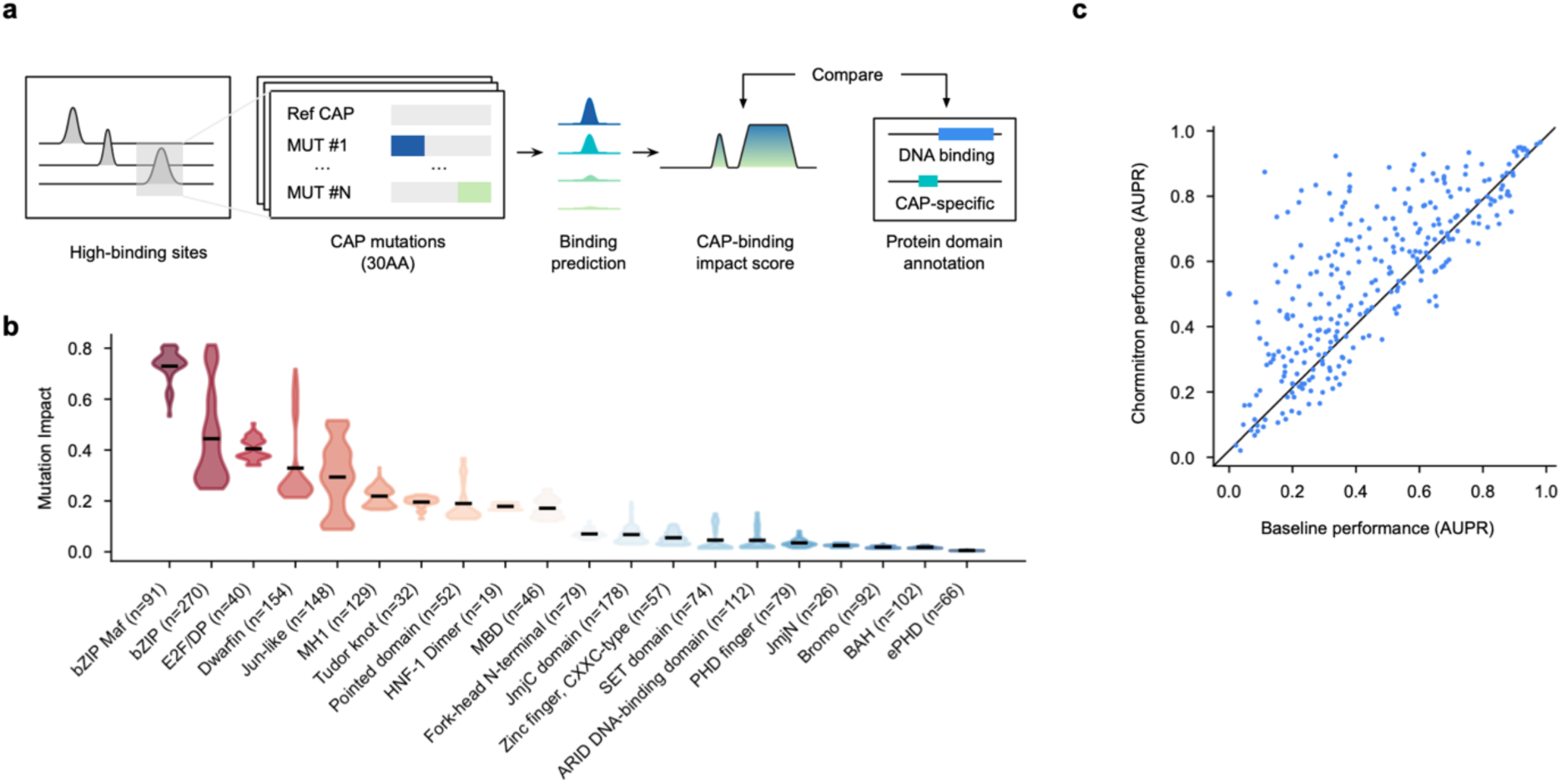
Chromnitron-based *in silico* CAP residue mutagenesis captures protein functional domains. **a.** Schematic of *in silico* mutation for CAP protein sequences. Upon mutation, predicted CAP binding intensities of mutant CAPs were compared to reference CAP to calculate protein residue impact scores (Methods). **b.** Violin plot distribution of impact scores across CAP residues, stratified by domain annotations. Top and bottom 10 domains are shown. **c.** *In silico* CAP residue mutagenesis by Chromnitron captures protein residues from annotated functional domains. Each dot in the scatter plot represents a CAP’s AUPR scores against protein domain annotations using *in silico* mutagenesis-identified residues (y-axis) and compared with that using randomly selected residues (x-axis).

**Supplementary Fig. 19:**
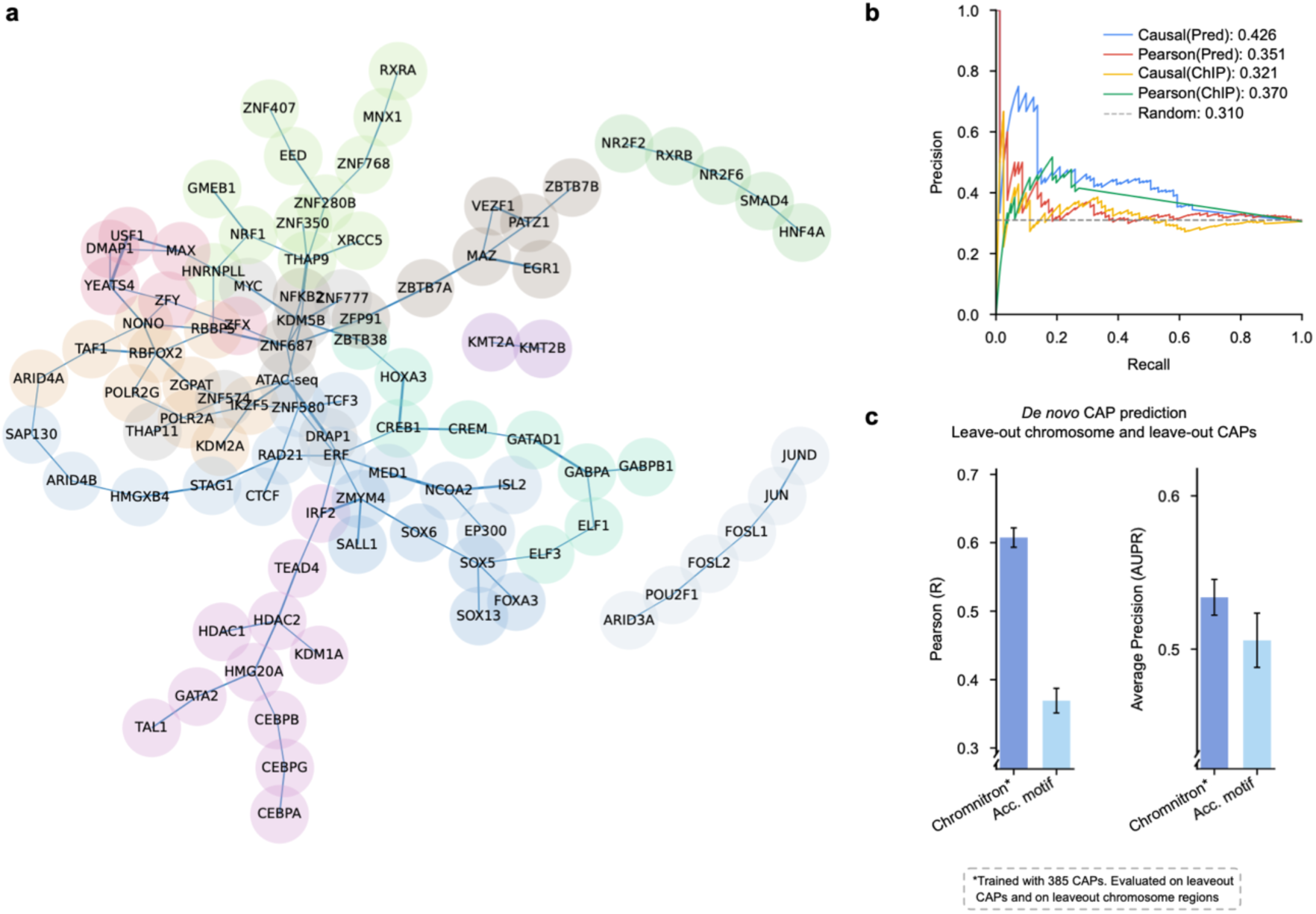
Chromnitron enables inference of protein–protein interactions and *de novo* binding prediction across proteins. **a.** Visualization of LiNGAM causal-discovery algorithm using predicted CAP binding profiles. Edges represent high interaction potential between adjacent CAPs. Nodes were colored by CAP community clusters. **b.** Performance comparison between different methods for discovering protein-protein interactions. Causal-inference-based and Pearson’s correlation-based interaction graphs derived from Chromnitron predictions and from ChIP-seq data were compared against curated interactions from the STRING database using AUPR (Methods). **c.** Evaluation of *de novo* CAP-binding prediction for unseen proteins. Performance evaluation of prototype Chromnitron pretrained on 385 CAPs (in HepG2 and K562 cells) and evaluated in leave-out CAPs and leave-out genomic region (chromosome 10). Corresponding CAP protein sequence was used for Chromnitron to generalize to new CAPs. Performance was compared with accessibility-weighted motif score (Acc. Motif) baseline, demonstrating the viability of CAP-sequence-encoded structural features that enables generalizable CAP-binding prediction.

**Supplementary Fig. 20:**
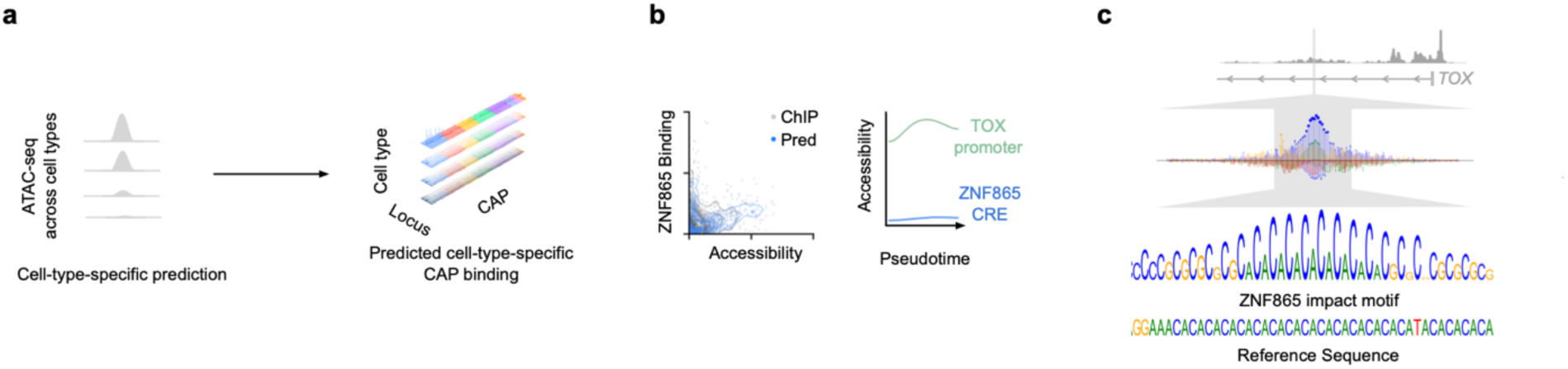
Profiling ZNF865 binding at *TOX* promoter-proximal locus. **a.** Predicting cell-type-specific CAP binding using ATAC-seq datasets across conditions. **b.** Characterization of ZNF865 binding profile at the *TOX* promoter-proximal regulatory element. Left: Scatter plot of ZNF865 experimental ChIP-seq binding (grey) and predicted binding (blue) versus chromatin accessibility (x-axis), indicating that ZNF865 preferentially binds at low-accessibility regions. Right: Chromatin accessibility signals at *TOX* promoter and the ZNF865-bound CRE across T cell exhaustion pseudo-time. **c.** Chromatin accessibility at the ZNF865-bound CRE (top) and the corresponding ZNF865 impact motif revealed by *in silico* mutagenesis (bottom).

**Supplementary Fig. 21:**
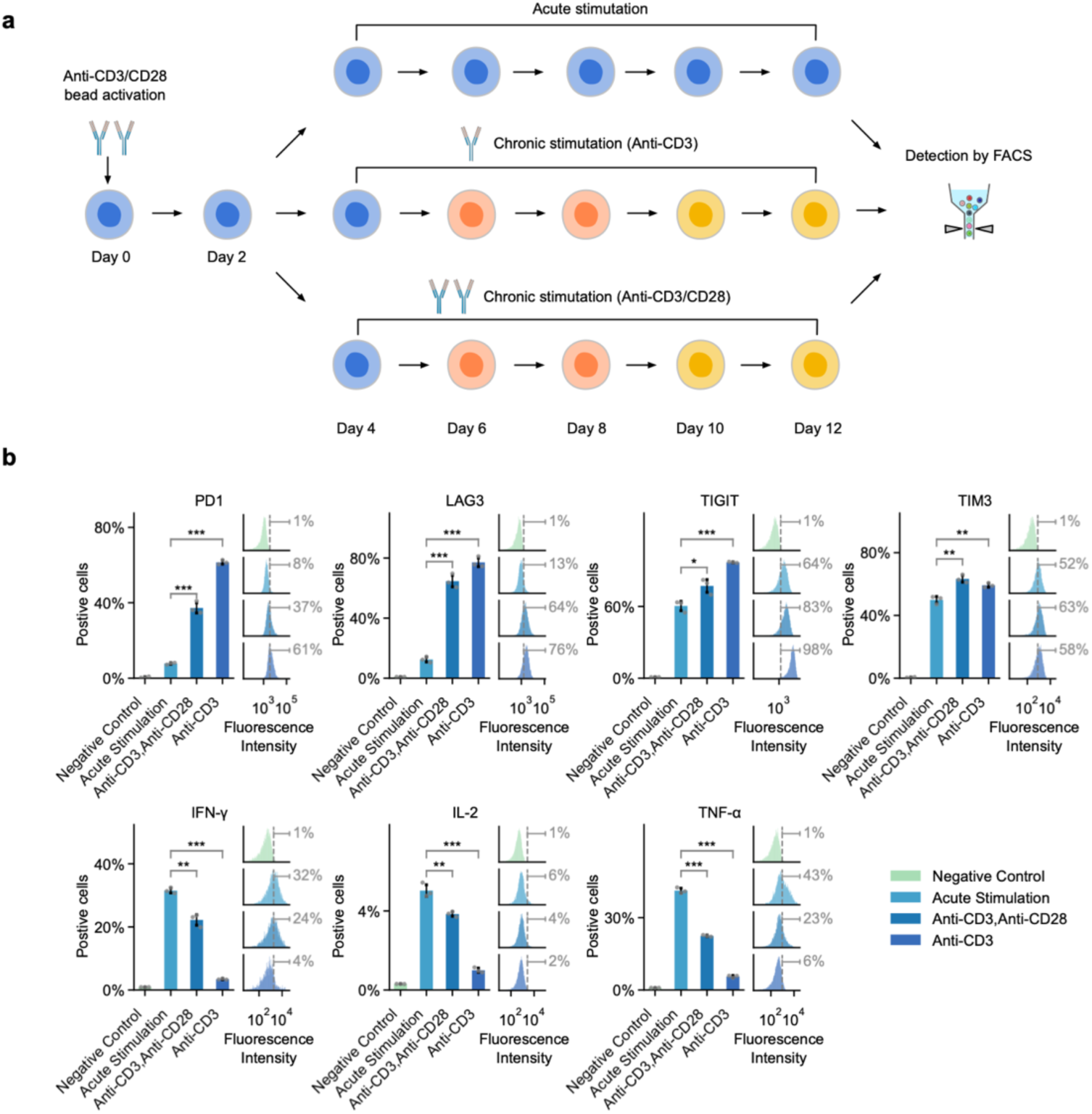
*In vitro* chronic stimulation of human primary CD8^+^ T cells to model T cell exhaustion trajectory. **a.** Schematic of the *in vitro* treatment of human CD8^+^ T cells across acute stimulation (top), chronic stimulation by anti-CD3 antibodies (middle), and chronic stimulation by anti-CD3/CD28 antibodies (bottom). **b.** Flow cytometry quantification of cell surface markers of T cell exhaustion (PD1, LAG3, TIGIT, and TIM3) and intracellular cytokine production (IFN-γ, IL-2, and TNF-α) in different stimulation groups at day 12. Bar plots show mean ± s.d. for n = 3 independent replicates. Statistical comparisons performed using two-sided unpaired Student’s *t*-tests with significance denoted as *, P < 0.05; **, P < 0.01; ***, P < 0.001.

**Supplementary Fig. 22:**
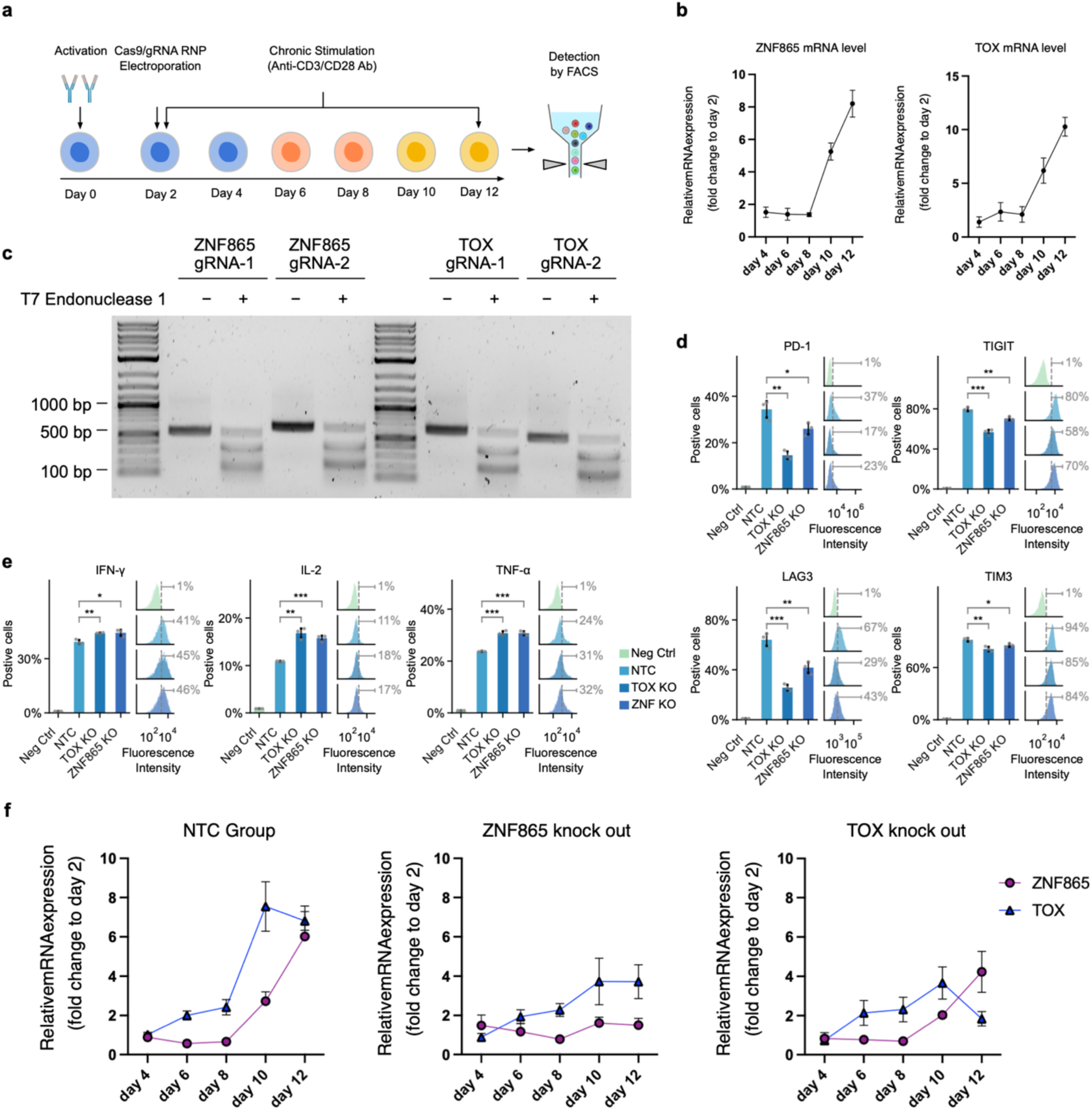
Validating ZNF865 as an upstream regulator of *TOX* for T cell exhaustion. **a.** Schematic of *in vitro* model of T cell exhaustion through persistent anti-CD3/CD28 stimulation, coupled with CRISPR/Cas9-based gene knockout in primary T cells. **b.** RT-qPCR quantification of *TOX* and *ZNF865* mRNA expression level changes across chronic stimulation of human CD8^+^ T cells from day 4 to day 12, indicating increased expression along the trajectory. Relative expression levels were normalized to day 2 using the ΔΔCt method. Each dot represents an independent biological replicate (n = 4); error bars indicate mean ± s.e.m. **c.** Surveyor assay confirmed efficient CRISPR knockout gRNA targeting *TOX* and *ZNF865*. Each gene includes two gRNAs for knock-out perturbation. Genomic DNA was amplified around the gRNA target sites and the fragments were digested with T7 Endonuclease I. **d-e.** Flow cytometry quantification of T cell exhaustion markers (**e**, PD1, TIGIT, LAG3 and TIM3) and cytokines (**f,** IFN-γ, IL-2, and TNF-α) across negative control, non-targeting control (NTC), *TOX* knockout, or *ZNF865* knockout conditions. Bar plots show mean ± s.d. for n = 3 independent replicates. Statistical comparisons performed using two-sided unpaired Student’s *t*-tests with significance denoted as *, P < 0.05; **, P < 0.01; ***, P < 0.001. **f.** RT–qPCR quantification of *ZNF865* and *TOX* mRNA level changes across chronic stimulation days with individual gene knockouts (*ZNF865* KO and *TOX* KO). Relative expression levels were normalized to day 2. Each dot represents an independent biological replicate (n = 4); error bars indicate mean ± s.e.m.

**Supplementary Fig. 23:**
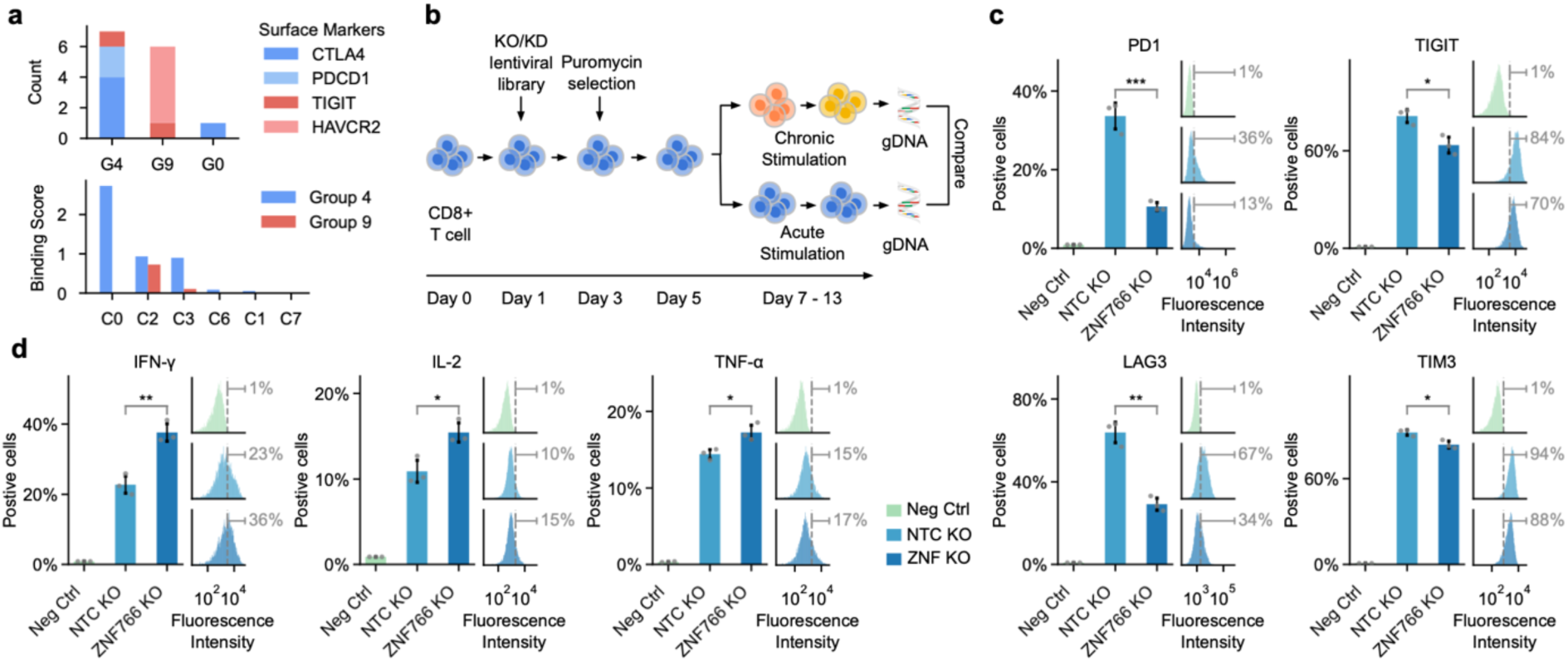
Validation of CRISPR-mediated *ZNF766* knock-out effects on chronic-stimulated T cells. **a.** Distribution of reported exhausted T cell (Tex) marker genes (CTLA4, PDCD1, TIGIT, HAVCR2) in co-regulated CRE groups (top) and their corresponding co-regulating CAP clusters (bottom). **b.** Schematic of CRISPR dropout screens in human CD8⁺ T cells using *in vitro* chronic stimulation model. **c-d.** Flow cytometry quantification of T cell exhaustion markers (**c**, PD1, TIGIT, LAG3 and TIM3) and cytokine secretion levels (**d,** IFN-γ, IL-2, and TNF-α) across negative control, non-targeting control (NTC), or *ZNF766* knockout conditions. Bar plots show mean ± s.d. for n = 3 independent replicates. Statistical comparisons performed using two-sided unpaired Student’s *t*-tests with significance denoted as *, P < 0.05; **, P < 0.01; ***, P < 0.001.

**Supplementary Fig. 24:**
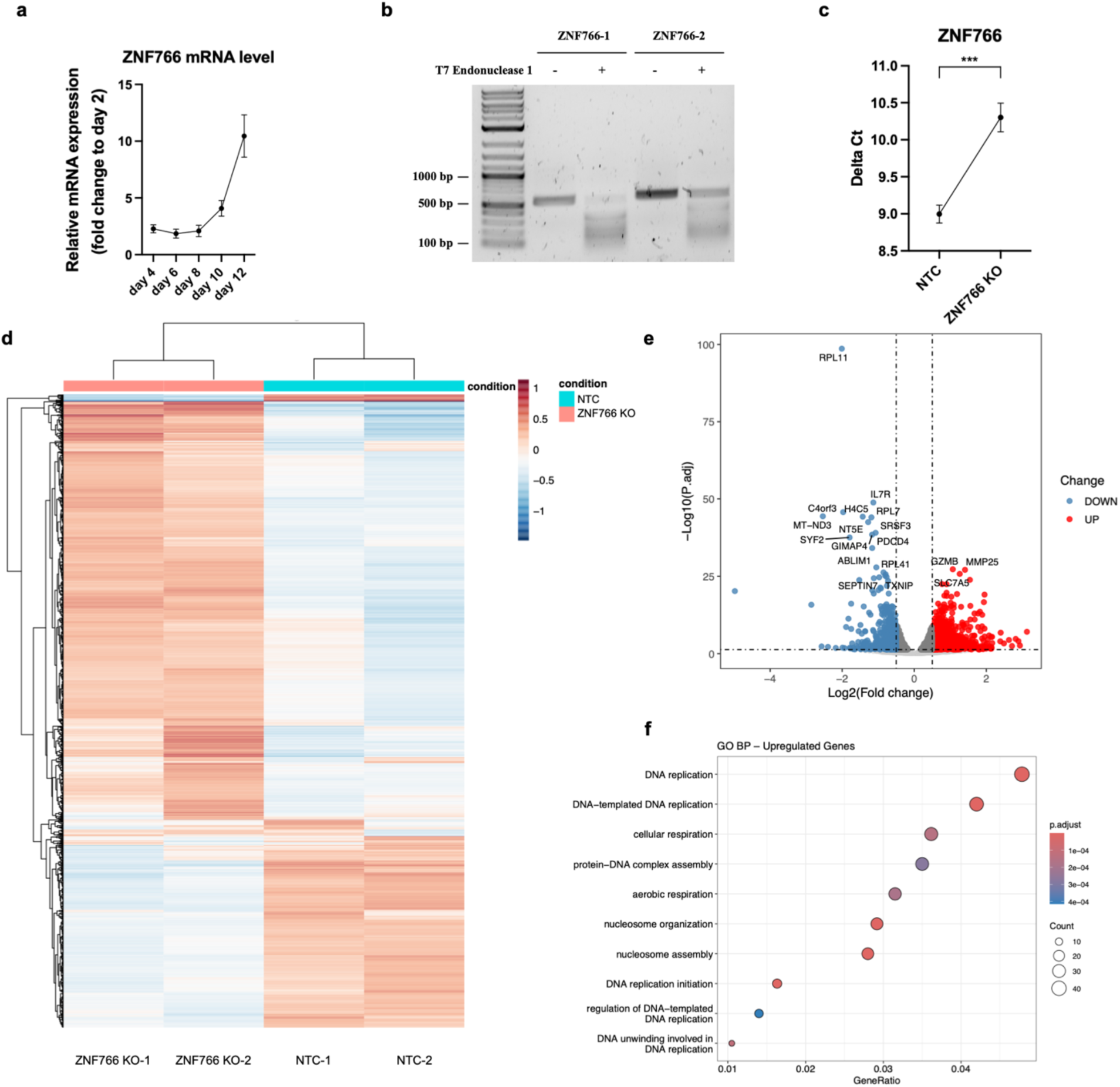
Transcriptomic profiling of *ZNF766* knock-out effects in chronic-stimulated T cells. **a.** RT-qPCR quantification of *ZNF766* mRNA level changes across chronic stimulation of CD8^+^ T cells from day 4 to day 12, indicating increased expression along the trajectory. Relative expression levels were normalized to day 2 using the ΔΔCt method. Each dot represents an independent biological replicate (n = 4); error bars indicate mean ± s.e.m. **b.** Surveyor assay confirmed efficient CRISPR knockout gRNA targeting *ZNF766*. **c.** RT-qPCR quantification of *ZNF766* mRNA expression levels after gene knockout. Primers were designed flanking the CRISPR cutting site to confirm transcript disruption. Increased ΔCt number after perturbation indicates efficient knock down of unperturbed *ZNF766* mRNA. Error bars indicate mean ± s.e.m. Statistical significance was determined using a two-tailed paired Student’s *t*-test (P < 0.001). **d.** Heatmap of differentially expressed genes between *ZNF766* knockout and non-targeting control (NTC) T cells. **e.** Volcano plot of differentially expressed genes between *ZNF766* knockout and NTC groups. Red and blue dots indicate significantly upregulated and downregulated genes, respectively, in the knock-out condition (|log₂FC| > 1, p.adj < 0.05). Representative genes involved in T cell activation (e.g., *GZMB*, *MMP25*, *SLC7A5*) and exhaustion-associated regulators (e.g., *PDCD4*, *NT5E*, *TXNIP*) were individually labeled. **f.** Gene Ontology (Biological Process) enrichment analysis of significantly upregulated genes in *ZNF766* knockout group. Dot size corresponds to gene count per term; color indicates adjusted p-value. Enriched pathways highlight enhanced DNA replication, chromatin organization, and mitochondrial respiration, suggesting a functional restoration of exhausted T cells.

**Supplementary Fig. 25:**
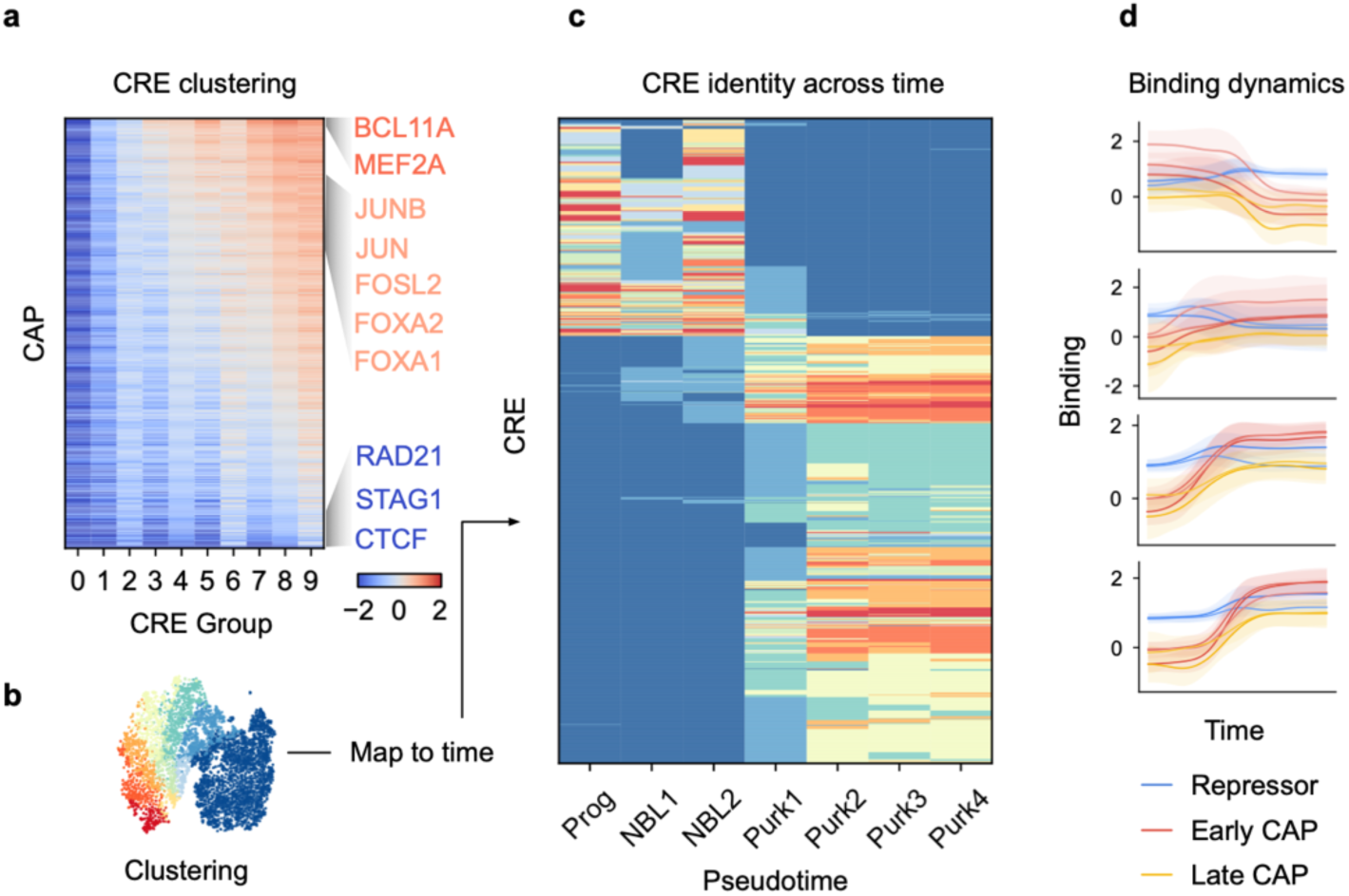
Dynamics of CAP binding and CRE activities during Purkinje cell differentiation. **a.** CAP binding profiles across CRE groups. Representative CAPs were labeled. Color scale indicates normalized binding. **b.** T-SNE embedding of CREs colored by activity clusters (same as Fig. 4D). **c.** Heatmap of CRE identity transitions across Purkinje cell differentiation stages. **d.** Average binding dynamics for four different CRE sets across pseudo-time. Each CRE set includes three CAP classes (Repressor: e.g. TCF7, RCOR1; Early CAP: e.g. BCL11A, MEF2A, ATF2; Late CAP: e.g. FOXA1, FOXA2). Shading denotes 1 standard deviation.

**Supplementary Fig. 26:**
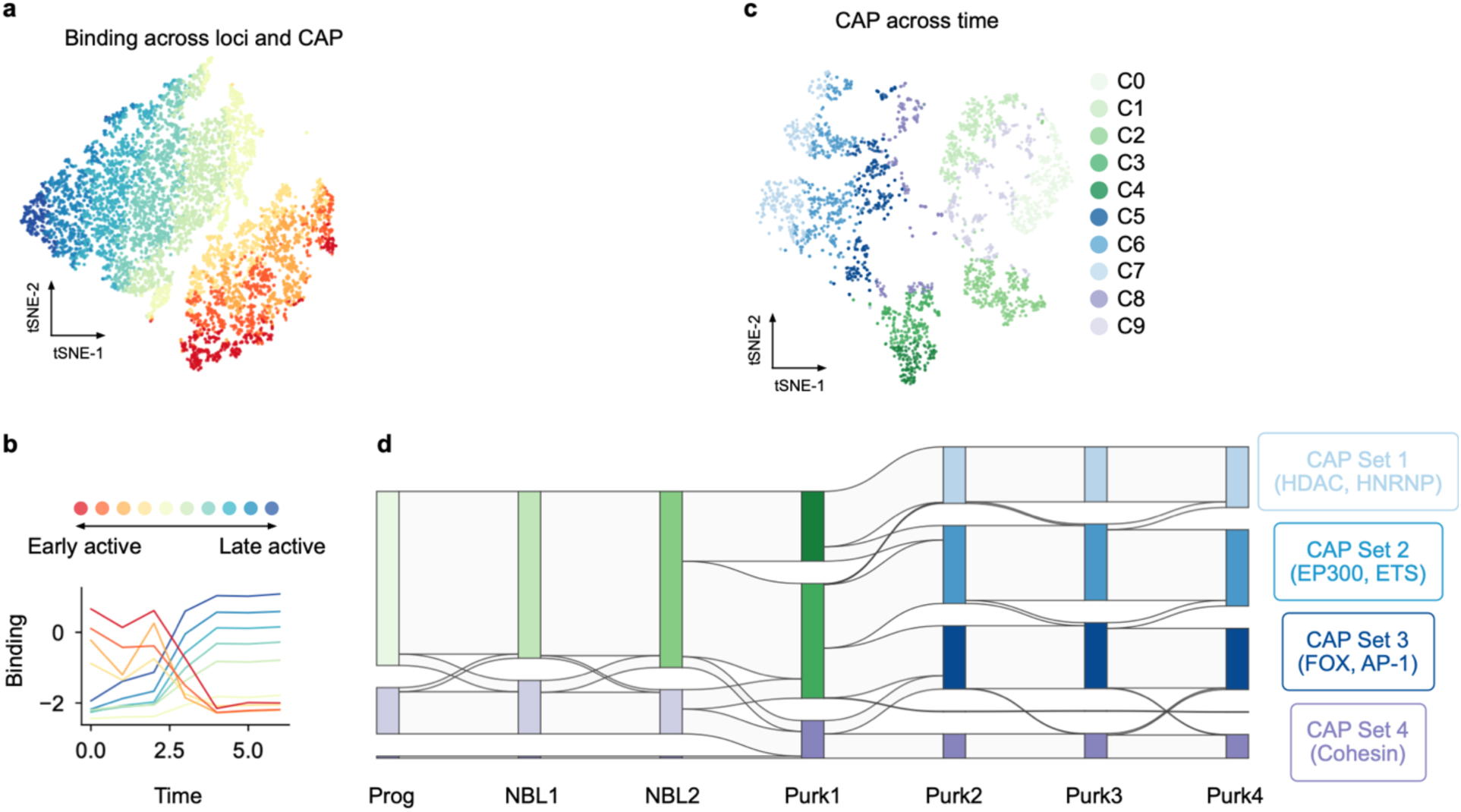
Characterizing Purkinje cell differentiation dynamics using pseudo-time and CRE as organizing features. **a.** T-SNE embedding of individualized CAP binding across loci using time trajectory as feature, colored by trajectory trend cluster. Each dot represents a CAP binding at a single locus across all pseudo-time stages. **b.** Average CAP-binding trajectories for each cluster across time, illustrating both early- and late-acting regulatory programs. **c.** T-SNE embedding of CAPs across time using CRE-level binding profiles as features, colored by CAP cluster identities. Each dot represents a CAP’s binding profile across all loci at a given time point. **d.** Alluvial diagram showing transitions in CAP cluster identity across developmental stages. Representative CAPs in each cluster were annotated on the right, highlighting coordinated shifts in regulatory factor usage during Purkinje cell maturation.

## References

1. Arendt, D. et al. The origin and evolution of cell types. Nat. Rev. Genet. 17, 744–757 (2016).

2. Xia, B. & Yanai, I. A periodic table of cell types. Dev. Camb. Engl. 146, dev169854 (2019).

3. Gurdon, J. B. Cell Fate Determination by Transcription Factors. Curr. Top. Dev. Biol. 116, 445–454 (2016).

4. Chen, X. et al. Integration of external signaling pathways with the core transcriptional network in embryonic stem cells. Cell 133, 1106–1117 (2008).

5. Boyer, L. A. et al. Core transcriptional regulatory circuitry in human embryonic stem cells. Cell 122, 947–956 (2005).

6. Gerstein, M. B. et al. Architecture of the human regulatory network derived from ENCODE data. Nature 489, 91–100 (2012).

7. Li, Y. E. et al. A comparative atlas of single-cell chromatin accessibility in the human brain. Science 382, eadf7044 (2023).

8. Lander, E. S. et al. Initial sequencing and analysis of the human genome. Nature 409, 860–921 (2001).

9. Lambert, S. A. et al. The Human Transcription Factors. Cell 172, 650–665 (2018).

10. Vaquerizas, J. M., Kummerfeld, S. K., Teichmann, S. A. & Luscombe, N. M. A census of human transcription factors: function, expression and evolution. Nat. Rev. Genet. 10, 252–263 (2009).

11. Weirauch, M. T. et al. Determination and inference of eukaryotic transcription factor sequence specificity. Cell 158, 1431–1443 (2014).

12. Jolma, A. et al. DNA-binding specificities of human transcription factors. Cell 152, 327–339 (2013).

13. Vierstra, J. et al. Global reference mapping of human transcription factor footprints. Nature 583, 729–736 (2020).

14. Murre, C. Helix-loop-helix proteins and the advent of cellular diversity: 30 years of discovery. Genes Dev. 33, 6–25 (2019).

15. Doughty, B. R. et al. Single-molecule states link transcription factor binding to gene expression. Nature 636, 745–754 (2024).

16. Durdu, S. et al. Chromatin-dependent motif syntax defines differentiation trajectories. Mol. Cell 85, 2900–2918.e16 (2025).

17. Kim, S. & Wysocka, J. Deciphering the multi-scale, quantitative cis-regulatory code. Mol. Cell 83, 373–392 (2023).

18. Kim, S. et al. DNA-guided transcription factor cooperativity shapes face and limb mesenchyme. Cell 187, 692–711.e26 (2024).

19. Statello, L., Guo, C.-J., Chen, L.-L. & Huarte, M. Gene regulation by long non-coding RNAs and its biological functions. Nat. Rev. Mol. Cell Biol. 22, 96–118 (2021).

20. Smith, E. & Shilatifard, A. The chromatin signaling pathway: diverse mechanisms of recruitment of histone-modifying enzymes and varied biological outcomes. Mol. Cell 40, 689–701 (2010).

21. Moore, J. E. et al. An expanded registry of candidate cis-regulatory elements. Nature 10.1038/s41586-025-09909-9 (2026) doi:10.1038/s41586-025-09909-9.

22. Yosef, N. et al. Dynamic regulatory network controlling TH17 cell differentiation. Nature 496, 461–468 (2013).

23. Johnson, D. S., Mortazavi, A., Myers, R. M. & Wold, B. Genome-wide mapping of in vivo protein-DNA interactions. Science 316, 1497–1502 (2007).

24. Tsankov, A. M. et al. Transcription factor binding dynamics during human ES cell differentiation. Nature 518, 344–349 (2015).

25. Park, P. J. ChIP-seq: advantages and challenges of a maturing technology. Nat. Rev. Genet. 10, 669–680 (2009).

26. Skene, P. J. & Henikoff, S. An efficient targeted nuclease strategy for high-resolution mapping of DNA binding sites. eLife 6, e21856 (2017).

27. Partridge, E. C. et al. Occupancy maps of 208 chromatin-associated proteins in one human cell type. Nature 583, 720–728 (2020).

28. Hammal, F., de Langen, P., Bergon, A., Lopez, F. & Ballester, B. ReMap 2022: a database of Human, Mouse, Drosophila and Arabidopsis regulatory regions from an integrative analysis of DNA-binding sequencing experiments. Nucleic Acids Res. 50, D316–D325 (2022).

29. Moyers, B. A. et al. Characterization of human transcription factor function and patterns of gene regulation in HepG2 cells. Genome Res. 33, 1879–1892 (2023).

30. Wang, J. et al. Sequence features and chromatin structure around the genomic regions bound by 119 human transcription factors. Genome Res. 22, 1798–1812 (2012).

31. Avsec, Ž., et al. Base-resolution models of transcription-factor binding reveal soft motif syntax. Nat. Genet. 53, 354–366 (2021).

32. Linder, J., Srivastava, D., Yuan, H., Agarwal, V. & Kelley, D. R. Predicting RNA-seq coverage from DNA sequence as a unifying model of gene regulation. Nat. Genet. 57, 949–961 (2025).

33. Avsec, Ž., et al. AlphaGenome: advancing regulatory variant effect prediction with a unified DNA sequence model.

34. Avsec, Ž., et al. Base-resolution models of transcription-factor binding reveal soft motif syntax. Nat. Genet. 53, 354–366 (2021).

35. Lin, Z. et al. Evolutionary-scale prediction of atomic-level protein structure with a language model.

36. Abramson, J. et al. Accurate structure prediction of biomolecular interactions with AlphaFold 3. Nature 630, 493–500 (2024).

37. Nguyen, E. et al. Sequence modeling and design from molecular to genome scale with Evo. Science 386, eado9336 (2024).

38. Fu, X. et al. A foundation model of transcription across human cell types. Nature 10.1038/s41586-024-08391-z (2025) doi:10.1038/s41586-024-08391-z.

39. Hayes, T. et al. Simulating 500 million years of evolution with a language model. Preprint at 10.1101/2024.07.01.600583 (2024).

40. Theodoris, C. V. et al. Transfer learning enables predictions in network biology. Nature 618, 616–624 (2023).

41. Cazares, T. A. et al. maxATAC: Genome-scale transcription-factor binding prediction from ATAC-seq with deep neural networks. PLOS Comput. Biol. 19, e1010863 (2023).

42. Avsec, Ž., et al. Effective gene expression prediction from sequence by integrating long-range interactions. Nat. Methods 18, 1196–1203 (2021).

43. Hu, E. J., et al. LoRA: Low-Rank Adaptation of Large Language Models. Preprint at 10.48550/arXiv.2106.09685 (2021).

44. Bravo González-Blas, C., et al. SCENIC+: single-cell multiomic inference of enhancers and gene regulatory networks. Nat. Methods 20, 1355–1367 (2023).

45. Hu, Y. et al. Multiscale footprints reveal the organization of cis-regulatory elements. Nature 638, 779–786 (2025).

46. Barral, A. & Zaret, K. S. Pioneer factors: roles and their regulation in development. Trends Genet. 40, 134–148 (2024).

47. Balsalobre, A. & Drouin, J. Pioneer factors as master regulators of the epigenome and cell fate. Nat. Rev. Mol. Cell Biol. 23, 449–464 (2022).

48. Stoltenburg, R., Reinemann, C. & Strehlitz, B. SELEX—A (r)evolutionary method to generate high-affinity nucleic acid ligands. Biomol. Eng. 24, 381–403 (2007).

49. Ren, B. et al. Genome-Wide Location and Function of DNA Binding Proteins. Science 290, 2306–2309 (2000).

50. Khan, O. et al. TOX transcriptionally and epigenetically programs CD8+ T cell exhaustion. Nature 571, 211–218 (2019).

51. Seo, H. et al. TOX and TOX2 transcription factors cooperate with NR4A transcription factors to impose CD8 ^+^ T cell exhaustion. Proc. Natl. Acad. Sci. 116, 12410–12415 (2019).

52. Yao, C. et al. Single-cell RNA-seq reveals TOX as a key regulator of CD8+ T cell persistence in chronic infection. Nat. Immunol. 20, 890–901 (2019).

53. Mann, T. H. & Kaech, S. M. Tick-TOX, it’s time for T cell exhaustion. Nat. Immunol. 20, 1092–1094 (2019).

54. Scott, A. C. et al. TOX is a critical regulator of tumour-specific T cell differentiation. Nature 571, 270–274 (2019).

55. Alfei, F. et al. TOX reinforces the phenotype and longevity of exhausted T cells in chronic viral infection. Nature 571, 265–269 (2019).

56. Hashimoto, M. et al. CD8 T Cell Exhaustion in Chronic Infection and Cancer: Opportunities for Interventions. Annu. Rev. Med. 69, 301–318 (2018).

57. Baessler, A. & Vignali, D. A. A. T Cell Exhaustion. Annu. Rev. Immunol. 42, 179–206 (2024).

58. Satpathy, A. T. et al. Massively parallel single-cell chromatin landscapes of human immune cell development and intratumoral T cell exhaustion. Nat. Biotechnol. 37, 925–936 (2019).

59. Jenkins, E., Whitehead, T., Fellermeyer, M., Davis, S. J. & Sharma, S. The current state and future of T cell exhaustion research. Oxf. Open Immunol. 4, iqad006 (2023).

60. Tay, T. et al. Degradation of IKZF1 prevents epigenetic progression of T cell exhaustion in an antigen-specific assay. Cell Rep. Med. 5, 101804 (2024).

61. Belk, J. A. et al. Genome-wide CRISPR screens of T cell exhaustion identify chromatin remodeling factors that limit T cell persistence. Cancer Cell 40, 768–786.e7 (2022).

62. Mannens, C. C. A. et al. Chromatin accessibility during human first-trimester neurodevelopment. Nature 10.1038/s41586-024-07234-1 (2024) doi:10.1038/s41586-024-07234-1.

63. Pampari, A. et al. ChromBPNet: bias factorized, base-resolution deep learning models of chromatin accessibility reveal cis-regulatory sequence syntax, transcription factor footprints and regulatory variants. Preprint at 10.1101/2024.12.25.630221 (2024).

64. Chen, K. M., Wong, A. K., Troyanskaya, O. G. & Zhou, J. A sequence-based global map of regulatory activity for deciphering human genetics. Nat. Genet. 54, 940–949 (2022).

65. Hendrycks, D. & Gimpel, K. Gaussian Error Linear Units (GELUs). Preprint at 10.48550/arXiv.1606.08415 (2023).

66. Vaswani, A., et al. Attention Is All You Need. ArXiv170603762 Cs http://arxiv.org/abs/1706.03762 (2017).

67. Meng, Q. et al. A generic reference defined by consensus peaks for scATAC-seq data analysis. Preprint at 10.1101/2023.05.30.542889 (2023).

68. Amemiya, H. M., Kundaje, A. & Boyle, A. P. The ENCODE Blacklist: Identification of Problematic Regions of the Genome. Sci. Rep. 9, 9354 (2019).

69. Kingma, D. P. & Ba, J. Adam: A Method for Stochastic Optimization. Preprint at 10.48550/arXiv.1412.6980 (2017).

70. Hu, E. J., et al. LoRA: Low-Rank Adaptation of Large Language Models. Preprint at 10.48550/arXiv.2106.09685 (2021).

71. Shrikumar, A., et al. Technical Note on Transcription Factor Motif Discovery from Importance Scores (TF-MoDISco) version 0.5.6.5. Preprint at 10.48550/arXiv.1811.00416 (2020).

72. Blum, M. et al. InterPro: the protein sequence classification resource in 2025. Nucleic Acids Res. 53, D444–D456 (2025).

73. Ikeuchi, T., Ide, M., Zeng, Y., Maeda, T. N. & Shimizu, S. Python package for causal discovery based on LiNGAM.

74. Szklarczyk, D. et al. The STRING database in 2023: protein–protein association networks and functional enrichment analyses for any sequenced genome of interest. Nucleic Acids Res. 51, D638–D646 (2023).

75. Wolf, F. A., Angerer, P. & Theis, F. J. SCANPY: large-scale single-cell gene expression data analysis. Genome Biol. 19, 15 (2018).

76. Robinson, J. T. et al. Integrative genomics viewer. Nat. Biotechnol. 29, 24–26 (2011).

77. Li, W. et al. MAGeCK enables robust identification of essential genes from genome-scale CRISPR/Cas9 knockout screens. Genome Biol. 15, 554 (2014).

